# The components of directional and disruptive selection in heterogeneous group-structured populations

**DOI:** 10.1101/2020.03.02.974279

**Authors:** Hisashi Ohtsuki, Claus Rueffler, Joe Yuichiro Wakano, Kalle Parvinen, Laurent Lehmann

## Abstract

We derive how directional and disruptive selection operate on scalar traits in a heterogeneous group-structured population for a general class of models. In particular, we assume that each group in the population can be in one of a finite number of states, where states can affect group size and/or other environmental variables, at a given time. Using up to second-order perturbation expansions of the invasion fitness of a mutant allele, we derive expressions for the directional and disruptive selection coefficients, which are sufficient to classify the singular strategies of adaptive dynamics. These expressions include first- and second-order perturbations of individual fitness (expected number of settled offspring produced by an individual, possibly including self through survival); the first-order perturbation of the stationary distribution of mutants (derived here explicitly for the first time); the first-order perturbation of pairwise relatedness; and reproductive values, pairwise and three-way relatedness, and stationary distribution of mutants, each evaluated under neutrality. We introduce the concept of individual *k*-fitness (defined as the expected number of settled offspring of an individual for which *k −* 1 randomly chosen neighbors are lineage members) and show its usefulness for calculating relatedness and its perturbation. We then demonstrate that the directional and disruptive selection coefficients can be expressed in terms individual *k*-fitnesses with *k* = 1, 2, 3 only. This representation has two important benefits. First, it allows for a significant reduction in the dimensions of the system of equations describing the mutant dynamics that needs to be solved to evaluate explicitly the two selection coefficients. Second, it leads to a biologically meaningful interpretation of their components. As an application of our methodology, we analyze directional and disruptive selection in a lottery model with either hard or soft selection and show that many previous results about selection in group-structured populations can be reproduced as special cases of our model.

## 1 Introduction

Many natural populations are both group-structured – with the number of individuals interacting at the local scale being finite – and heterogeneous – with different groups being subject to different demographic and environmental conditions (*e.g.*, varying group size and temperature, respectively). Understanding how evolution, and in particular natural selection, moulds phenotypic traits in such systems is complicated as both local heterogeneity and demographic stochasticity need to be taken into account. In order to predict the outcome of evolution in heterogeneous populations, evolutionists are generally left with the necessity to approximate the evolutionary dynamics, as a full understanding of this process is yet out of reach.

A standard approximation to predict evolutionary outcomes is to assume that traits are quantitative, that the details of inheritance do not matter (“phenotypic gambit”, Grafen, 1991), and that mutations have weak (small) phenotypic effects (*e.g.* Grafen, 1985; Taylor, 1989; Parker and Maynard Smith, 1990; Rousset, 2004). Under these assumptions, directional trait evolution can be quantified by a phenotypic selection gradient that captures first-order effects of selection. Thus, phenotypic change occurs in an up-hill direction on the fitness landscape. This directional selection either causes the trait value to change endlessly (for instance, due to macro environmental changes or cycles in the evolutionary dynamics), or the trait value eventually approaches a local equilibrium point, a so-called *singular strategy*, where directional selection vanishes. Such a singular strategy may be locally uninvadable (“evolutionary stable”) and thus a local end-point of the evolutionary dynamics. However, when the fitness landscape is dynamic due to selection being frequency-dependent, then it is also possible that, as the population evolves uphill on the fitness landscape, this landscape changes such that the population eventually finds itself at a singular strategy that is located in a fitness valley. In this case, directional selection turns into disruptive selection, which means that a singular strategy that is an attractor of the evolutionary dynamics (and thus convergence stable) is invadable by nearby mutants and thus an *evolutionary branching point* (Metz et al., 1996; Geritz et al., 1998). Further evolutionary dynamics can then result in genetic polymorphism in the population, thus possibly favoring the maintenance of adaptive diversity in the long term (see Rueffler et al., 2006, for a review). Disruptive selection at a singular point is quantified by the disruptive selection coefficient (called quadratic selection gradient in the older literature: Lande and Arnold, 1983; Phillips and Arnold, 1989), which involves second-order effects of selection.

A central question concerns the nature and interpretation of the components of the selection gradient and the disruptive selection coefficient on a quantitative trait in heterogeneous populations. For the selection gradient, this question has been studied for a long time and a general answer has been given under the assumption that individuals can be in a finite number of states (summarized in Rousset, 2004). Then, regardless of the complexity of the spatial, demographic, environmental, or physiological states individuals can be in or experience (in the kin-selection literature commonly referred to as class-structure, *e.g.* Taylor, 1990; Frank, 1998; Rousset, 2004), the selection gradient on a quantitative trait depends on three key components (Taylor, 1990; Frank, 1998; Rousset, 2004). The first component are individual fitness differentials, which capture the marginal gains and losses of producing offspring in particular states to parents in particular states. The second component are (neutral) reproductive values weighting these fitness differentials. These capture the fact that offspring settling in different states contribute differently to the future gene pool. The third component are (neutral) relatedness coefficients. These also weight the fitness differentials, and capture the fact that some pairs of individuals are more likely to carry the same phenotype (inherited from a common ancestor) than randomly sampled individuals. This results in correlations between the trait values of interacting individuals. Such correlations matter for selection (“kin selection”, *e.g.* Michod, 1982) and occur in populations subject to limited genetic mixing and small local interaction groups. At the risk of oversimplifying, reproductive values can be thought of as capturing the effect of population heterogeneity on directional selection, while relatedness captures the effect of demographic stochasticity under limited genetic mixing.

The situation is different with respect to the coefficient of disruptive selection, *i.e.*, the second-order effects of selection. The components of the disruptive selection coefficient have not been worked out in general and are studied only under the assumptions of well-mixed or spatially structured populations, but with otherwise homogeneous individuals. For the spatially structured case the effects of selection on relatedness has been shown to matter, as selection changes the number of individuals expressing similar trait values in a certain group (Ajar, 2003; Wakano and Lehmann, 2014; Mullon et al., 2016), resulting in a reduced strength of disruptive selection under limited dispersal. For the general case that individuals can be in different states one expects intuitively that disruptive selection also depends on how selection affects the distribution of individuals over the different states. But this has not been analyzed so far even though it is captured implicitly when second-order derivatives of invasion fitness are computed as has been done in several previous works investigating evolutionary branching in some specific models of class-structured populations (*e.g.* Massol et al., 2011; Rueffler et al., 2013; Massol and Débarre, 2015; Kisdi, 2016; Parvinen et al., 2018, 2020).

In the present paper, we develop an evolutionary model for a heterogeneous group-structured population that covers a large class of biological scenarios. For this model, we show that the disruptive selection coefficient can be expressed in terms of individual fitness differentials weighted by the neutral quantities appearing in the selection gradient. This both significantly facilitates concrete calculations under complex scenarios and allows for a biological interpretation of selection. Our results contain several previous models as special cases.

The remainder of this paper is organized as follows. (1) We start by describing a demographic model for a heterogeneous group-structured population and present some background material underlying the characterization of uninvadable (“evolutionary stable”) strategies by way of invasion fitness for this model. We here also introduce a novel individual fitness concept – individual *k*-fitness – defined as the expected number of settled offspring of an individual for which *k −* 1 randomly chosen neighbors are relatives (*i.e.*, members of the same lineage). This fitness concept plays a central role in our analysis. (2) Assuming quantitative scalar traits, we present first- and second-order perturbations of invasion fitness (*i.e.*, the selection gradient and disruptive selection coefficient, respectively), discuss their components and the interpretations thereof, and finally express all quantities in terms of individual *k*-fitness with *k* = 1, 2, 3. (3) We present a generic lottery model under spatial heterogeneity for both soft and hard selection regimes and show that the selection gradient and the disruptive selection coefficient can be computed explicitly under any scenario falling into this class of models. We then apply these results to a concrete local adaptation scenario where we derive conditions for evolutionary stability and convergence stability of singular trait values, and show their dependence on migration rate and group size. In doing so, we recover and extend previous results from the literature and show how our model connects seemingly different approaches.

## 2 Model

### 2.1 Biological assumptions

We consider a population of haploid individuals that is subdivided into infinitely many groups that are connected to each other by dispersal (*i.e.*, the infinite island model). Dispersal between groups may occur by individuals alone or by groups of individuals as in propagule dispersal, but is always random with respect to the destination group. We consider a discrete-time reproductive process and thus discrete census steps. At each census, each group is in a state 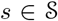 with 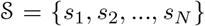 where *N* denotes the number of possible states. The state *s* determines the number of individuals in a group and/or any environmental factor determining the survival, reproduction, and dispersal of all individuals within a group. For the sake of simplicity, we will consider only a finite number of discrete states in this paper. The state *s* does not need to be a fixed property of a group but can change in time and be affected by individual trait values and thus be determined endogenously. However, we assume that such state changes are governed by a time-homogeneous Markov chain, meaning that there are no temporal trends in dynamics of the group states. We denote by *n_s_* the finite number of adult individuals in a group in state *s*, which can thus change over time if the group state changes. We assume that group size is bounded as a result of density dependence acting at the local scale (hence there is an upper bound on group size). The described set-up includes a variety of classical models.

1. Purely spatially structured populations: The state *s* is identical for all groups (*N* = 1) and so there is only one group size. This is essentially the island model as developed by Wright (1943), which has been a long-term work horse for understanding the effect of spatial structure on evolutionary dynamics (*e.g.* Eshel, 1972; Bulmer, 1986; Rousset, 2004).
2. Stochastic population dynamics at the group level: The state *s* determines the number of individuals in a group, which can potentially vary in time (*e.g.* Metz and Gyllenberg, 2001; Rousset and Ronce, 2004). This case covers the situation in which each group is embedded in a community consisting of several interacting species and where the state *s* determines the number of individuals for each of the other species (*e.g.* Chesson, 1981).
3. Environmental heterogeneity: The state *s* determines an aspect of the within-group environment, which affects the survival and/or reproduction of its group members. An example is heterogeneity in patch quality or size (*e.g.* Wild et al., 2009; Massol et al., 2011; Rodrigues and Gardner, 2012). We note that in the limit of infinite group size this coincides with models of temporal and spatial heterogeneity as reviewed in Svardal et al. (2015).
4. Group splitting: This is a special case in which migration between groups is in fact absent but groups can be connected to each other if they originate through splitting of a parental group. The state *s* again determines the number of adults in a group. This model is inspired by compartmentalized replication in prebiotic evolution (stochastic corrector model, Szathmary and Demeter, 1987; Grey et al., 1995).
5. Purely physiologically structured population: In the special case with only a single individual in a group, the state *s* can be taken to represent the physiological state of an individual such as age or size or combinations thereof (*e.g.* Ronce and Promislow, 2010). In the special case of complete and independent offspring dispersal (*i.e.*, no group dispersal) but arbitrary group size, the state *s* can be taken to represent the combination of individual physiological states of all members in a group so that the model covers within group heterogeneity.

Since we are mainly interested in natural selection driven by recurrent invasions by possibly different mutants, we can focus on the initial invasion of a mutant allele into a monomorphic resident population. Hence, we assume that at any time at most two alleles segregate in the population, a mutant allele whose carriers express the trait value *x* and a resident allele whose carriers express the trait value *y*. We furthermore assume that traits are one-dimensional and real-valued (*x, y ∈* ℝ). Suppose that initially the population is monomorphic (*i.e.*, fixed) for the resident allele *y* and a single individual mutates to trait value *x*. How do we ascertain the extinction or spread of the mutant?

### 2.2 Multitype branching process and invasion fitness

Since any mutant is initially rare, we can focus on the initial invasion of the mutant into the total population and approximate its dynamics as a discrete-time multitype branching process (Harris, 1963; Karlin and Taylor, 1975; Wild, 2011). In doing so, we largely follow the model construction and notation used in Lehmann et al. (2016) (see section A in the Supplementary Material for a mathematical description of the stochastic process underlying our model). In particular, in order to ascertain uninvadability of mutants into a population of residents it is sufficient to focus on the transition matrix ****A**** = {*a*(*s′, i′|s, i*)} whose entry in position (*s′, i′*; *s, i*), denoted by *a*(*s′, i′|s, i*), is the expected number of groups in state *s′* with *i′ ≥* 1 mutant individuals that descend from a group in state *s* with *i ≥* 1 mutant individuals over one time step in a population that is otherwise monomorphic for *y*. In the following, we refer to a group in state *s* with *i* mutants and *n_s_ − i* residents as an (*s, i*)-group for short. The transition matrix ****A**** is a square matrix that is assumed to be primitive (we note that primitivity will obtain under all models listed in section 2.1 but may be induced for different reasons). Thus, a positive integer *ℓ* (possibly depending on *x* and *y*) exists such that every entry of ***A****^ℓ^* (*ℓ*th power of ****A****) is positive. The entries *a*(*s′, i′|s, i*) of the matrix ****A**** generally depend on both *x* and *y*, but for ease of exposition we do not write these arguments explicitly unless necessary. The same convention applies to all other variables that can in principle depend on *x* and *y*.

From standard results on multitype branching processes (Harris, 1963; Karlin and Taylor, 1975) it follows that a mutant *x* arising as a single copy in an arbitrary group of the population, *i.e.*, in any (*s,* 1)-group, goes extinct with probability one if and only if the largest eigenvalue of ****A****, denoted by *ρ*, is less than or equal to one,

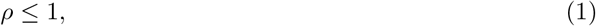

where *ρ* satisfies

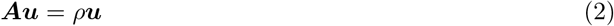

and where ****u**** is the leading right eigenvector of ****A****. We refer to *ρ* as the *invasion fitness* of the mutant. If eq.(1) holds, then we say that *y* is *uninvadable* by *x*. To better understand what determines invasion fitness, we introduce the concept of the *mutant lineage*, which we define as the collection of descendants of the initial mutant: its direct descendants (possibly including self through survival), the descendants of its immediate descendants, and so on. Invasion fitness then gives the expected number of mutant copies produced over one time step by a randomly sampled mutant from its lineage in an otherwise monomorphic resident population that has reached demographic stationarity (Mullon et al., 2016; Lehmann et al., 2016). The mutant stationary distribution is given by the vector ****u**** with entries *u*(*s, i*) describing, after normalization, the asymptotic probability that a randomly sampled group containing at least one mutant is in state *s* and contains *i ≥* 1 mutants. In other words, invasion fitness is the expected number of mutant copies produced by a lineage member randomly sampled from the distribution ****u**** (see eq.(C8) in the Supplementary Material and the explanation thereafter).

### 2.3 Statistical description of the mutant lineage

We use the matrix ****A**** = {*a*(*s′, i′|s, i*)} and its leading right eigenvector ****u**** to derive several quantities allowing us to obtain an explicit representation of invasion fitness, which will be the core of our sensitivity analysis.

#### 2.3.1 Asymptotic probabilities and relatedness of *k*-individuals

We start by noting that the asymptotic probability for a mutant to find itself in an (*s, i*)-group is given by

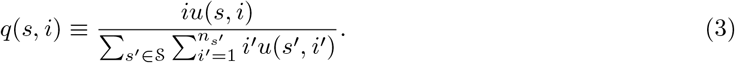

From this, we can compute two state probabilities. First, the asymptotic probability that a randomly sampled mutant finds itself in a group in state *s* is given by

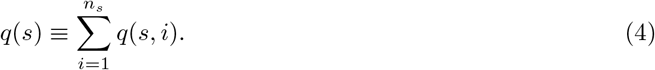

Second, the asymptotic probability that, conditional on being sampled in a group in state *s*, a randomly sampled mutant finds itself in a group with *i* mutants is given by

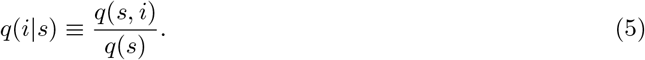

Let us further define

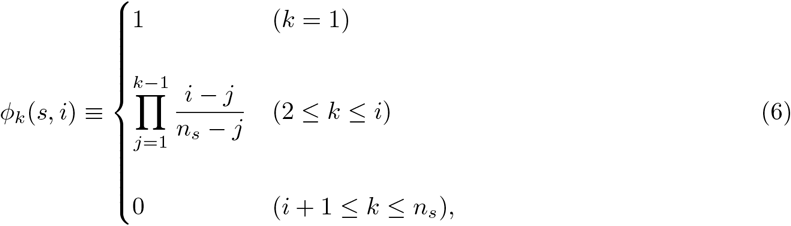

which, for *k >* 1, can be interpreted as the probability that, given a mutant is sampled from an (*s, i*)- group, *k −* 1 randomly sampled group neighbors without replacement are all mutants. This allows us to define the relatedness between *k* individuals in a group in state *s* as

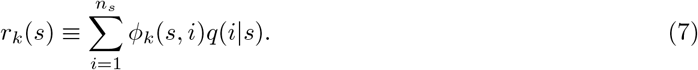

This is the probability that *k−*1 randomly sampled neighbors without replacement of a randomly sampled mutant in state *s* are also mutants (*i.e.*, they all descend from the lineage founder). For example,

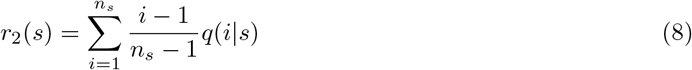

is the asymptotic probability of sampling a mutant among the neighbors of a random mutant individual from a group in state *s* and thus provides a measure of pairwise relatedness among group members. Likewise,

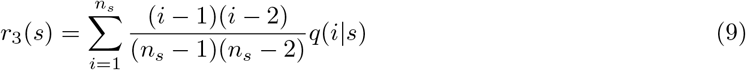

is the asymptotic probability that, conditional on being sampled in a group in state *s*, two random neighbors of a random mutant individual are also mutants.

#### 2.3.2 Individual fitness and individual *k*-fitness

Consider a mutant in an (*s, i*)-group and define

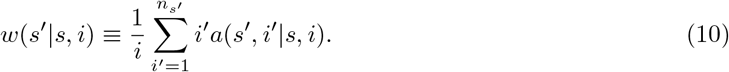

The sum on the right-hand side of eq.(10) counts the expected total number of mutants in groups in state *s′* produced by an (*s, i*)-group, and the share from a single mutant in this (*s, i*)-group is calculated by dividing this lineage productivity by *i*. Hence, *w*(*s′|s, i*) is the expected number of offspring of a mutant individual (possibly including self through survival), which settle in a group in state *s′*, given that the mutant resided in a group in state (*s, i*) in the previous time period. Thus *w*(*s′|s, i*) is an individual fitness^1^.

We now extend the concept of individual fitness to consider a collection of offspring descending from a mutant individual. More formally, for any integer *k* (1 *≤ k ≤ n_s′_*) we let

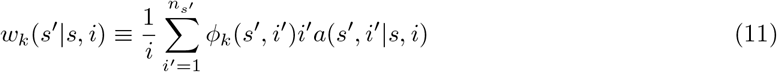

be the expected number of offspring produced by a single mutant individual in an (*s, i*)-group (possibly including self through survival) that settle in a group in state *s′* and have *k −* 1 randomly sampled group neighbors (without replacement) that are also mutants. We refer to *w_k_*(*s′|s, i*) as “individual *k*-fitness” regardless of the states *s′* and (*s, i*) (see Figure 1 for an illustrative example).

Note that individual 1-fitness equals *w*(*s′|s, i*) as defined in eq.(10). Hence, individual *k*-fitness *w_k_*(*s′|s, i*) is a generalization of this fitness concept. The difference between eq.(10) and eq.(11) is the term *φ_k_*(*s′, i′*), which shows that *k*-fitness counts an individual’s number of offspring (possibly including self through survival) that experience a certain identity-by-descent genetic state in their group. Under our assumption of infinitely many groups, more than one dispersing offspring can settle in the same group only with propagule dispersal. Thus, without propagule dispersal dispersing offspring do not contribute to *k*-fitness for *k >* 1.

**Figure 1:**
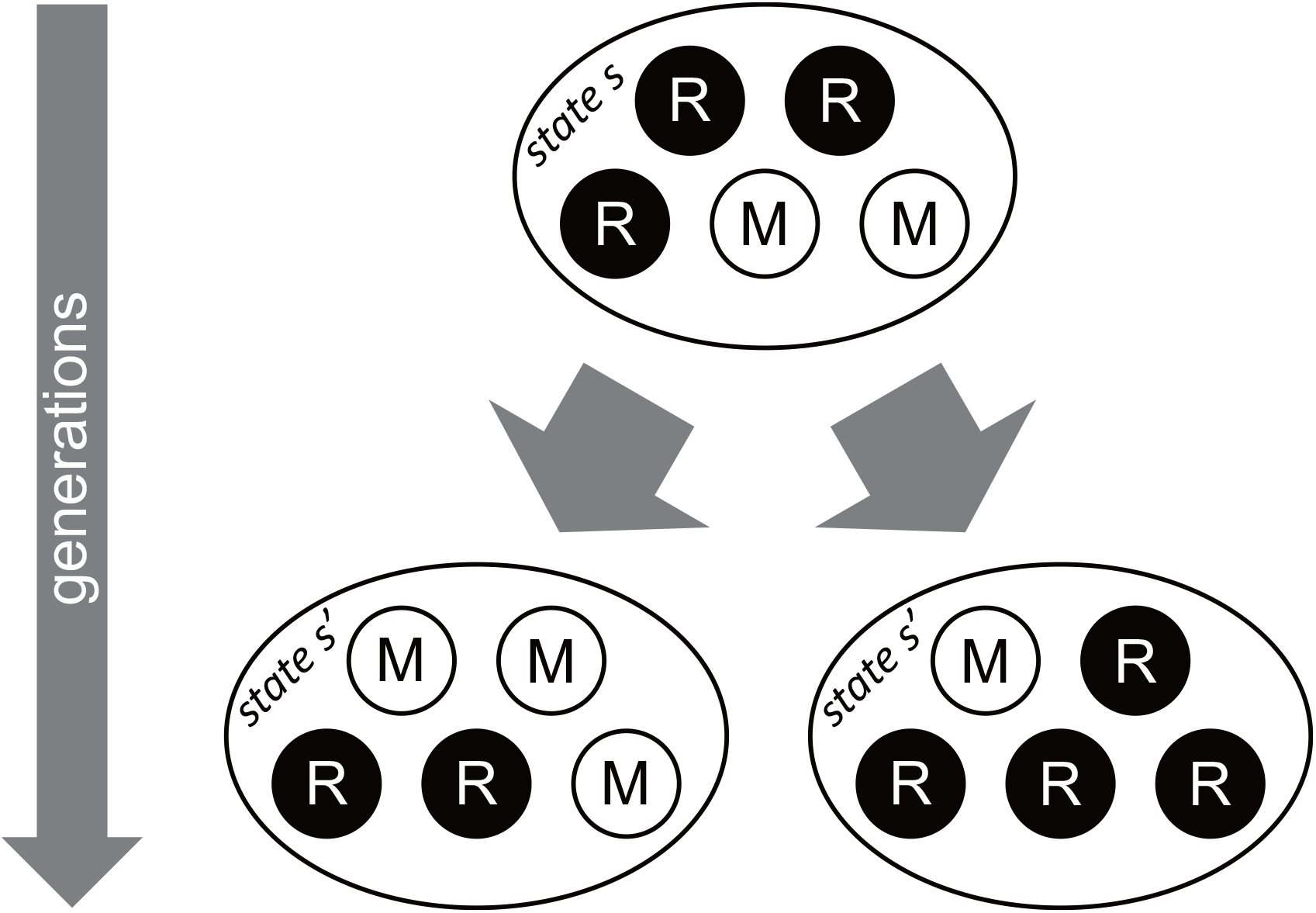
A schematic example for the calculation of individual *k*-fitness. Symbols M and R represent mutants and resident individuals, respectively. In this example, an (*s,* 2)-group “produced” one (*s′,* 3)-group and one (*s′,* 1)-group. Individual 1-fitness of each mutant in the parental generation is the total number of mutants in the following generation (3 + 1 = 4) divided by the number of mutants in the (*s,* 2)-group (= 2). Thus *w*_1_(*s′ s,* 2) = 4/2 = 2. For individual 2-fitness we calculate the weighted number of mutants in the following generation, where the weights are the probabilities that a random neighbor of a mutant is also a mutant, and then divide it by the number of mutants in the (*s,* 2)-group (= 2). These probabilities are 2/4 for the (*s′,* 3)-group and 0/4 for the (*s′,* 1)-group. Thus, the weighted number of mutants is 3 · (2/4)+ 1 · (0/4) = 3/2, and the individual 2-fitness is *w*_2_(*s′|s,* 2) = (3/2)/2 = 3/4. Similarly, *w*_3_(*s′|s,* 2) = {3 · (1/6) + 1 · (0/6)*}/*2 = 1/4 and *w*_4_(*s′|s,* 2) = *w*_5_(*s′|s,* 2) = 0.

#### 2.3.3 Notation for perturbation analysis

Since our goal is to perform a sensitivity analysis of *ρ* to evaluate the selection gradient and disruptive selection coefficient, we assume that the mutant and resident trait values are close to each other and write

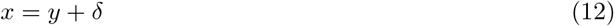

with *δ* sufficiently small (*i.e.*, *|δ| «* 1). Thus, *ρ* can be Taylor-expanded with respect to *δ*.

For invasion fitness *ρ*, or more generally, for any smooth function *F* that depends on *δ*, we will use the following notation throughout this paper. The Taylor-expansion of *F* with respect to *δ* is written as

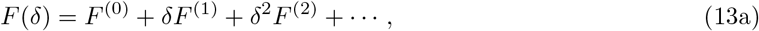

where *F* ^(*ℓ*)^ is given by

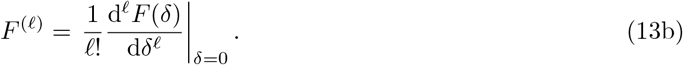

#### 2.3.4 Properties of the monomorphic resident population

The zeroth-order coefficient in eq.(13) corresponds to the situation where the function *F* is evaluated under the supposition that individuals labelled as “mutant” and “resident” are the same. In that case, individuals in groups with the same state are assumed to be exchangeable in the sense that they have the same reproductive characteristics (the same distribution of fitnesses, *i.e.*, the same mean fitness, the same variance in fitness, and so on). This results in a neutral evolutionary process, *i.e.*, a monomorphic population.

We now characterize the mutant lineage dynamics under a neutral process as this plays a crucial role in our analysis. From eq.(10), the individual 1-fitness in an (*s, i*)-group, written under neutrality, equals

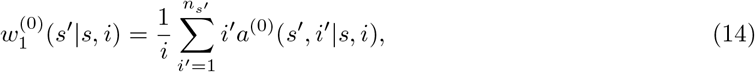

where each *a*^(0)^(*s′, i′|s, i*) is an entry of the matrix ****A**** under neutrality. By our exchangeability assumption, eq.(14) does not depend on *i*, the number of the individuals labeled as “mutants” in this group (see section A.2 (iv) in the Supplementary Material). If this would not be the case, mutants in a group (*s, i*_1_) and in a group (*s, i*_2_) with *i*_1_ ≠ *i*_2_ would have different reproductive outputs and mutants and residents would not be exchangeable. Therefore, from now on we write 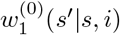 simply as 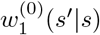. We collect these neutral fitnesses in the *N × N* matrix 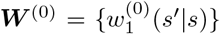. Its entry (*s′, s*) gives the expected number of descendants (possibly including self through survival) settling in groups of state *s′* that descend from an individual residing in an *s*-group (mutant or resident since they are phenotypically indistinguishable).

The assumptions that each group is density regulated (see Section 2.1) and that the resident population has reached stationarity guarantee that the largest eigenvalue of ****W**** ^(0)^ equals 1 (see section A.2 (v) in the Supplementary Material). This is the unique largest eigenvalue because ****W**** ^(0)^ is primitive due to the assumption that ****A**** is primitive. Thus, there is no demographic change in populations in which all individuals carry the same trait *y* and that have reached stationarity.

The fact that under neutrality 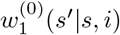 is independent of *i* and ****W**** ^(0)^ has the unique largest eigenvalue of 1 imposes constraints on the matrix ****A****^(0)^ = {*a*^(0)^(*s′, i′|s, i*)} that describes the growth of a mutant lineage under neutrality. Let us denote the left eigenvector of ****W**** ^(0)^ corresponding to the eigenvalue 1 by ****v****^(0)^ = {*v*^(0)^(*s*)}, which is a strictly positive row vector of length *N*. Each entry *v*^(0)^(*s*) gives the reproductive value of an individual in state *s*, which is the asymptotic contribution of that individual to the gene pool. Note that *v*^(0)^(*s*) does not depend on *δ* because it is defined from ****W**** ^(0)^, which is independent of *δ*. We now construct a row vector 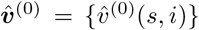 of length 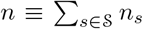 by setting 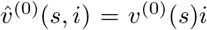. It has been shown that 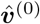 is a positive left eigenvector of the matrix ****A****^(0)^ = {*a*^(0)^(*s′, i′|s, i*)} corresponding to the eigenvalue 1, and therefore – since 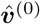 is strictly positive – the Perron-Frobenius theorem implies that the largest eigenvalue of ****A****^(0)^ is *ρ*^(0)^ = 1 (see Appendix A in Lehmann et al., 2016, for a proof and more details). We also show that the column vector {*q*^(0)^(*s*)} of length *N*, denoting the stable asymptotic distribution given by eq.(4) under neutrality, is the right eigenvector of the matrix ****W**** ^(0)^ corresponding to the eigenvalue of 1 (see section C.2.1 in the Supplementary Material). There is freedom of choice for how to normalize the left eigenvector ****v****^(0)^ and here we employ the convention that 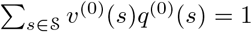. This means that the reproductive value of a randomly sampled mutant individual from its lineage is unity.

To summarize, under neutrality, the stable asymptotic distribution of mutants and the reproductive value of individuals satisfy

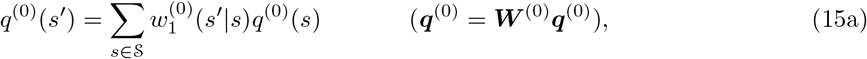

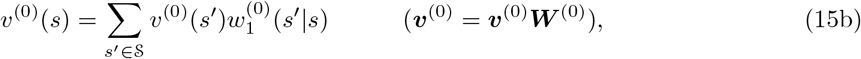

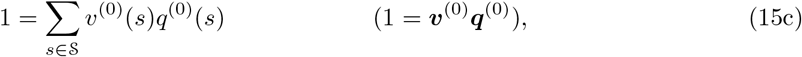

where ****v****^(0)^ is a row-vector with entries *v*^(0)^(*s*) and ****q****^(0)^ is a column-vector with entries *q*^(0)^(*s*).

### 2.4 Invasion fitness as reproductive-value-weighted fitness

Equation (2) for the leading eigenvalue and eigenvector of the matrix ****A**** can be left-multiplied on both sides by any non-zero vector of weights. This allows to express *ρ* in terms of this vector of weights and ****A**** and ****u****. If one chooses for the vector of weights the vector of neutral reproductive values 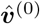 discussed above, then invasion fitness can be expressed as

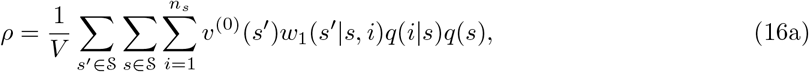

where

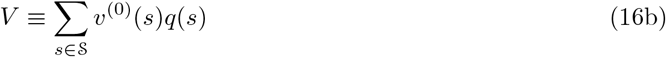

(see Lehmann et al., 2016, Appendix C, eq.(C.5), for the proof). This representation of *ρ* is useful to do concrete calculations. The intuition behind it is as follows. The inner sum, taken over *i*, represents the reproductive-value-weighted average number of offspring in states *s′* given a parental mutant resides in an *s*-group, where the average is taken over all possible mutant numbers experienced by the parental mutant in an *s*-group. The middle sum takes the average over all states *s* in which mutants can reside in the parental generation, and the outer sum takes the average over all possible states *s′* in which mutant offspring can reside (possibly including parents through survival).

Hence, the numerator in eq.(16a) is the reproductive-value-weighted average individual 1-fitness of a mutant individual randomly sampled from the mutant lineage, while the denominator *V* can be interpreted (in force of eq.(15b)) as the reproductive-value-weighted average of the neutral 1-fitness of an individual sampled from the asymptotic state distribution of the mutant lineage. Hence, *ρ* is the ratio of the reproductive-value-weighted average fitness of a mutant individual and that of a mutant individual under neutrality where both individuals are sampled from the same distribution. Note that in eq.(16a) the quantities *w*_1_(*s′|s, i*), *q*(*s*) and *q*(*i|s*) depend on *δ* while *v*^(0)^(*s′*) does not.

Our goal is to compute from eq.(16a) the selection gradient and disruptive selection coefficients,

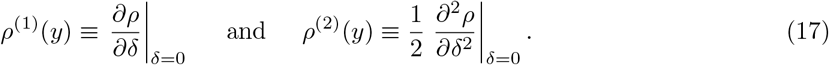

These coefficients are all we need to classify singular strategies (Metz et al., 1996; Geritz et al., 1998). Indeed, a singular strategy *y\** satisfies

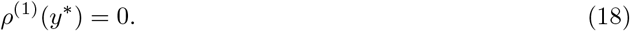

This strategy is locally convergence stable (*i.e.*, a local attractor point of the evolutionary dynamics) when

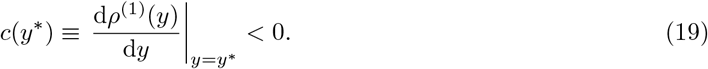

Note that convergence stability hinges on mutants with small phenotypic deviation *δ* invading and substituting residents (“invasion implies substitution”), which holds true when *|δ| «* 1 under the demographic assumptions of our model (Rousset, 2004, pp. 196 and 206). Furthermore, the singular point is locally uninvadable if

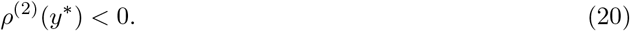

A singular strategy can then be classified by determining the combination of signs of the disruptive selection coefficient *ρ*^(2)^(*y\**) and the convergence stability coefficient *c*(*y\**) at *y\** (Metz et al., 1996; Geritz et al., 1998).

## 3 Sensitivity analysis

### 3.1 Eigenvalue perturbations

Using eq.(16a), as well as the normalization of reproductive values given in eq.(15c), we show in section B in the Supplementary Material that the first-order perturbation of *ρ* with respect to *δ* is given by

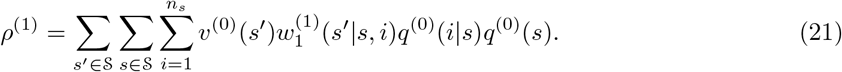

Thus, *ρ*^(1)^ is simply a weighted perturbation of individual 1-fitnesses *w*_1_. For the second-order perturbation of *ρ* with respect to *δ*, given that *ρ*^(1)^ = 0, we find that

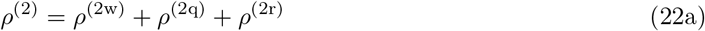

where

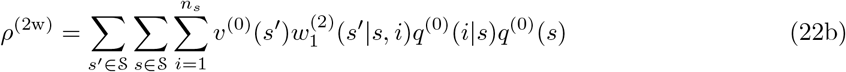

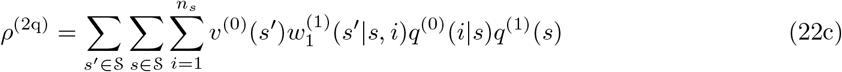

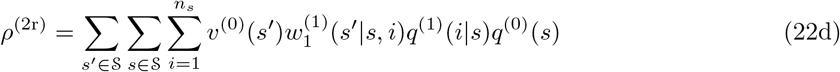

(section B in the Supplementary Material). The first term, labelled *ρ*^(2w)^, comes from the secondorder perturbation of individual 1-fitnesses. The second term, labelled *ρ*^(2q)^, comes from the first-order perturbation of the stationary distribution of mutants in the different states, and the third term, labelled *ρ*^(2r)^, comes from the first-order perturbation of the stationary distribution of the number of mutants in any given state.

While eqs.(21) and (22) give some insights into how selection acts on mutants, in particular, they emphasize the role of selection on the distributions *q*(*s*) and *q*(*i|s*), these expressions remain complicated as they involve weighted averages of fitness derivatives 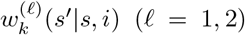 over the neutral and perturbed mutant distributions *q*^(1)^(*i|s*) and *q*^(1)^(*s*). To obtain more insightful expressions for these sensitivities, we express in the next section *w_k_*(*s′|s, i*) for *k* = 1, 2, 3 in terms of trait values. This will allow us to carry out rearrangements and simplifications of *ρ*^(1)^ and *ρ*^(2)^.

### 3.2 Individual fitness functions

#### 3.2.1 Individual 1-fitness

Consider a focal individual in a focal group in state *s* and denote by *z*_1_ the trait value of that individual. Suppose that the other *n_s_ −* 1 neighbors adopt the trait values 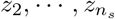 and almost all individuals outside this focal group adopt the trait value *z*. Let then

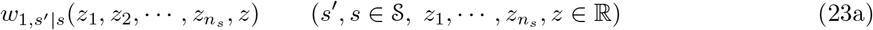

be the expected number of offspring in state *s′* that descend from a focal in state *s*. Equation (23a) expresses individual 1-fitness in terms of the phenotypes of all interacting individuals and will be referred to as an individual fitness function. It is a common building block of phenotypic models (see Frank, 1998; Rousset, 2004, for textbook treatments) and is the fitness that has to be considered if an exact description of a population is required, for instance, in an individual-based stochastic model, where each individual may have a different phenotype.

Because the only heterogeneity we consider are the different group states (we have no heterogeneity in individual states within groups), the individual 1-fitness function *w*_1,*s*′*|s*_ is invariant under permutations of 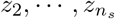. With this, we can rewrite eq.(23a) as

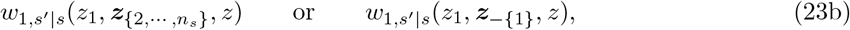

where the set-subscripted vector 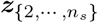 represents a vector of length *n_s_ −*1 in which each of 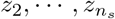 appears in an arbitrary order but exactly once. The subscript *−{1*} is used as a shorthand notation of the set difference {1, 2, *· · ·, n_s}_* \ {1} = {2, *· · ·, n_s_*} and used when the baseline set {1, 2, *· · ·, n_s_*} is clear from the context. Therefore, ***z****_−{1}_* is the same as 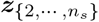. Similarly, in the following the subscript *−*{1, 2} represents the set difference {1, 2, *· · ·, n_s_*} \ {1, 2} = {3, *· · ·, n_s_*}, and so forth. For example, 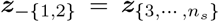 represents a vector of length *n_s_ −* 2 in which each of 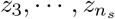 appears in an arbitrary order but exactly once.

For our two allele model *z_i_, z ∈ {x, y*}, we can write a mutant’s individual 1-fitness as

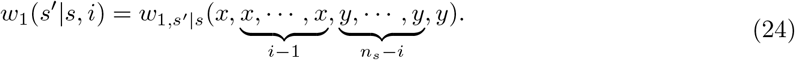

By using the chain rule and permutation invariance, the zeroth, first, and second order perturbations of *w*_1_(*s′|s, i*) with respect to *δ* are

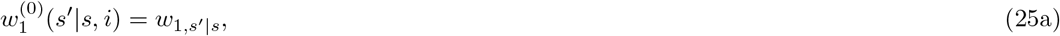

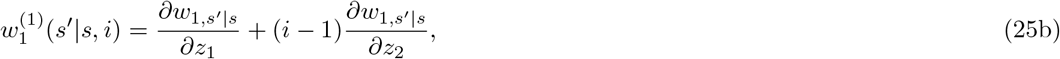

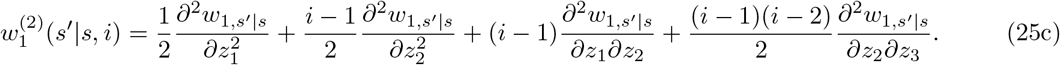

Here, all functions and derivatives that appear without arguments are evaluated at the resident population, (*y, · · ·, y*), a convention we adopt throughout. Note that some derivatives appearing in eqs.(25) are ill-defined for *n_s_* = 1 and *n_s_* = 2, but they are always nullified by the factors (*i −* 1) and (*i −* 1)(*i −* 2). Thus, by simply neglecting these ill-defined terms, eq.(25) is valid for any 1 *≤ i ≤ n_s_*.

#### 3.2.2 Individual 2- and 3-fitness

Consider again a focal individual with trait value *z*_1_ in a group in state *s* in which the *n_s_ −* 1 group neighbors have the trait values 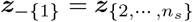 in a population that is otherwise monomorphic for *z*.

For this setting, we define two types of individual 2-fitness functions. First, let

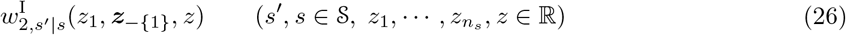

be the expected number of offspring in state *s′* that descend from the focal individual and that have a random neighbor that also descends from the focal individual (see Figure 2). Intuitively speaking, 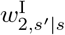 measures the number of sibling pairs produced by a focal individual. Hence, when one considers the reproductive process backward in time, 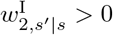 means that coalescence events do occur. We call 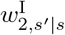 the “same-parent individual 2-fitness”, because the offspring involved in it descend from the same individual.

Second, for *n_s_ ≥* 2 consider a neighbor of the focal individual with trait value *z*_2_, called the *target* individual, in a group in which the remaining *n_s_ −* 2 neighbors have the trait profile 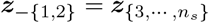.

**Figure 2:**
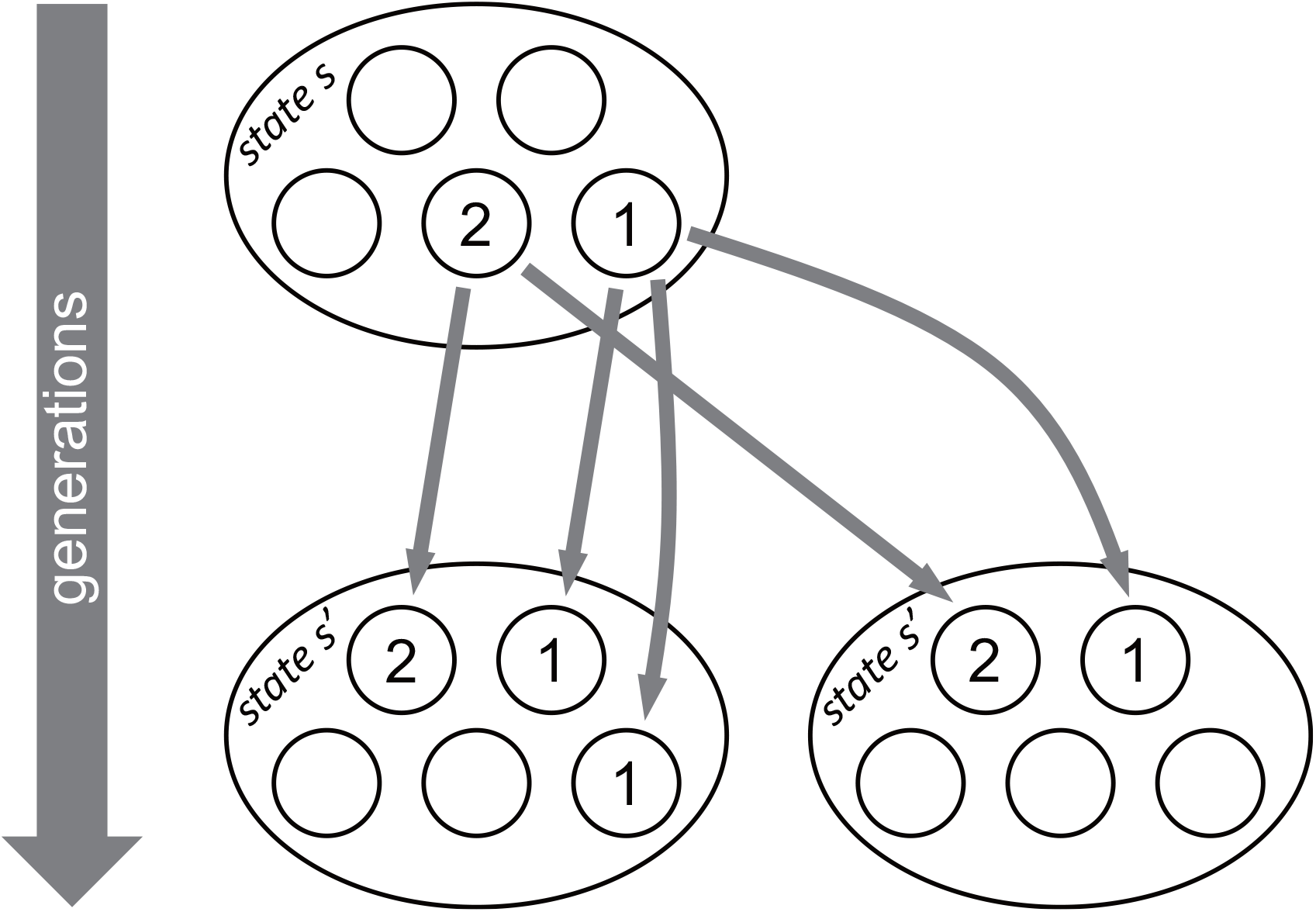
A schematic example of how we calculate the individual 2-fitnesses 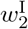 and 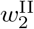. Gray arrows represent reproduction (or survival). We label by “1” the focal individual with trait value *z*_1_ in the parental generation and its offspring (possibly including self through survival) in the following generation. Similarly, we label by “2” the target individual with trait value *z*_2_ in the parental generation and its offspring (possibly including self through survival) in the offspring generation. Because each of the two descendants of the focal individual in the bottom-left group (those with label “1”) finds with probability 1/4 a random neighbor whose label is “1”, whereas the one descendant of the focal individual in the bottom-right group in the offspring generation finds no neighbors whose label is “1”, the same-parent individual 2-fitness of the focal is calculated as 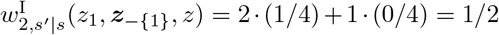. Similarly, because each of the two descendants of the focal individual in the bottom-left group finds a random neighbor whose label is “2” with probability 1/4, and because the one descendant of the focal in the bottom-right group finds a random neighbor whose label is “2” with probability 1/4, the different-parent individual 2-fitness of the focal is 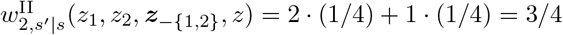.

Let

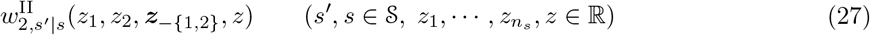

be the expected number of offspring in state *s′* that descend from the focal individual with trait value *z*_1_ and that have a random neighbor that descends from the *target* individual with trait value *z*_2_ (see Figure 2). We call 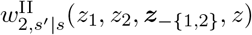 the “different-parent individual 2-fitness”, because the offspring involved in it descend from two different pre-selected individuals, which can thus collectively be thought of as the focal set of individuals under consideration. We note that this fitness function is invariant under the permutation of the trait values *z*_1_ and *z*_2_ of individuals from the focal set^2^ and it is also invariant under the permutation of the trait values in ***z***_−{1,2}_. But since 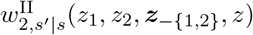 counts the offspring number (of a certain type) per individual with trait *z*_1_, 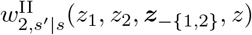 is a type of individual fitness.

Using the notation of mutant and resident phenotypes we have for 2 *≤ i ≤ n_s_* that

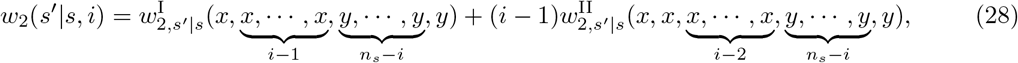

because a mutant neighbor of an offspring of a focal mutant either descends from the focal itself or is an offspring of one of the *i −* 1 mutant neighbors of the focal. The zeroth and first order perturbations of *w*_2_(*s′|s, i*) with respect to *δ* are given by

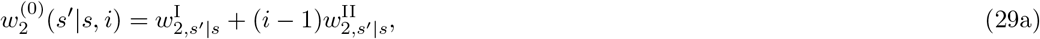

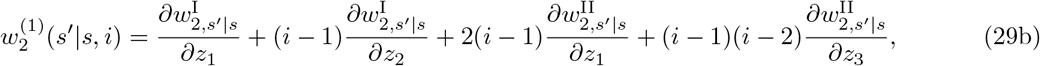

where the derivatives 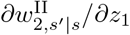 and 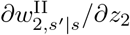 (the latter is equal to 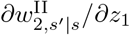 due to the permutation invariance, and hence the coefficient “2” appears in eq.(29b)) involve the trait values of the individuals of the focal set and 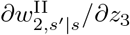 involves the trait values of a third individual. Note that some derivatives in eqs.(29) are ill-defined for *n_s_* = 1, 2 but they are always nullified by the factor (*i −* 1) or (*i −* 1)(*i −* 2). Thus, by simply neglecting these ill-defined terms eq.(29) is valid for any 1 *≤ i ≤ n_s_*.

Following the same line of reasoning as for individual 1- and 2-fitness, we similarly define three different types of individual 3-fitness. See section D in the Supplementary Material for more detailed explanations. Specifically, 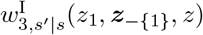 is defined as the expected number of offspring in state *s′* that descend from a focal individual in state *s* with trait value *z*_1_ and that have two random neighbors sampled without replacement both descending from the focal individual. Furthermore, 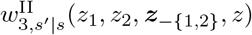 is defined as the expected number of offspring in state *s′* that descend from the focal individual in state *s* with trait value *z*_1_ and with two random neighbors sampled without replacement both descending from a target individual with trait value *z*. Finally, 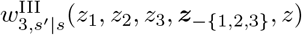 is defined as the expected number of offspring in state *s′* that descend from the focal individual in state *s* with trait value *z*_1_ with two random neighbors sampled without replacement, one of which descends from a first target individual with trait value *z*_2_ and the other descends from a second target individual with trait value *z*_3_. With these definitions, we show in section D in the Supplementary Material that the zeroth-order perturbation of 3-fitness *w*_3_(*s′|s, i*) with respect to *δ* is given by

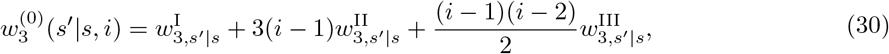

where 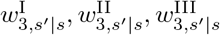 are those three different individual 3-fitness functions evaluated in a resident monomorphic population, (*y, · · ·, y*).

### 3.3 Sensitivity results

We now write *ρ*^(1)^ and *ρ*^(2)^ from Section 3.1 in terms of the just defined derivatives of the individual fitness functions.

#### 3.3.1 Selection gradient

By substituting eq.(25b) into eq.(21) we obtain

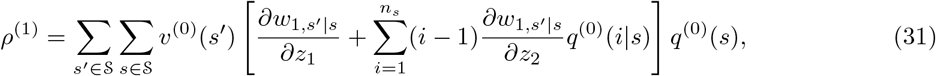

and by applying eq.(8) to the second term in square brackets we obtain

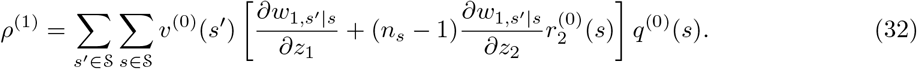

Thus, in order to be able to evaluate *ρ*^(1)^ it is sufficient to compute the neutral pairwise relatedness 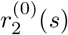 while the explicit evaluation of the *q*^(0)^(*i|s*) distribution is not needed. It is indeed a well-known result that the selection gradient *ρ*^(1)^ can be expressed in terms of reproductive values and relatedness-weighted fitness derivatives (see Frank, 1998; Rousset, 2004, for textbook treatments) and where *q*^(0)^(*s*) and *v*^(0)^(*s*) are given by eq.(15) with 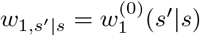.

Equation (32) can be interpreted as the expected first-order effect of all members of a lineage changing to expressing the mutant allele on the fitness of a focal individual that is a random member of this lineage. The recipient is sampled from state *s* with probability *q*^(0)^(*s*) and the derivative in the first term in the square brackets of *ρ*^(1)^ is the effect of the focal changing its own trait value on its individual fitness. The derivative in the second term in the square brackets describes the effect of the group neighbors of the focal changing their trait value on the focal’s individual fitness. This term is weighted by pairwise neutral relatedness since this is the likelihood that any such neighbor carries the same allele as the focal in the neutral process. Equation (32) is the inclusive fitness effect of mutating from the resident to the mutant allele for a demographically and/or environmentally structured population and the term in brackets can be thought of as the state-*s*-specific inclusive fitness effect on offspring in state *s′*. Equation (32) has previously been derived by Lehmann et al. (2016, Box 2) and is in agreement with eqs.(26) and (27) of Rousset and Ronce (2004), who derived the first-order perturbation *ρ*^(1)^ in terms of other quantities under the assumptions of fluctuating group size.

We show in section E in the Supplementary Material that by substituting eq.(29a) into eq.(C15), pairwise relatedness (eq.(8)) under neutrality satisfies the recursion

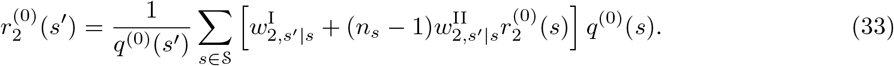

This expression for 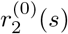, formulated in terms of individual 2-fitnesses, is novel but is in full agreement with previous results. In particular, eq.(29) of Rousset and Ronce (2004) can be shown to reduce to eq.(33) (see section G.2 in the Supplementary Material for a proof of this connection).

In summary, consistent with well established results, we present a biologically meaningful representation of *ρ*^(1)^. The ingredients in this representation can be obtained from the three systems of linear equations defined by eqs.(15a), (15b) and (33). This system of equations is fully determined once the individual *k*-fitnesses functions for *k* = 1, 2, namely, 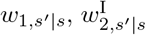, and 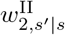 are specified for a resident population, and the *k*-fitness functions can usually be evaluated once a life-cycle has been specified. The dimension of this combined equation system has maximally three times the number of states *N*. This is significantly lower than the dimension of the matrix ****A**** we began with, especially, if group size *>* 10. In the next section, we extend these results to the disruptive selection coefficient.

#### 3.3.2 Disruptive selection coefficient

Assuming that *ρ*^(1)^ = 0 and substituting eq.(25) into eq.(22), rearrangements given in section E in the Supplementary Material show that

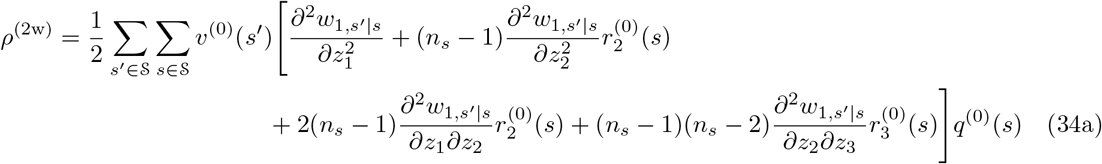

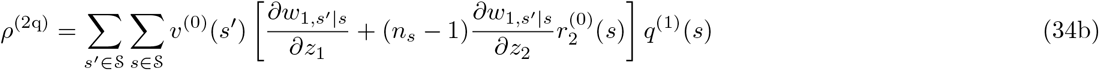

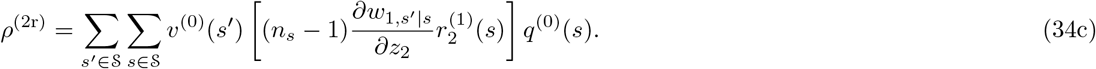

Equation (34a) depends on four different types of qualitative effects on the fitness of a focal individual: (i) The second-order effect on own fitness of the focal changing its trait value, which is positive, and then contributes to disruptive selection, if fitness is convex in own phenotype. (ii) The second-order effect resulting from the neighbors of the focal changing their trait values, which is positive if the focal’s fitness is convex in phenotype of group neighbors. This contributes to disruptive selection proportionally to pairwise relatedness 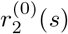, since this is the likelihood that a random neighbor carries the same allele as the focal individual. (iii) The joint effect of the focal individual and any of its neighbors changing their trait value, which is positive if the effect of increased trait values of own and others complement each other. This again contributes to disruptive selection in proportion to the likelihood that any neighbor is a mutant. (iv) The joint effect of pairs of neighbors of the focal changing their trait values, which is positive if the effect of increased trait values in neighbors complement each other. This contributes to disruptive selection with the probability 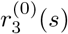 that a pair of neighbors carry the the same allele as the focal individual.

Equation (34b) depends, for each state, on the product of the state specific inclusive fitness effect (recall the term in brackets in eq.(32)) multiplied with the perturbation *q*^(1)^(*s*) of the group state probability. A contribution to disruptive selection occurs if the mutant allele increases its probability to be in a given state while simultaneously increasing the individual fitness of its carriers in that state. Similarly, eq.(34c) depends, for each state, on the product of the state specific indirect effect of others on own fitness (recall the second term in brackets in eq.(32)) and the relatedness perturbation 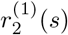. This contributes to disruptive selection if the mutant allele increases the probability that a focal has mutant neighbors while simultaneously increasing the individual fitness of those neighbors. Finally, we note that in the presence of a single state (*i.e.*, no state heterogeneity among groups) *ρ*^(2q)^ = 0. This is the case in all previously published expressions for the disruptive selection coefficient (Day, 2001; Ajar, 2003; Wakano and Lehmann, 2014; Mullon et al., 2016), which therefore reduce to *ρ*^(2w)^ + *ρ*^(2r)^ as defined by eqs.(34a) and (34c).

In order to compute *ρ*^(2)^ we need, in addition to eqs.(15a), (15b) and (33), expressions for 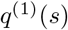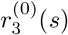, and 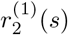. In section E in the Supplementary Material, we derive the corresponding recursions for *ρ*^(1)^ = 0. In particular, we show that *q*^(1)^(*s*) satisfies

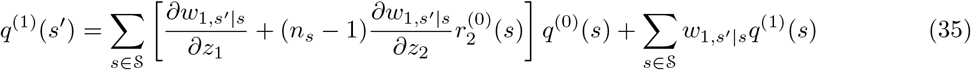

and that 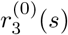 satisfies

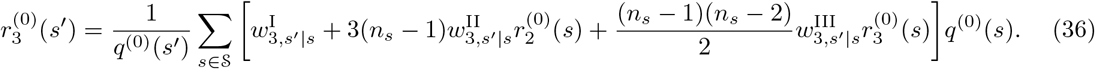

Finally, we show that 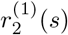 satisfies the recursion

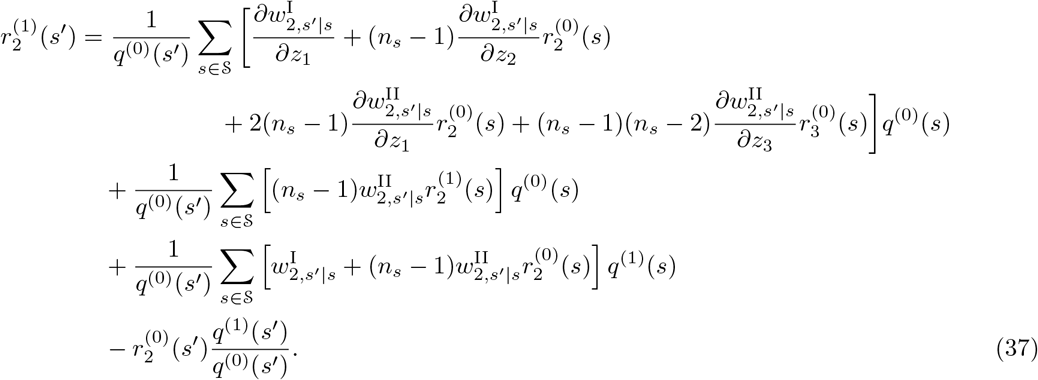

Equation (35) shows that *q*^(1)^(*s*) depends on the state-specific inclusive fitness effect (compare the first summand in eq.(35) to the term in brackets in eq.(32)). Thus, the probability that a mutant is in a certain state *s* increases with its state-specific inclusive fitness effect. Equation (36) for the three-way relatedness coefficient depends on 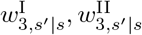 and 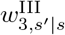 and it is a generalization of the pairwise relatedness coefficient given by eq.(33). Finally, eq.(37) shows that 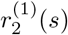 depends on direct and indirect effects on 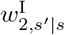 and 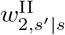. Note, that eq.(37) together with eqs.(15a), (15b), (33), (35), and (36) form a linear system of equations with a dimension equal to six times the number of states *N*. Its solution allows us to determine the disruptive selection coefficient *ρ*^(2)^. This system of equations in turn is fully determined once the the *k*-fitnesses for *k* = 1, 2, 3 are specified for a resident population, namely, 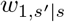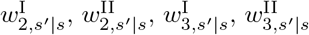, and 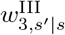.

In general, if the state space 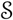 is large, solving this system of equations (and those needed for *ρ*^(1)^) may be complicated. Similarly, the 2- and 3-fitnesses may be complicated. We here give two directions for approximating *ρ*^(1)^ and *ρ*^(2)^. First, individual fitness generally depends on vital rates like fecundity and survival (see eqs.(45)–(46) for a concrete example) and variation of these vital rates may have small effects on fitness, which induces weak selection regardless of the magnitude of the phenotypic deviation *δ* (called “*ω*-weak selection” by Wild and Traulsen, 2007, and “weak payoff” by Van Cleve, 2015). For “weak payoffs” (or *ω*-weak selection), *ρ*^(2)^ *≈ ρ*^(2w)^ because one can neglect *ρ*^(2q)^ and *ρ*^(2r)^. Indeed, both these terms involve products of marginal changes in fitness, which implies that these products are of second-order effect under weak payoffs and first-order effects will thus dominate. Since *ρ*^(2w)^ only involves first-order effects it dominates the disruptive selection coefficient. See Van Cleve (2015) for an applications of this approximation to *ρ*^(1)^ and Wakano and Lehmann (2014) and Mullon et al. (2016) to *ρ*^(2)^. Second, variation of vital rates and fitness across states may be small under certain biological scenarios in which case one may apply a so-called small noise approximation (e.g., Tuljapurkar, 1990; Caswell, 2001) to *ρ*^(1)^ and *ρ*^(2)^, whereby the magnitude of variation are taken to be small. This simplification has been used to approximate *ρ*^(1)^ in a multi-species meta-population model that is covered by our general model (Mullon and Lehmann, 2018), but has not yet been applied to *ρ*^(2)^, which would be interesting in future work.

Finally, for some specific life-cycles the 2- and 3-fitness functions can be expressed in terms of components of the 1-fitness functions. This greatly simplifies the calculations because all recursions can then be solved explicitly. We will now provide an application of our model along this latter line, which still covers a large class of models.

## 4 Application to a lottery model with spatial heterogeneity

We now study a lottery model with overlapping generations and spatial heterogeneity. Such a model can be formulated for a variety of life-cycles and we here take a hierarchical approach in which we make increasingly more specific assumptions. Accordingly, this section is divided in three parts. Section 4.1 provides general results about the components of the selection coefficients based on the assumption of fixed group states *s*. In Section 4.2 we introduce two forms of population regulation resulting in hard and soft selection, respectively. Finally, in Section 4.3 we specify an explicit fitness function which allows us to present a fully worked example for the effect of group size and spatial heterogeneity on disruptive selection.

### 4.1 Spatial lottery model

#### 4.1.1 Decomposition into philopatric and dispersal components

We start by making the following three assumptions. (i) Group states *s* describe environmental variables that do not change in time. Thus, group states are fixed and we here refer to them as habitats. By *π_s_* we denote the relative proportion of groups in habitat *s*, hence 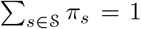. (ii) Individuals survive independently of each other with probability *γ_s_ <* 1 to the next time step in a group in habitat *s*. Note that *γ_s_* = 0 corresponds to the Wright-Fisher update where all adults die simultaneously, and that *γ_s_ ∼* 1 corresponds to the Moran update where at most one individual dies in a group. (iii) Dispersal occurs individually and independently to a random destination (no propagule dispersal). (iv) The evolving trait does not affect survival. With these assumptions we can decompose the 1-fitness of a focal individual into a philopatric and dispersal component as

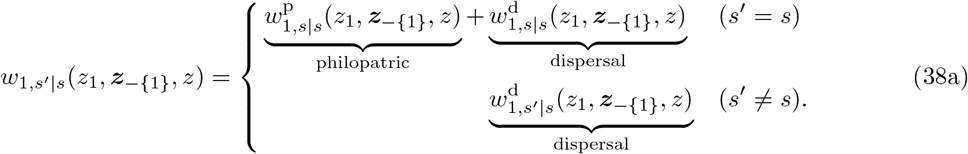

Offspring that have left from their natal group and successfully settled elsewhere are counted in the dispersal component 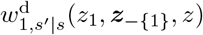. The philopatric component 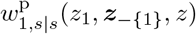 counts the number of non-dispersing offspring, possibly including self trough survival. Thus, we further decompose the philopatric part into a survival part and a reproduction part as

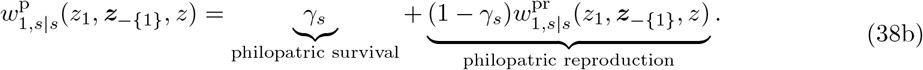

Similarly, for the dispersal part we write

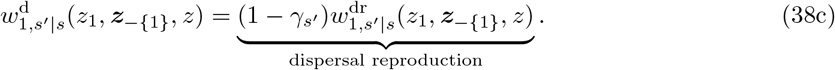

#### 4.1.2 General results for spatial lottery model

For this model, we explicitly compute the components of the selection gradient and disruptive selection coefficient in sections F.1 and F.2 of the Supplementary Material. In particular, we show that the probability that a random lineage member is sampled from a group in state *s* under neutrality equals the weighted frequency

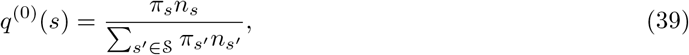

where the weights are the number of individuals in the group state.

For the reproductive value, it is instructive to provide a formula for *v*^(0)^(*s′*)*q*^(0)^(*s*), because the reproductive value always appears as a product with *q*^(0)^(*s*) in *ρ*^(1)^ (eq.(32)) and *ρ*^(2)^ (eq.(34)) (the only exception is eq.(34b), but see the discussion below eq.(43)). This product is given by

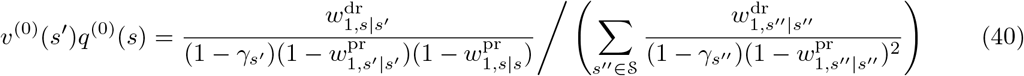

(section F.1 in the Supplementary Material). Furthermore, the neutral pairwise relatedness coefficient equals

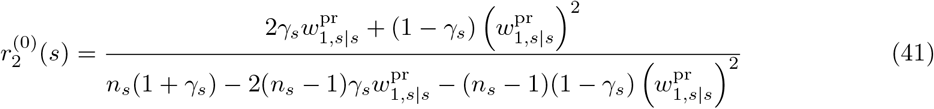

(section F.2 in the Supplementary Material). The general solution for 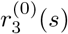 remains complicated (see eq.(F32) for the full expression), but for special cases it is

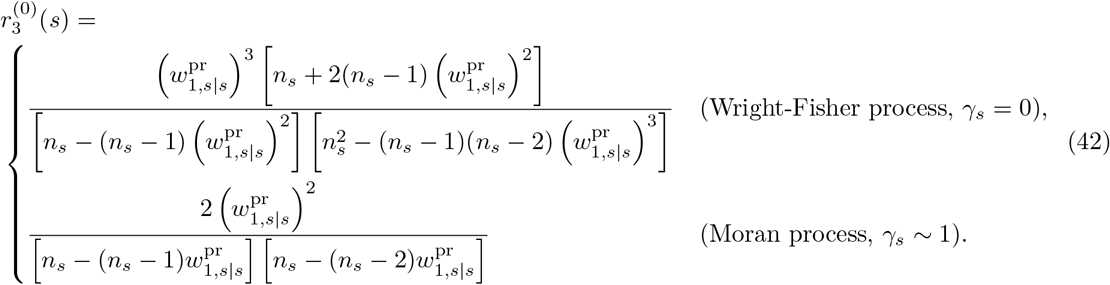

If the resident trait value is equal to the singular strategy where *ρ*^(1)^ = 0, then the first-order perturbation of the stationary mutant distribution is

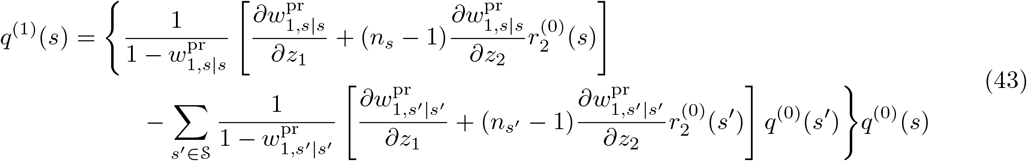

(section F.1 in the Supplementary Material). Note that we can obtain the fraction *q*^(1)^(*s*)*/q*^(0)^(*s*) by dividing both sides of eq.(43) by *q*^(0)^(*s*), which, when combined with eq.(40), allows to directly obtain the product *v*^(0)^(*s′*)*q*^(1)^(*s*). This quantity is required to compute eq.(34b). Finally, for *ρ*^(1)^ = 0 we have

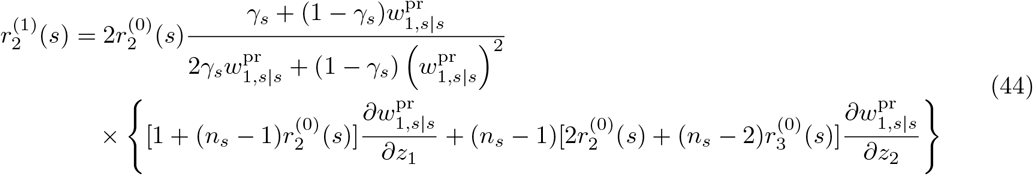

(see Section F.2 in the Supplementary Material where we also make the connection to previous work).

With eqs.(40) and (41) we can compute the first-order perturbation of invasion fitness, eq.(32), explicitly given specific life-cycle assumptions (since all recursions have been solved). Similarly, under the assumption that *ρ*^(1)^ = 0, and with eqs.(39)–(44) in hand, we can explicitly compute the second-order perturbation of invasion fitness, eq.(34).

### 4.2 Fecundity selection under two different forms of density regulation

We further refine our assumptions in order to arrive at two life-cycles with concrete expressions for 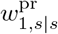 and 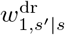. The first one is as follows. (1) Each adult individual in a group in habitat *s* produces on average a very large number *f_s_* of offspring, and then either survives with probability *γ_s_* or dies with the complementary probability. (2) Offspring disperse independently of each other to a uniformly randomly chosen non-natal group with the non-zero probability *m_s_*. An offspring survives dispersal with probability *p_s_* when dispersing from a group in habitat *s*. (3) All offspring aspiring to settle in a group in habitat *s* compete for the average number (1 *− γ_s_*)*n_s_* of breeding sites vacated by the death of adults and are recruited until all *n_s_* breeding sites are occupied. (4) The evolving trait does not affect dispersal.

In this life cycle, density-dependent population regulation occurs after dispersal when offspring aspire to settle and we refer to this regime as *hard selection*. We also consider a *soft-selection* variant in which density regulation occurs in two steps (as in Fig. 1 of Svardal et al., 2015). First, a local trait-dependent stage of density-dependent regulation occurs immediately after reproduction (after stage (1) in the above life cycle) in which the offspring pool in each group is brought back to a size proportional to the local group size *n_s_*, say size 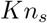, where *K* is a large number. From here on dispersal and recruitment (second regulation step) proceed as in the hard-selection life cycle.

For these two life cycles, the philopatric and dispersal fitness components can be written as

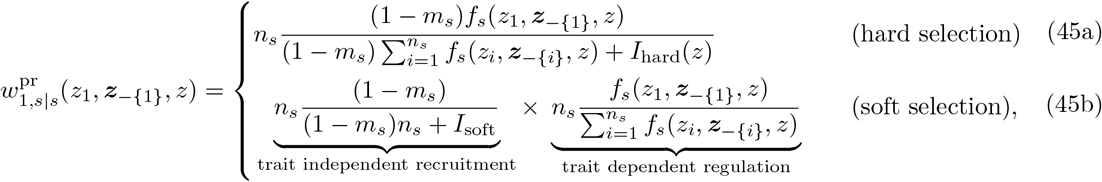

and

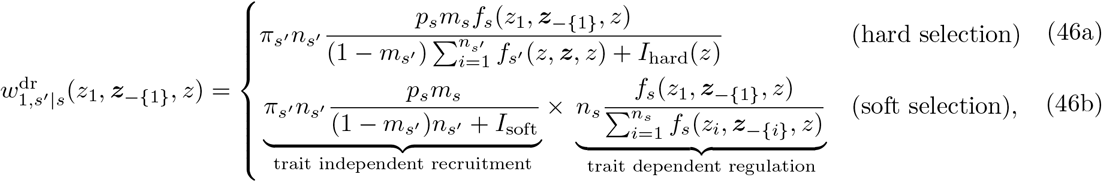

respectively, where *f_s_*(*z_i_, **z**_−{i}_, z*) is the fecundity of individual *i* in a group in habitat *s* and

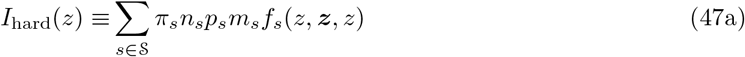

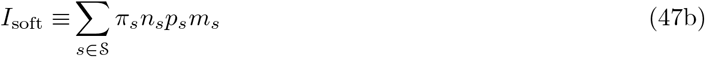

are the trait-dependent immigration terms for the hard-selection model and trait-independent immigration term for the soft selection model, respectively.

Equations (45b) and (46b) can be understood as follows. During the stage of trait-dependent regulation the local offspring pool in a group in habitat *s* is brought back to a size proportional to *n_s_*, namely *Kn_s_*, whereby the proportion of individuals among the surviving offspring descending from a focal individual is 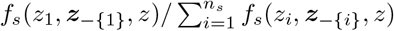. Each of these offspring either disperses or stays local and then competes to be recruited. With probability 1 *− m_s_* an offspring is philopatric, and this philopatric offspring gets recruited with probability 1/ [*K*((1 *− m_s_*)*n_s_* + *I*_soft_)] per open spot. Here *K*((1 *− m_s_*)*n_s_* + *I*_soft_) is the expected number of local competitors, where the number of migrant offspring competing in a given group for recruitment and coming from a group in habitat *s* is proportional to *π_s_n_s_p_s_m_s_*. Offspring dispersing to a group in habitat *s′* experience on average *K* ((1 *− m_s′_*)*n_s′_* + *I*_soft_) competitors and the probability to compete in such a group is *π_s′_*. The likelihood to be recruited (either after dispersing or without dispersing) is then multiplied by the expected number of open breeding sites, which equals *n_s_*(1 *− γ_s_*) in the natal group and *n_s′_* (1 *− γ_s′_*) in non-natal groups in habitat *s′*, but the factors (1 *− γ_s_*) and (1 *− γ_s′_*) are already accounted for in eqs.(38b) and (38c). Note that the constant *K* does not appear in eqs.(45b) and (46b) because it appears both in the numerator and denominator of these equations and thus cancels out.

Using eq.(45) and eq.(46) along with eqs.(39)–(44) allows to compute *ρ*^(1)^ and *ρ*^(2)^ for a large class of models. In sections G, H.3 and I.3 in the Supplementary Material, we show that we recover a number of previously published results belonging to this class of models, some of which were derived with quite different calculations (Pen, 2000; Ohtsuki, 2010; Lehmann and Rousset, 2010; Rodrigues and Gardner, 2012; Wakano and Lehmann, 2014; Svardal et al., 2015; Mullon et al., 2016; Parvinen et al., 2018). This indirectly confirms the validity of our calculations. For simplicity of notation we assumed that the evolving trait does neither affect survival nor dispersal (it only affects fecundity), extensions to include effects on survival and dispersal are in principle straightforward.

### 4.3 Selection analysis

In this section, we finally present explicit expressions for the selection gradient *ρ*^(1)^ and the coefficient of disruptive selection *ρ*^(2)^ for both the model of hard and soft selection. We then introduce an explicit fecundity function, which, under some additional symmetry assumptions, allows us to have a completely worked example.

#### 4.3.1 Hard selection

Inserting eqs.(45a) and (46a) into eqs.(38b) and (38c), respectively, we show in section H in the Supplementary Material that the selection gradient for the hard selection lottery model is

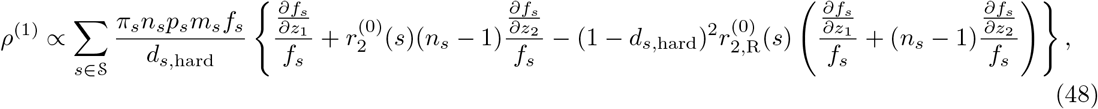

where the proportionality constant is positive (and given by the inverse of eq.(H4)) and *d_s,_*_hard_ is the backward migration rate from groups in state *s* under neutrality defined as

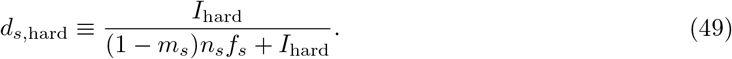

This rate depends on *y* because *I*_hard_ and *f_s_* are evaluated at (*y, · · ·, y*). Equation (48) further depends on

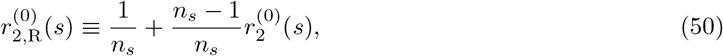

which is the relatedness between two individuals sampled with replacement in a group in state *s* and where

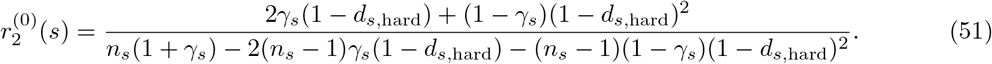

Equation (48) can be understood as follows. The first term in the curly brackets is the marginal fecundity effect by a focal individual on itself, while the second term is the marginal fecundity effect conferred by all group members to the focal individual weighted by the coefficient of pairwise relatedness. Finally, the third term reflects competition for the finite number of breeding spots in a group. A change in the trait value of a focal individual that increases its fecundity or that of its neighbors increases the strength of local competition. This reduces the fitness of the focal individual if the additional offspring remain philopatric and compete with own offspring. Equation (48) is a generalization of previous results obtained for the island model (see section H in the Supplementary Material for the detail of these connections).

Similarly, inserting eqs.(45a) and (46a) into eqs.(38b) and (38c), respectively, and using these in eq.(34), we obtain a general expression for the disruptive selection coefficient *ρ*^(2)^ under hard selection. The resulting expression, while useful for numerical calculations, is too lengthy to be presented here and we refer to section H in the Supplementary Material for details. Therein, we show that under a Wright-Fisher process (*γ_s_* = 0) the results of Parvinen et al. (2018) are recovered, who obtained an expression of *ρ*^(2)^ expressed in terms of first- and second-order derivatives of *f_s_*.

To complement these results and to approach a fully worked example, we assume a Moran process (*i.e.*, *γ_s_ ∼* 1) and that fecundity of an adult individual depends only on its own phenotype (*i.e.*, *f_s_*(*z*_1_, ***z**_−{_*_1}_, z) = *f_s_*(*z*_1_)). Under these assumptions, we show in section J.1 in the Supplementary Material that the selection gradient is a weighted sum of d*f_s_/*d*z*_1_ over different states *s* (see eq.(J1)), and that the disruptive selection coefficient is

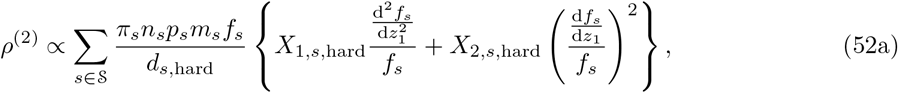

where the positive proportionality constant is the same as in eq.(48), and

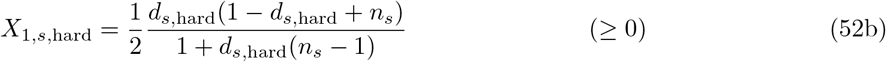

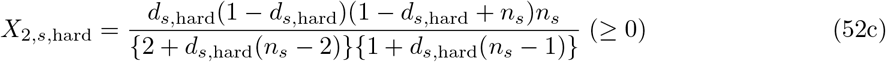

For complete dispersal (*i.e.*, *d_s,_*_hard_ = 1)^3^ we obtain that *X*_1,*s,*hard_ = 1/2 and *X*_2,*s,*hard_ = 0. As the dispersal rate *d_s,_*_hard_ decreases, the ratio *X*_2,*s,*hard_*/X*_1,*s,*hard_ increases monotonically. Hence, as dispersal becomes more limited, relatively more weight is put on the squared first-order derivative (d*f_s_/*d*z*_1_)^2^ compared to the second-order derivative 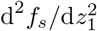, indicating that limited dispersal facilitates disruptive selection (and, if the singular strategy *y\** is convergence stable and remains so when varying dispersal, then evolutionary branching is facilitated). On the other hand, for a fixed *d_s,_*_hard_ *<* 1, the ratio *X*_2,*s,*hard_*/X*_1,*s,*hard_ monotonically decreases as group size decreases. Hence, with decreasing group size less weight is put on the squared first-order derivative (d*f_s_/*d*z*_1_)^2^, which acts to limit disruptive selection. We finally note that the functional form of eq.(52a) holds beyond the Moran process, provided all other assumptions are the same. While the weights will depend on the specifics of the reproductive process, we conjecture that the weights will feature the same qualitative dependence on dispersal and group size.

We now make two further assumptions. First, we follow Svardal et al. (2015) and assume that fecundity is under Gaussian stabilising selection with habitat specific optimum *y*_op,*s*_. Thus,

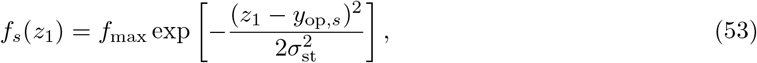

where *f*_max_ is the maximal fecundity of an individual and 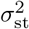 is inversely proportional to the strength of stabilising selection. Second, we assume that group size, migration and juvenile survival are identical for all habitats, *i.e.*, *n_s_* = *n*, *m_s_* = *m*, and *p_s_* = *p* for all *s*. Hence, habitats only differ in the trait value *y*_op,*s*_ that maximizes fecundity.

Under these assumptions, the singular strategy *y\** is implicitly given by

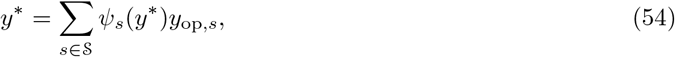

which is a weighted average of the habitat specific trait optima with the weights *ψ_s_* being complicated functions of the model parameters (see section J.1 in the Supplementary Material). The condition for the disruptive selection coefficient at the singular point *y\** (eq.(52a)) being positive can be expressed as

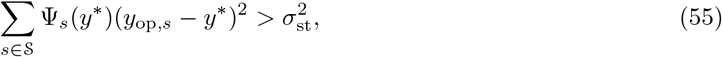

where the ψ_*s*_’s are again complicated weights (section J.1 in the Supplementary Material).

These expressions greatly simplify when we consider only two habitats with equal proportions, *i.e.* 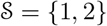 with *π*_1_ = *π*_2_ = 1/2, no mortality in dispersal, *p* = 1, and symmetric optima in the sense that *y*_op,2_ = *−y*_op,1_. Due to this symmetry, *y\** = 0 is a solution of eq.(53) and therefore a singular strategy. Furthermore, in section J.1 in the Supplementary Material, we find that under the aforementioned assumptions

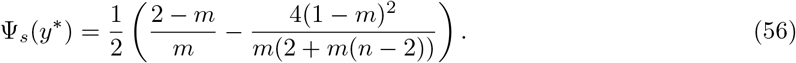

Then, by using the variance of the habitat optima defined by

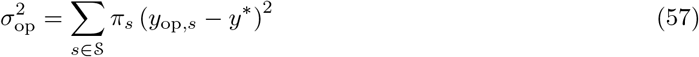

(in the current case, with *π*_1_ = *π*_2_ = 1/2), condition (55) can be written as

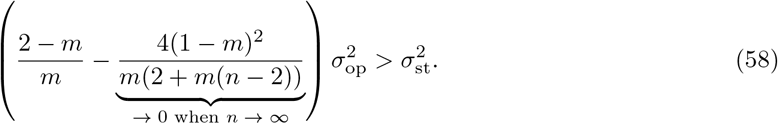

The first term in the parenthesis is the effect of limited dispersal on disruptive selection in the absence of kin selection (that is, under infinite group size). This term increases with decreasing dispersal, which facilitates disruptive selection. Indeed, low dispersal increases the probability that lineage members experience the same group-specific state favoring local adaptation. The second term in the parenthesis captures the effect of kin selection. The absolute value of this negative term increases with both decreasing dispersal and decreasing group size, which inhibits disruptive selection. This effect can be understood as follows. All philopatric offspring within a group compete with each other for the limited number of spots to settle within a group. Relatedness among individuals within a group increases with decreasing group size. Thus, in smaller groups competing individuals are more likely to be related with each other and this diminishes the benefit of mutations increasing adaptation to the group-specific state. This effect becomes more pronounced with decreasing dispersal since this increases relatedness within groups even more. We therefore expect that the singular point *y\** is more likely to be uninvadable for small groups and this is indeed what we observe in Figure 3, especially evident in panel (f). It can be shown that the effect of decreasing dispersal on the first term on the left-hand side of (58) dominates the effect on the second term. Thus, decreasing *m* indeed facilitates disruptive selection as illustrated in Figure 3(b-f).

**Figure 3:**
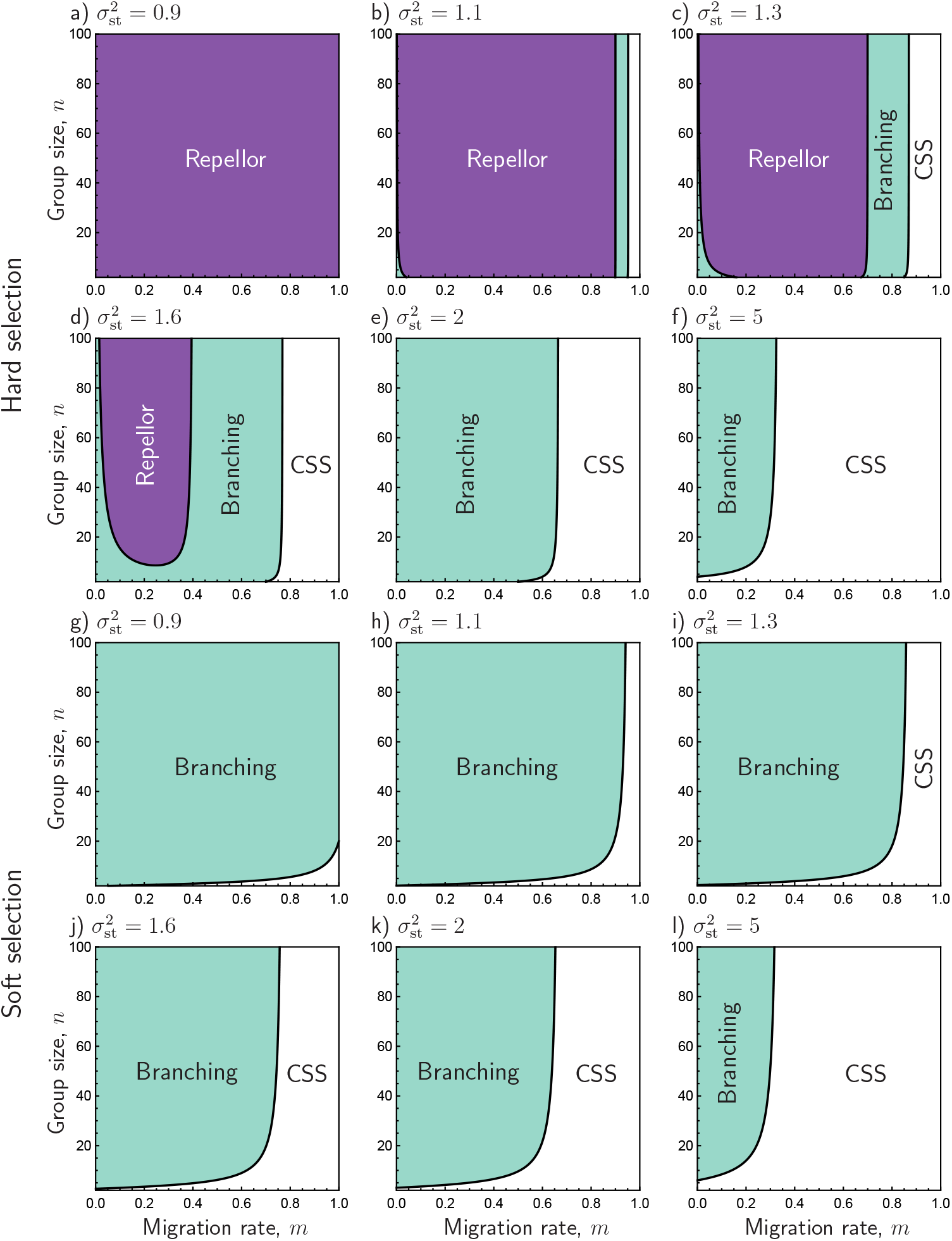
Bifurcation diagrams for the singular point *y\** = 0 as a function of the migration rate *m* (x-axis) and group size *n* (y-axis) for six different values of the within group selection parameter 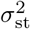 (see eq.(53)). (a-f) Hard selection, (g-l) soft selection. Purple: evolutionary repellor, blue: evolutionary branching point, white: uninvadable and convergence stable singular point, *i.e.*, continuously stable strategy (CSS). Other parameter values: *y*_op,1_ = 1 = *−y*_op,2_ (implying 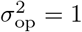).

In the limit of *m* = 0 and *m* = 1 the condition for the disruptive selection coefficient being positive (58) becomes

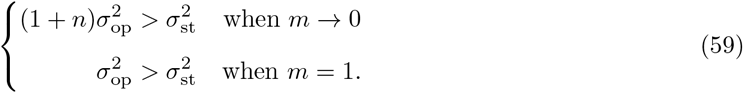

Thus, at very low dispersal the singular point changes from being uninvadable to invadable when group size exceeds 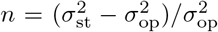 (as can be seen in Figure 3(f) where the boundary between CSS and branching point for very low *m* occurs at *n* = 4). At complete dispersal, the singular point is uninvadable for 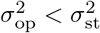 and invadable otherwise. Finally, the singular strategy is more likely to be under stabilizing selection the larger the ratio 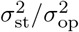, as is clearly illustrated in Figure 3(a-f).

A singular point at which selection is disruptive is an evolutionary branching point if it is also convergence stable. Substituting eq.(48) under all mentioned assumptions into eq.(19) we obtain after rearrangements that *y\** = 0 is convergence stable if

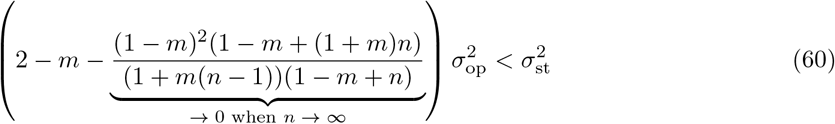

and repelling otherwise. From inspecting the left-hand side of this condition, the coefficient of 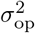 is a unimodal function of *m* and takes the minimum value 1 at *m* = 0, 1 and the maximum at

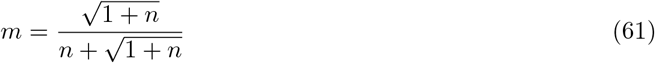

for any fixed *n*. Therefore, it is clear that 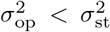 is a necessary but not sufficient condition for convergence stability. More generally, increasing 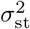 relative to 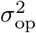 increases the space in the (*m, n*)-plane for which the singular point is convergence stable (*cf*. Figure 3(a-f)). In section J.1 in the Supplementary Material we show that 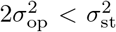 is a sufficient condition for convergence stability (*cf*. Figure 3(e-f)). Interestingly, from the unimodality above, the singular point can be repelling for intermediate values of *m* as can be seen in Figure 3(b-d). For large group size, condition (60) becomes 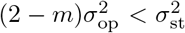 and therefore convergence stability changes at 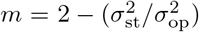, which coincides very well with where the singular point turns from convergence stable to repelling at group size *n* = 100 in Figure 3(b-d). For the effect of group size *n* on convergence stability, the coefficient of 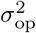 in condition (60) is, for any fixed 0 *< m <* 1, an increasing function of *n*. Thus, smaller group sizes are more favorable for convergence stability of the singular point *y\** = 0.

An immediate conclusion from these observations is that for *m* = 1 evolutionary branching does not occur under hard selection (with fecundity given by eq.(53)). This is so because for *m* = 1 competition is global and does not occur between individuals within a group. This removes any frequency-dependent selection effect. Indeed, under our assumptions setting *m* = 1 (and *p* = 1) in eq.(45a) and eq.(46a) results in 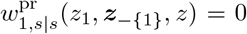 and 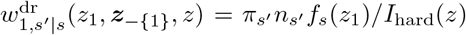 for all *s′* and *s*. Thus, there is no longer any state specific frequency-dependence, since *I*_hard_(*z*) is common to all fitness functions. In this case, the singular point is both convergence stable and uninvadable if 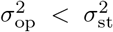 and both repelling and invadable if 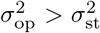. This is in agreement with the well-known finding that under hard selection and complete dispersal selection is frequency-independent and adaptive polymorphism cannot be maintained by spatial heterogeneity alone (Dempster, 1955; Ravigné, 2004; Ravigné et al., 2009; Débarre and Gandon, 2011).

#### 4.3.2 Soft selection

Inserting eqs.(45b) and (46b) into eqs.(38b) and (38c), respectively, we show in section I in the Supplementary Material that the selection gradient for the soft selection lottery model is

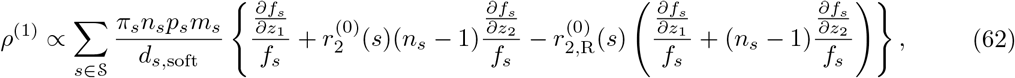

where the positive proportionality constant is positive (and given by the inverse of eq.(I4)) and

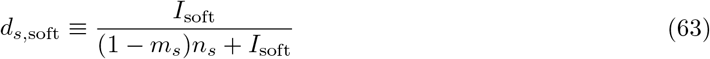

is the backward migration rate from groups in habitat *s* under neutrality. In contrast to the case of hard selection, eq.(63) is independent of *y*. Pairwise relatedness under neutrality 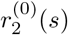 takes the same form as in eq.(51) where all *d_s,_*_hard_ have to be replaced with *d_s,_*_soft_. The key difference between eq.(48) and eq.(62) is that under soft selection the competition term is larger than under hard selection because the weighting by the backward dispersal probability has disappeared in the latter case. This reflects the fact that under soft selection density regulation occurs before dispersal. Again, eq.(62) is a generalization of previous results as detailed in section I in the Supplementary Material.

Similarly, inserting eqs.(45b) and (46b) into eqs.(38b) and (38c), respectively, and using these in eq.(34), we obtain a general expression for the disruptive selection coefficient *ρ*^(2)^ under soft selection. As was the case for hard selection, the resulting expression can be useful for numerical calculations, but is too lengthy to be presented here and we refer to section I in the Supplementary Material for details.

Paralleling the analysis under hard selection, we assume a Moran process (*i.e.*, *γ_s_ ∼* 1) and that the fecundity of adult individuals depends only on their own phenotype (*f_s_*(*z*_1_, ***z**_−{_*_1}_, *z*) = *f_s_*(*z*_1_)). Under these assumptions we show in section J.2 in the Supplementary Material that

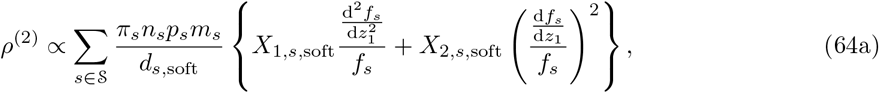

where the positive proportionality constant is the same as in eq.(62), and

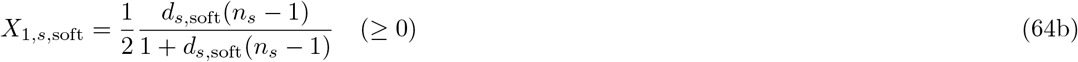

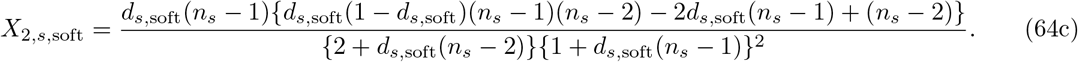

The ratio of these weights, *X*_2,*s,*soft_*/X*_1,*s,*soft_, shows qualitatively the same behavior as the corresponding expressions under hard selection (eqs.(52b) and (52c)) with respect to changes in *d_s,_*_soft_ and *n_s_*. However, a notable difference from the hard selection case is that *X*_2,*s,*soft_ (and hence the ratio, *X*_2,*s,*soft_*/X*_1,*s,*soft_) can be negative for small *n_s_* and large *d_s_*. We finally note that, as was the case for eq.(52a), the functional form of eq.(64a) holds beyond the Moran process, provided all other assumptions are the same.

Under the assumption of Gaussian fecundity selection (eq.(53)) and *n_s_* = *n*, *m_s_* = *m*, *p_s_* = *p* = 1 for all states *s*, which entails *d*_soft_ = *m*, we again obtain a fully worked example. The value *y\** for the singular strategy is given by the average habitat optimum,

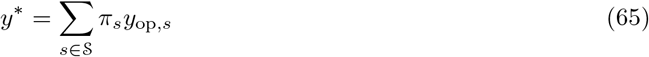

(section J.2 in the Supplementary Material). Furthermore, the coefficient of disruptive selection is positive if and only if

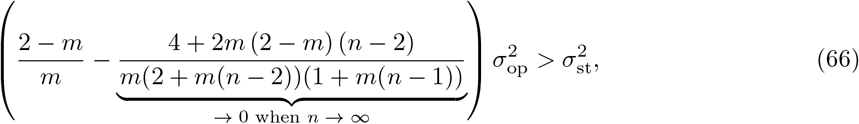

where 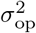 is the variance in the habitat optima defined by eq.(57). Note that condition (66) is valid only for *n ≥* 2 (because otherwise eqs.(64b) and (64c) evaluate to zero). The two terms in parenthesis on the left-hand side of condition (66) have the same interpretation as the corresponding terms in condition (58) for the case of hard selection and they respond in the same direction with respect to changes in dispersal *m* and group size *n*. In the limit of infinitely large group size (*n → ∞*) the second term vanishes and we recover eq.(C.15) of Svardal et al. (2015).

In section J.2 in the Supplementary Material, we show that *y\** as given by eq.(65) is convergence stable for any value of 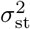 and 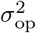 and independent of group size *n* and dispersal probability *m*. Thus, the singular point is an evolutionary branching point when it is invadable and an endpoint of the evolutionary dynamics (continuously stable strategy, CSS) when uninvadable. For the special case of only two habitats with *y*_op,1_ = 1 = *−y*_op,2_, Figure 3 shows how *n*, *m* and 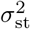 determine whether *y\** = 0 is a branching point or a CSS. In summary, stronger selection (smaller values of 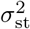), lower migration and larger groups favor adaptive diversification at an evolutionary branching point.

## 5 Discussion

The main result of this paper is an expression for the disruptive selection coefficient *ρ*^(2)^ in heterogeneous group-structured populations (eq.(34)). We show that *ρ*^(2)^ depends on three types of differentials: (a) the first- and second-order perturbations of the expected number of offspring in different states produced by an individual in a given state, (b) the first-order perturbation of the probability that an individual is in the different states, and (c) the first-order perturbation of the probability that a randomly sampled neighbor of an individual carries alleles identical by descent (perturbation of relatedness). These differentials depend on and are weighted by three quantities evaluated under neutrality: (i) the reproductive values *v*^(0)^(*s*) of individuals in state *s*, (ii) the pairwise and three-way relatedness coefficients 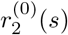 and 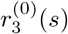 in state *s*, and (iii) the probability *q*^(0)^(*s*) that a randomly sampled individual resides in a group in state*s*.

At a conceptual level, our results about the components of *ρ*^(2)^ can be thought of as a direct extension of the result that the three types of neutral weights – reproductive values, relatednesses, and probabilities of occurrence in state *s* – are needed to evaluate the selection gradient *ρ*^(1)^ for quantitative traits in group-structured populations (Taylor and Frank, 1996; Frank, 1998; Rousset, 2004). All the above mentioned differentials and their weights can be obtained by solving systems of linear equations that are at most of dimension *N*, *i.e.*, the number of states groups can be in. This represents a significant reduction compared to the dimension of the state space of the original evolutionary process, which is equal to the dimension of the mutant transition matrix ****A****.

A distinctive and novel feature of our analysis is the introduction of the concept of individual *k*-fitness, *w_k_*(*s′|s, i*), which describes the expected number of descendants of a mutant in an (*s, i*) group (possibly including self through survival) that settle in state-*s′* groups and have *k −* 1 randomly sampled neighbors that are also mutants (*i.e.*, that descend from the same common ancestor). In the context of our perturbation analysis, we show that *w_k_*(*s′|s, i*) can be themselves expressed in terms of individual *k*-fitness functions for *k* = 1, 2, 3 where individuals are labelled as focal, group neighbor and population member, and which are sufficient to evaluate all aforementioned quantities and thus *ρ*^(1)^ and *ρ*^(2)^ (see sections 3.2.1–3.3). These latter individual *k*-fitness functions do not depend on the mutant type and provide for *k* = 2, 3 the generalizations of the fitness functions for *k* = 1 already in use in the direct fitness method (Taylor and Frank, 1996; Frank, 1998; Rousset, 2004). They are thus sufficient biological ingredients to determine whether or not disruptive selection occurs. In a well-mixed populations in which individuals do not interact with relatives only individual 1-fitness functions are required to evaluate *ρ*^(1)^ and *ρ*^(2)^. Individual 2- and 3-fitnesses describe the possibility that under limited dispersal the offspring of a given parent can have neighbors (here one or two) that belong to the same lineage and are thus more likely to have the same trait value than randomly sampled individuals from the population. This causes non-random mutant-mutant interactions, which is well known to critically affect the nature of selection on traits affecting own and others’ reproduction and survival (Hamilton, 1964; Michod, 1982; Frank, 1998; Rousset, 2004). Because the individual *k*-fitnesses describe group configurations in which offspring have neighbors that belong to the same lineage, the ancestral lineages of the *k* interacting individuals must coalesce in a common ancestor, and this can occur only if there is a non-zero probability that at least two individuals descend from the same parent over a generation (see section G.2 in the Supplementary Material for the connection to coalescence theory). Neutral relatedness in evolutionary models is indeed usually computed by using coalescence arguments and thus use a “backward” perspective on allele transmission (*e.g.* Taylor and Frank, 1996; Frank, 1998; Rousset, 2004). This may somewhat disconnect relatedness from the “forward” perspective of allele transmission induced by reproduction. Using individual 2-fitnesses to evaluate relatedness (see eq.(33)) brings upfront the connection between relatedness and reproduction (note that the “backward” approach may nevertheless be more useful for concrete calculations of relatedness).

As an application of our results, we analyze a lottery model with overlapping generations in heterogeneous habitats that allows for both hard and soft selection regimes. For this scenario, we show that *ρ*^(1)^ and *ρ*^(2)^ can in principle be solved explicitly (all systems of equation can be solved explicitly) but that generic expressions remain complicated functions, since they apply to any kind of social interactions (*i.e.*, any “game”) and different ecologies. In doing these calculations, we recover a number of previous results concerning relatedness, selection gradients and disruptive selection coefficients for lottery models (in particular those of Pen, 2000; Rousset and Ronce, 2004; Ohtsuki, 2010; Lehmann and Rousset, 2010; Rodrigues and Gardner, 2012; Wakano and Lehmann, 2014; Svardal et al., 2015; Mullon et al., 2016;

Parvinen et al., 2018, see sections G, H.3 and I.3 in the Supplementary Material for details), which confirms the validity of our approach. Finally, as a fully worked example, we investigate the evolution of adaptive polymorphism due to local adaption by extending the soft selection model of Svardal et al. (2015) to finite group size and hard selection. We confirm that adaptive polymorphism is generally favored by limited migration under soft selection and that small group size does not change this result qualitatively but tends to inhibit disruptive selection. For hard selection, however, the situation is more complicated as limited dispersal and finite group size favors not only disruptive selection but also repelling generalist strategies so that it becomes less likely that polymorphism can emerge from gradual evolution (Figure 3). With respect to limited migration this finding is also described by Débarre and Gandon (2011).

While our model allows for many different types of interactions between individuals within groups, it also has several limitations. At the individual level, we consider only scalar traits, but multidimensional (or functional-valued) traits can be taken into account by replacing derivatives by directional derivatives, which will not change the structure of our perturbation analysis. At the group level, we do not consider heterogeneity within groups, but in natural populations individuals within groups are likely to differ in their physiological state such as age, size and sex. To incorporate physiological heterogeneity requires an extension of the state space 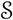 and to take into account the distribution of mutants within sub-groups of individuals belonging to the same physiological state in a group. The structure of our perturbation analysis, however, will remain unchanged by adding within-group heterogeneity, and only additional reproductive values and relatednesses will be needed. Likewise, in order to take isolation-by-distance into account, one again needs to extend the state space 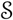, while to include diploidy one needs to extend the number of genetic states and this should only impact the relatedness coefficients. While such extensions remain to be done (and have all been done for the selection gradient *ρ*^(1)^ (*e.g.* Rousset, 2004)), they are unlikely to change the required components of the disruptive selection coefficient *ρ*^(2)^ and how they are connected algebraically. We thus conjecture that the representation of *ρ*^(2)^ holds generally.

In conclusion, for a large class of models we describe the consequences of limited dispersal and finite group size on evolutionary stability and diversification in heterogeneous populations, which we hope will help to formulate and analyze concrete biological models.

## Acknowledgments

HO and KP received support from the SOKENDAI Advanced Sciences Synergy Program (SASSP). HO and JYW received support from JSPS KAKENHI (No.16K07524). HO received support from JSPS KAKENHI (No.20K06812). JYW received support from JSPS KAKENHI (No.16K05283 and 16H06412). We are grateful to Dr. Yu Uchiumi for discussions.

## Supplementary Material

### A Mathematical properties of the baseline model

In this section, we provide a mathematical description of the stochastic process underlying the mutant dynamics that we consider in our paper.

#### A.1 Multitype branching process

We study the process of invasion of a mutant arising as a single copy (or a finite number of copies) in a monomorphic resident population and consider mutant dynamics as long as the mutant remains rare. Specifically, we pay attention to the number of groups including at least one adult mutant. Groups can differ in their state 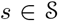 and in the number *i ∈* {1, *· · ·, n_s_*} of mutants. We therefore count the number of each “type” of group, where a type, denoted by *τ* here and thereafter, is specified by the vector *τ* = (*s, i*). The set of all possible types is 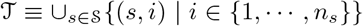 and there are 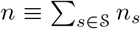 different group types.

A “population state” describes how the mutant is distributed among group types as long as it is rare in a population otherwise monomorphic for the resident type. It is specified by a vector ****M**** = {*M_τ}_ ∈* ℕ^*n*^, where ℕ = {0, 1, 2, *· · ·*} and where each *M_τ_* represents the number of type-*τ* groups at a given time. We consider that the change in state is given by a discrete-time and time-homogeneous Markov chain, denoted by 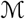, defined on N^*n*^, where the transition probability from population state ****M**** to ***M** ′* is given by *P* (***M*** → **M** ′). Here, *P* implicitly depends both on the mutant and resident trait values, *x* and *y*, respectively, and allows to define the generating function

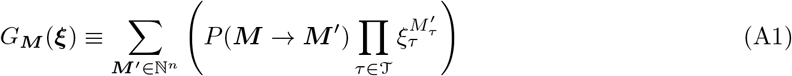

induced by the Markov chain, where 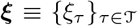 is a vector of dummy variables. This Markov chain has one absorbing state, which is the extinction of the mutant, and otherwise only transient states with the possibility that the absorbing state is never reached and so the number of mutants grows without bound.

We assume that the Markov chain 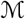 is a multitype branching process, meaning that each group “behaves” independently of the other groups. Mathematically, this assumption is embodied by the generating function of the Markov chain (eq.(A1)) being given by

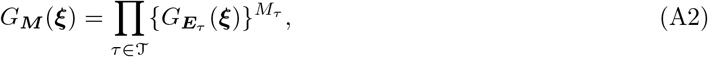

where ***E**_τ_* is a vector of length *n* whose *τ* -th component is 1 and all the others are zero. Intuitively speaking, eq.(A2) shows that *P* (***M** → **M** ′*) is uniquely determined by the “fundamental” transitions probabilities, 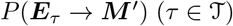, each representing how many groups of the same and different types are “produced” by a single type-*τ* group in the previous time step (by “produced” we mean that the survival and reproduction of individuals in a single type-*τ* group affect the composition of that group in the descendant generation, as well as the composition of other groups by emigration of offspring, *e.g.*, eq.(1) of Lehmann et al., 2016).

For a given initial population state ****M****_0_ *∈* ℕ^*n*^, the ultimate extinction probability of mutants is defined as

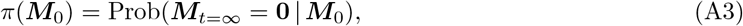

where ***M***_t=*∞*_ is the population state vector when the number of time steps *t → ∞*, and **0** represents a vector of zeroes of length *n*. Now we define

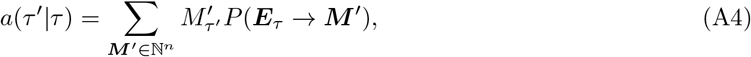

which is the expected number of type-*τ ′* groups that are “produced” by a single type-*τ* group^1^. We collect the expectations *a*(*τ ′|τ*) for all *τ, τ ′* to construct matrix ****A**** = {*a*(*τ ′|τ*)}. We assume that (i) matrix ****A**** is primitive, as specified in section 2.2 in the main text, and that (ii) 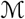 is not “singular” (where 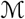 being “singular” means that all of 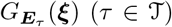 are linear functions without a constant term; see Harris, 1963). Let *ρ* be the largest eigenvalue of ****A**** (since ****A**** is primitive it follows from the Perron-Frobenius theorem that such a unique positive *ρ* exists). Then, a standard result in multitype branching process theory (Harris, 1963; Karlin and Taylor, 1975) guarantees that the following relations hold between *ρ* and the *π*(***E***_*τ*_):

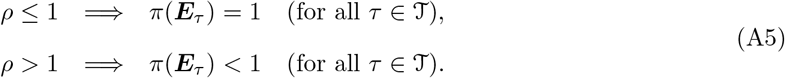

It is precisely matrix ****A**** = {*a*(*τ ′|τ*)} = {*a*(*s′, i′|s, i*)} that we study in the main text. Because our focus is only on the uninvadability of the resident with respect to invasion of single mutants, which translates to whether or not *π*(****E****_(*s,*1)_) = 1, we do not have to distinguish two different multitype branching processes that yield the same ****A****-matrix. Therefore, an evolutionary invasion analysis can be started from ****A****, and does not need to detail the transition probabilities of the underlying multitype branching process.

#### A.2 List of assumptions and their implications

We here summarize the basic mathematical assumptions for our model and their biological implications.

i. A time-homogeneous Markov chain 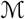, which depends on *x, y* and whose state space is 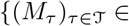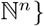, is a multitype branching process. This implies that each group that includes at least one mutant behaves independently of all other groups with mutants.
ii. Matrix ****A**** = {*a*(*τ ′|τ*)}, calculated from 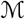 as in eq.(A4), is primitive. This implies that each type of group has a positive contribution to the production of any other type of group after some finite number of time steps.
iii. 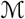 is not singular. Hence, we do not consider a degenerate multitype branching process in which each group always produces exactly one group at the next time step.
iv. Individual fitness 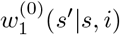 as defined in eq.(14) does not depend on *i* (and therefore can be written as 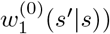. This implies that resident and mutant individuals are indistinguishable (and exchangeable) under neutrality.
v. Matrix ****W**** ^(0)^, whose entries are determined by 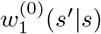, has 1 as its largest eigenvalue. This implies that a monomorphic population of resident individuals stays at the same average group size due to density-dependent regulation.

### B Derivation of perturbations of invasion fitness

We here prove eq.(21) and eq.(22). Before doing so, we list some frequently used relations:

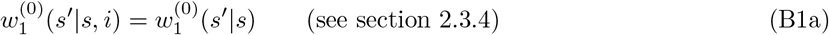

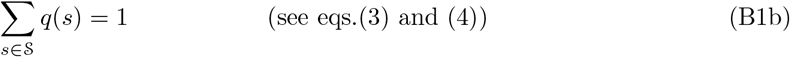

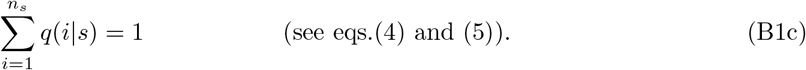

We decompose *w*_1_(*s′|s, i*) in eq.(16a) into a neutral part and a non-neutral part,

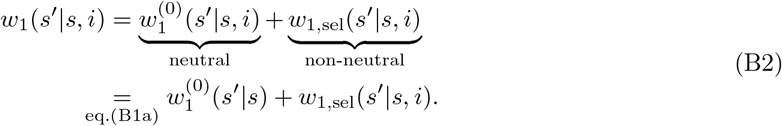

Note that, by definition, the *C*-th order perturbation of *w*_1,sel_(*s′|s, i*) with respect to *δ* equals

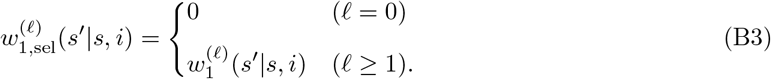

From eq.(16a), we then have

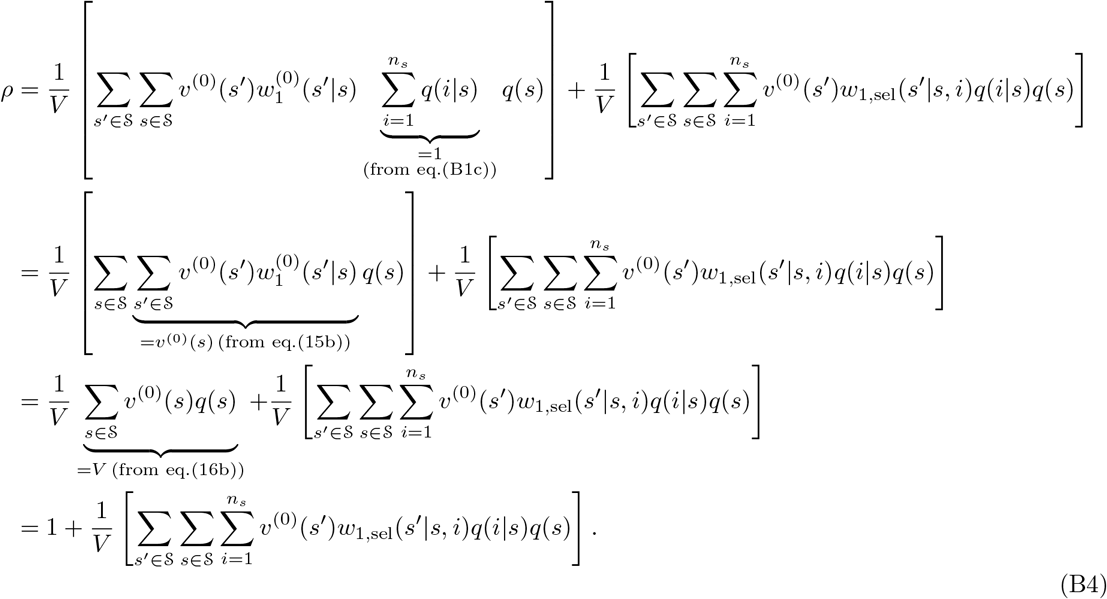

As a check, the zeroth order perturbation is

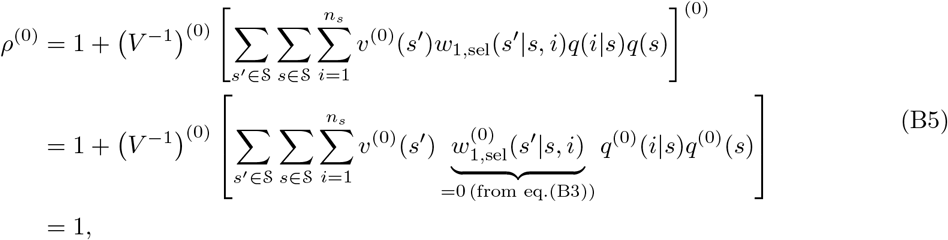

as expected. The first-order perturbation of eq.(B4) is

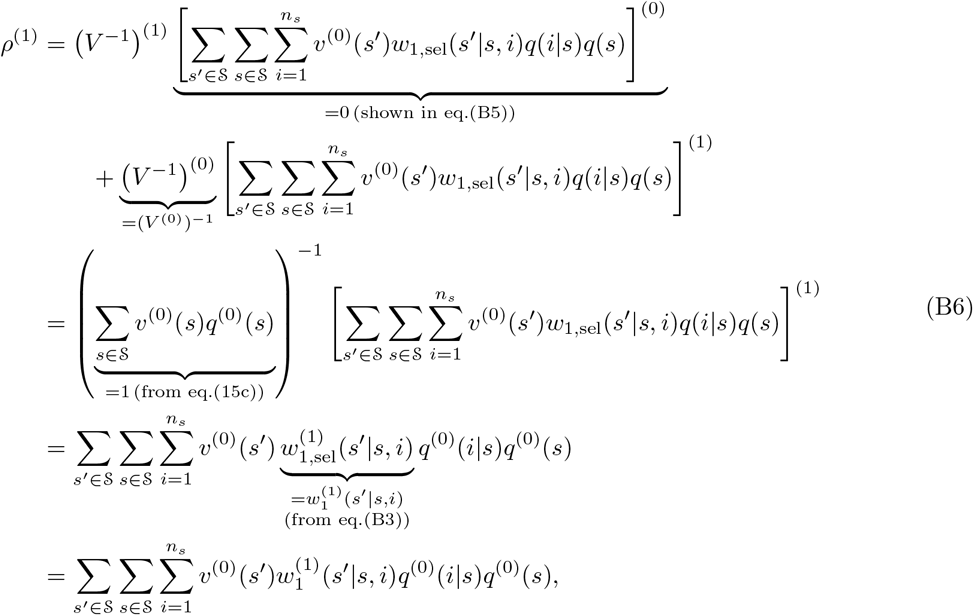

which reproduces eq.(E.14) in Lehmann et al. (2016). Note that the first-order perturbation of the term in square brackets in the third line of eq.(B6) can potentially produce more terms in the fourth line, but they are null because 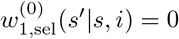 (see eq.(B3)). Equation (B6) proves eq.(21) in the main text.

Next, we study *ρ*^(2)^ under the condition *ρ*^(1)^ = 0. The second-order perturbation of eq.(B4) is

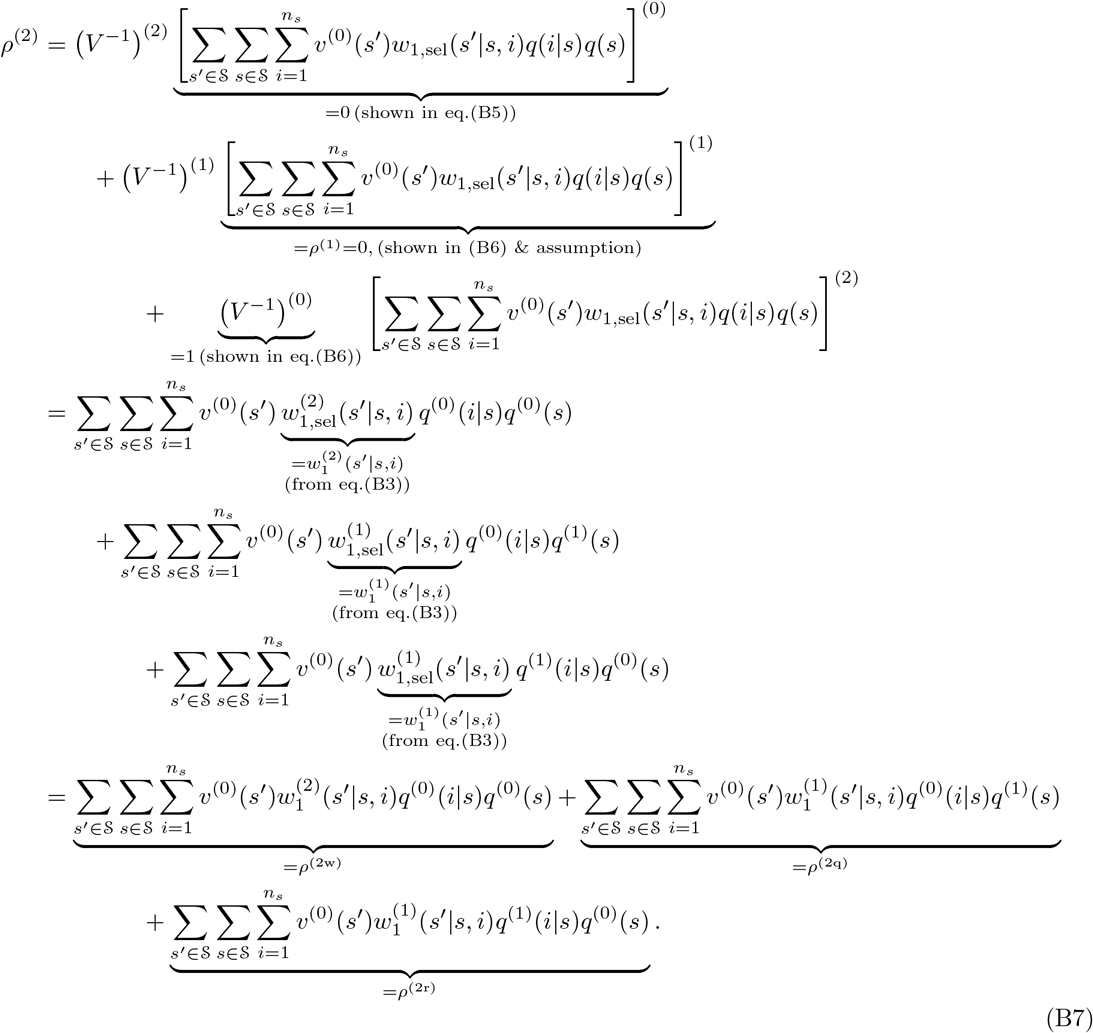

Note that the second-order perturbation of the term in square brackets in the third line of eq.(B7) can potentially produce more terms in the following lines, but they are null because 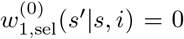 (see eq.(B3)). Equation (B7) proves eq.(22) in the main text.

### C Derivation of recursions

In order to practically use the general formulae for the first and second order derivatives of invasion fitness, eq.(21) and eq.(22), or to use the corresponding formulae derived for the individual fitness functions, eq.(32) and eq.(34), we need to know *q*(*s*) (the asymptotic distribution that a randomly sampled mutant finds itself in a group of state *s*) and *q*(*i|s*) (the asymptotic distribution that a randomly sampled mutant, given that it is sampled from a group in state *s*, finds itself in a group with *i* mutants) under neutrality as well as the first-order perturbation of these quantities with respect to *δ*. Due to eq.(7), knowing *q*(*i|s*) (*i* = 1, *· · ·, n_s_*) is equivalent to knowing relatedness, *r_k_*(*s*) (*k* = 1, *· · ·, n_s_*). The purpose of this section is to derive recursions that *q*(*s*) and *r_k_*(*s*) satisfy. Specifically, in section C.1 we derive recursions that are valid for any *δ*, and in section C.2 we describe their perturbations to the zeroth-(hence under neutrality) and first-order of *δ*.

#### C.1 Recursions of *q*(*s*) and *r_k_*(*s*) for arbitrary *δ*

For simplicity, we will from here on omit the lower and upper bound of the summation when obvious from the context.

##### C.1.1 Recursion for *q*(*s*)

Writing *ρ****u*** = ****Au**** component-wise gives

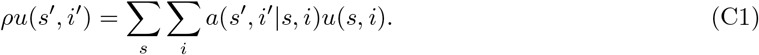

By multiplying both sides with *i′* we obtain

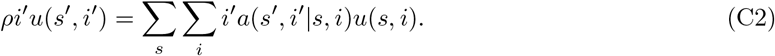

Summing eq.(C2) over *s′* and *i′* gives

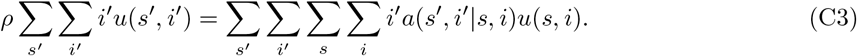

Dividing eq.(C2) by eq.(C3) results in

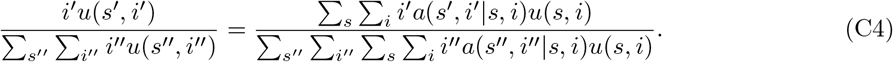

Using eq.(3) we note that the left-hand side equals *q*(*s′, i′*). Thus, eq.(C4) can be rewritten as

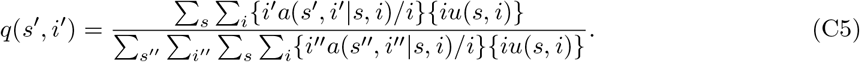

We divide both the numerator and the denominator of the right-hand side of eq.(C5) by the constant, ∑_*s*′_ ∑_*i*′_ *iIu*(*s′, i′*). By using eq.(3) once again we obtain

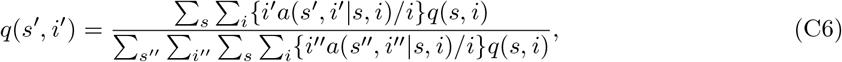

which is the recursion that *q*(*s, i*) obeys.

To obtain the recursion that *q*(*s*) obeys, we sum eq.(C6) over *i′* and obtain

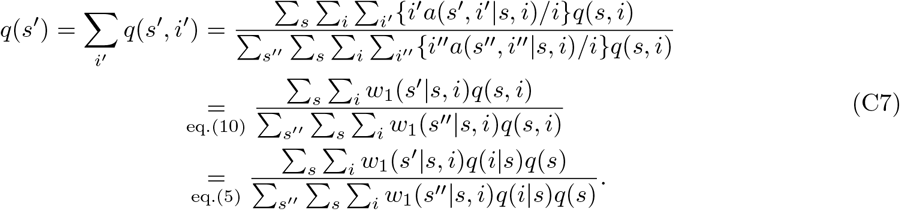

###### Interpretation of *ρ*

Dividing eq.(C3) by the constant ∑_*s*_ ∑_*i*_ *iu*(*s, i*) gives

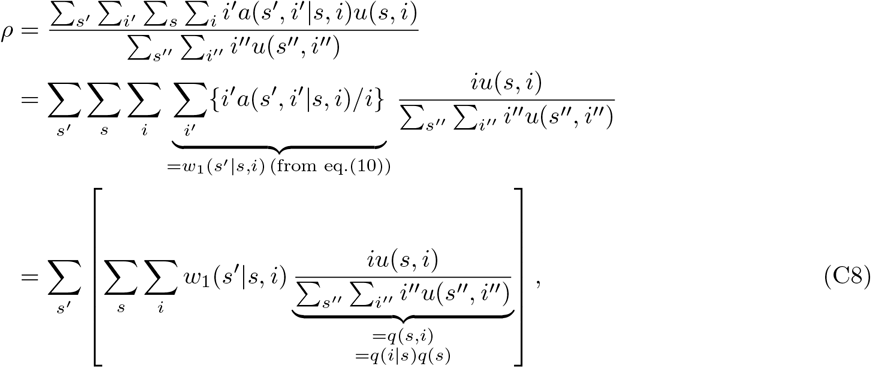

which reproduces eq.(5) of Lehmann et al. (2016). The term inside the square bracket of eq.(C8) can be interpreted as the state-*s′* component of the expected individual fitness of a mutant randomly sampled from the asymptotic distribution *u*(*s, i*) (*i.e.*, the probability for a randomly sampled mutant to find itself in an (*s, i*)-group is proportional to *iu*(*s, i*)). Thus, invasion fitness *ρ* can be interpreted as the expected number of mutant copies produced by a lineage member randomly sampled from the distribution *q*(*s, i*).

Combining eqs.(C7) and (C8) gives us a useful relationship

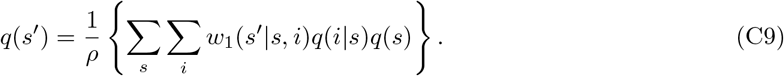

##### C.1.2 Recursion for *r_k_*(*s*)

From the definition of *r*_1_ in eq.(7) we have *r*_1_(*s*) = 1. Thus, we are interested in the recursions for *r_k_*(*s*) for *k ≥* 2. Using the definition for *q*(*s, i*) in eq.(5) and the expression eq.(C6), we have

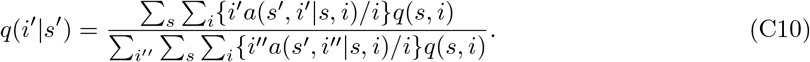

Multiplying both sides of the last equation by *φ_k_*(*s′, i′*) and summing over *i′* gives

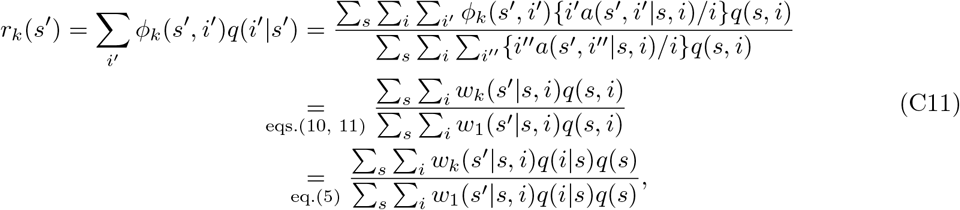

where *w_k_* denotes *k*-fitness as defined in eq.(11) in the main text.

#### C.2 Perturbations of *q*(*s*) and *r_k_*(*s*)

Here, we derive recursions satisfied by the zeroth- and first-order perturbations of *q*(*s*) and *r_k_*(*s*). These will be of practical use when computing the selection gradient and disruptive selection coefficient, *e.g.*, eq.(21) and eq.(22) or eq.(32) and eq.(34). We also show that the recursions for the perturbation of *r_k_*(*s*) can be greatly simplified if we make some additional assumptions on the fitness functions.

##### C.2.1 Perturbation of *q*(*s*)

The following observation is useful in later calculations. For the perturbations of eqs.(B1b) and (B1c) we obtain

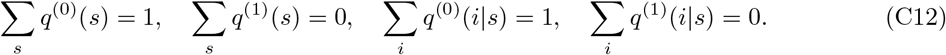

###### Zeroth-order perturbation of *q*(*s*)

The zeroth-order perturbation of eq.(C9) with respect to *δ* is given by

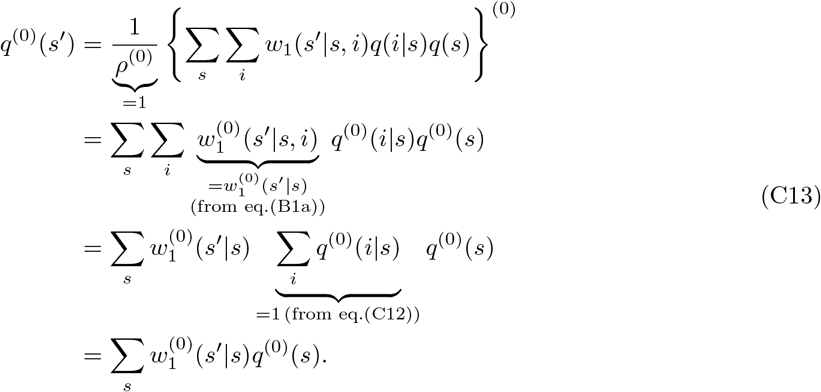

This proves eq.(15a) in the main text.

###### First-order perturbation of *q*(*s*)

Assuming *ρ*^(1)^ = 0 and using the quotient rule the first-order perturbation of eq.(C9) with respect to *δ* is given by

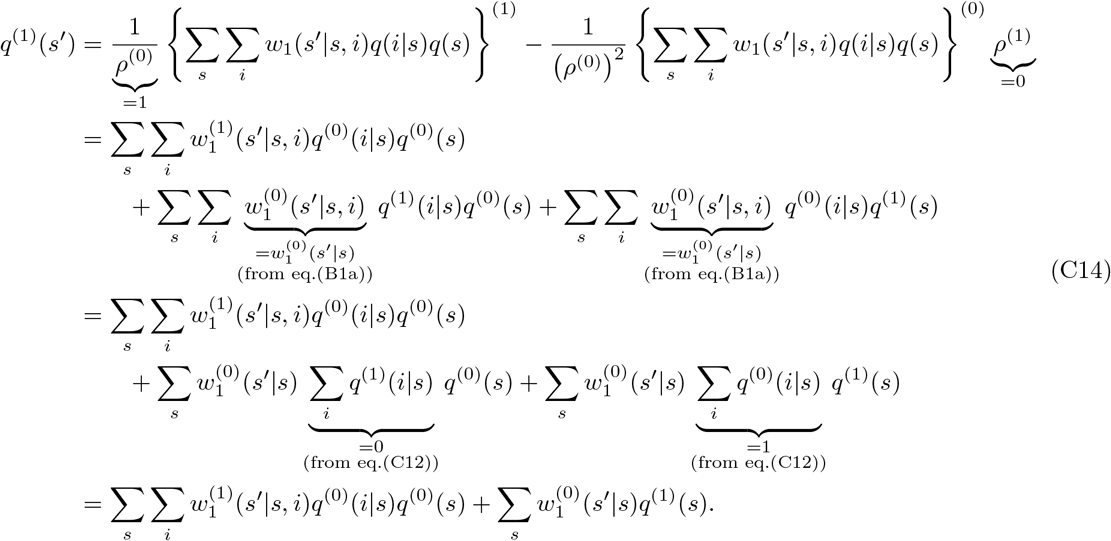

##### C.2.2 Perturbation of *r_k_*(*s*)

###### Zeroth-order perturbation of *r_k_*(*s*)

With respect to the zeroth-order perturbation of eq.(C11) with respect to *δ* we obtain

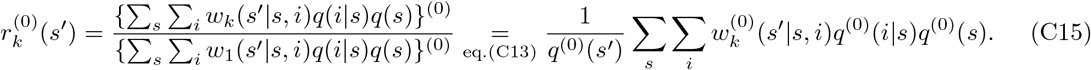

Of practical importance for *k ≥* 2 is the case that 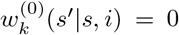 holds for all *s′ ≠ s* and all *i* = 1, *· · ·, n_s_*. This applies, for example, when the state of a given group does not change and mutants settle in new groups only as single individuals (no propagule dispersal). Then eq.(C15) simplifies to

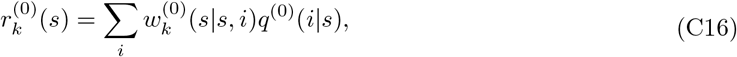

and 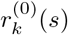 can be calculated independently of 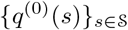.

###### First-order perturbation of *r_k_*(*s*)

Assuming *ρ*^(1)^ = 0 and using the quotient rule the first-order perturbation of eq.(C11) with respect to *δ* equals

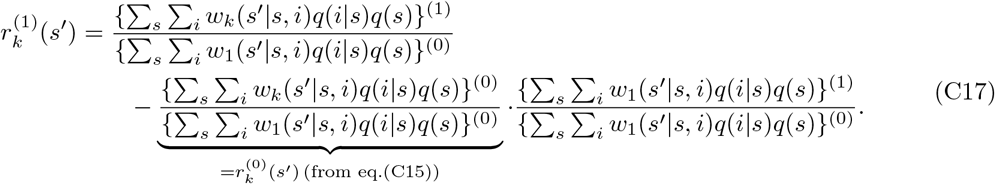

For *ρ*^(1)^ = 0 we observed in eq.(C14) that {∑*_s_ ∑_i_ w*_1_(*s′|s, i*)*q*(*i|s*)*q*(*s*)}^(1)^ = *q*^(1)^(*s′*) holds. Applying this relation and eq.(C13) we obtain

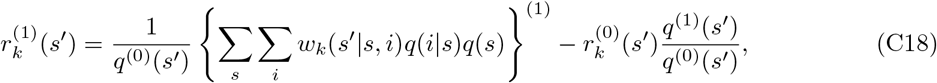

which, upon expanding the first-order perturbation on the right hand side more explicitly, becomes

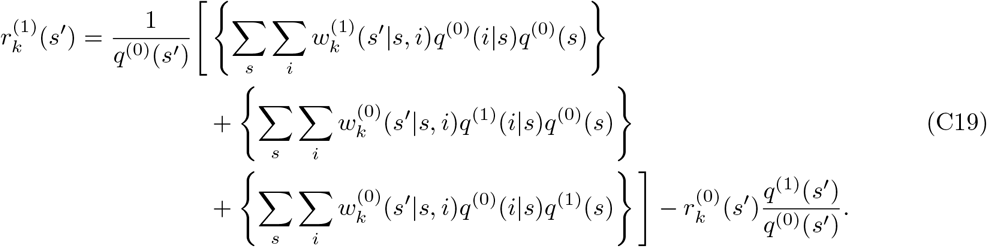

When considering the case that 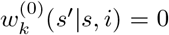 holds for all *s′* ≠ *s* and all *i* = 1, *· · ·, n_s_*, just as we did for the case of the zeroth-order perturbation, eq.(C19) simplifies to

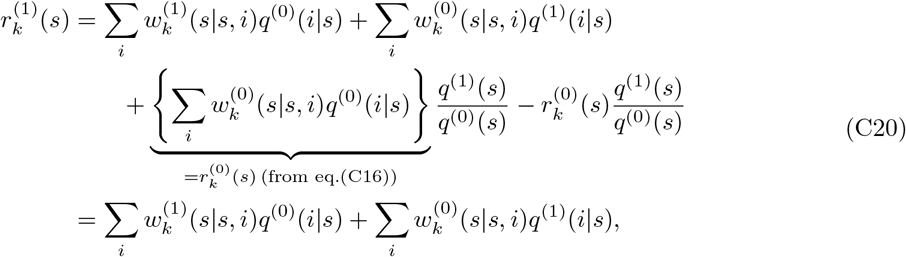

and 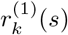 can be calculated independently of 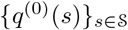 and 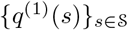.

### C.3 Closedness of the recursions in *q*(*s*) and *r_k_*(*s*)

In the previous sections, we obtained recursions for *q*(*s*), *r_k_*(*s*) and the perturbations thereof to the first order. However, it is not clear whether these actually form a closed system of equations in terms of the variables. The purpose of this section is to show that eqs.(C7) and (C11) (and the perturbation thereof) indeed constitute such a closed system of equations. To do this, we pay attention to the sum ∑_*i*_ *w*_*k*_(*s′|s, i*)*q*(*i|s*), which frequently appears in eqs.(C7) and (C11). We prove that this sum can be written as a linear combination of the *n_s_* relatedness coefficients 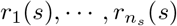. With this, it follows that the desired result indeed holds.

**Proof:** For fixed *k, s′* and *s* (and for fixed *x* and *y*), consider *n_s_* distinct points on a two-dimensional plane, (*i, w_k_*(*s′|s, i*)) *∈* ℝ^2^ (*i* = 1, *…, n_s_*). Then, from a standard result of polynomial interpolation, there exists a unique polynomial in *ξ*, denoted by *L_k,s′,s_*(*ξ*) (called a Lagrange polynomial), whose graph {(*ξ, L_k,s′,s_*(*ξ*)) *| ξ ∈* ℝ} passes through the above *n_s_* points and whose order as a polynomial in *ξ* is equal to or less than *n_s_ −* 1. Take such a polynomial *L_k,s′,s_*. By definition

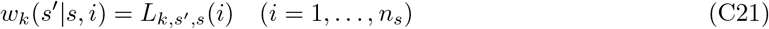

holds. Now we define a set of polynomials in *ξ*, {Φ_1,*s*_(*ξ*), *· · ·,* Φ_*ns,s*_(*ξ*)}, as

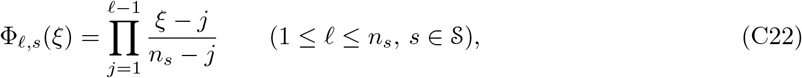

where we define Φ_1,*s*_(*ξ*) = 1. This set of *n_s_* polynomials of order 0 to *n_s_ −*1 can be written more explicitly as

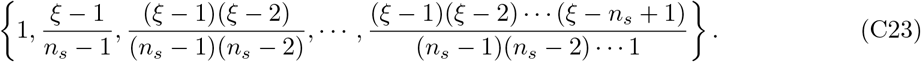

It is thus a basis of the vector space composed of all polynomials in *ξ* of order equal to or less than *n_s_ −* 1. Because *L_k,s′,s_*(*ξ*) is one such polynomial it can be written as

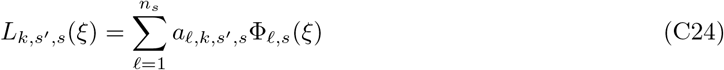

for some *a_ℓ,k,s′,s_ ∈* ℝ (*ℓ* = 1, *· · ·, n_s_*). By construction

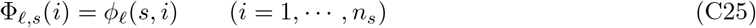

holds (compare eq.(C22) with eq.(6)).

We now consider ∑_*i*_ *wk*(*s′|s, i*)*q*(*i|s*). Using *Lk,s′*_*,s*_ and eqs.(C21),(C24) and (C25) we obtain

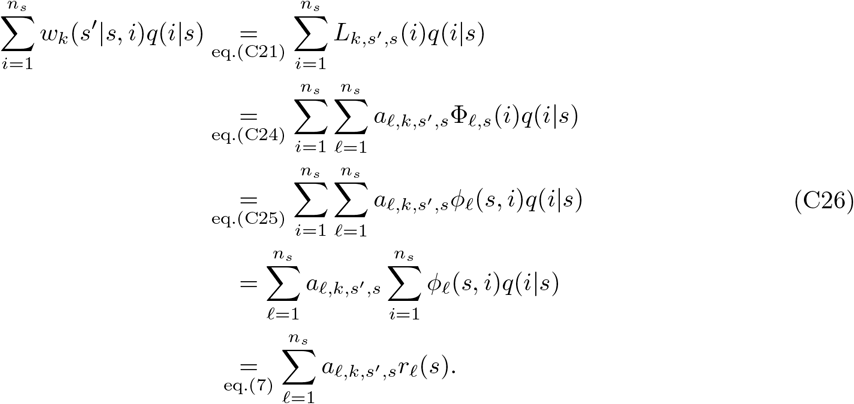

Given *r*_1_(*s*) = 1 for all 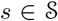, eqs.(C7) and (C11) form a large but closed system of equations with *q*(*s*) and *r_k_*(*s*) for all *k* = 2, *· · ·, n_s_* and all 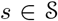. Its size is 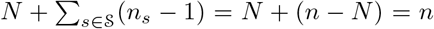, which makes sense since *n* is the dimension of matrix ****A****.

### D Individual 3-fitness

We define the following four different individual 3-fitness functions. For that purpose, consider a focal individual in a group in state *s* who adopts *z*_1_, and its *n_s_ −* 1 neighbors who adopt ***z***_*−{1*}_ in an otherwise monomorphic population for *z*.

#### Individual 3-fitness of type I

Define

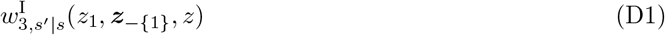

as the expected number of offspring in state *s′* that descend from the focal individual adopting *z*_1_ in state *s* and that have two random neighbors (sampled without replacement) that both descend from the focal individual.

#### Individual 3-fitness of type II

Consider one of the focal’s neighbor, called the target individual, who adopts *z*_2_. We define

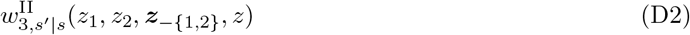

as the expected number of offspring in state *s′* that descend from the focal individual adopting *z*_1_ in state *s* and whose two random neighbors (sampled without replacement) both descend from the target individual (adopting *z*_2_).

#### Individual 3-fitness of type II’

Consider one of the focal’s neighbor, called the target individual, who adopts *z*_2_. We define

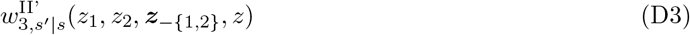

as the expected number of offspring in state *s′* that descend from the focal individual adopting *z*_1_ in state *s*, and where one of its two random neighbors (sampled without replacement) descends from the focal individual while the other descends from the target individual (adopting trait value *z*_2_). The following useful symmetry between 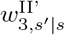 and 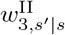 holds:

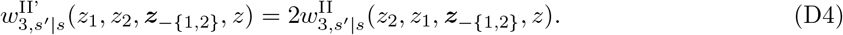

The reason for this symmetry is as follows. Consider a focal individual adopting *z*_1_ and a target individual adopting *z*_2_ in the same state-*s* group. Suppose that a group in state *s′* at the next time step comprises *A*_1_ individuals that descend from the focal individual and *A*_2_ individuals that descend from the target individual. Then, by definition such a group contributes to the 3-fitness of type II’ of the focal individual 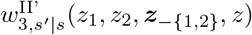 by

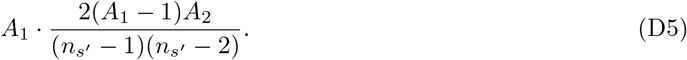

The same group contributes to the 3-fitness of type II of the target individual, 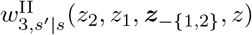 by

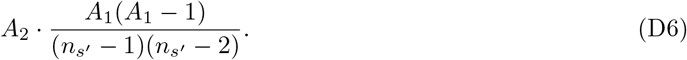

Equation (D5) is twice as large as eq.(D6), and therefore the symmetry eq.(D4) holds.

#### Individual 3-fitness of type III

Consider two neighbors of the focal individual, the one who adopts *z*_2_ (called the first target individual) and the one who adopts *z*_3_ (called the second target individual). Then we define

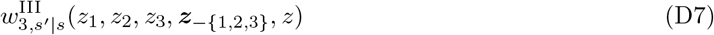

as the expected number of offspring in state *s′* that descend from the focal individual adopting *z*_1_ in state *s*, and where one of its two random neighbors (sampled without replacement) descends from the first target individual (adopting *z*_2_) and the other descends from the second target individual (adopting *z*_3_). The following symmetry exists for a similar reason as above. For any permutation of *σ* of the set {1, 2, 3} holds

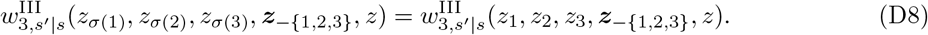

#### Calculation of *w*_3_

With these four individual 3-fitness functions the individual 3-fitness of a mutant in an (*s, i*)-group, for 3 *≤ i ≤ n_s_*, can be written as

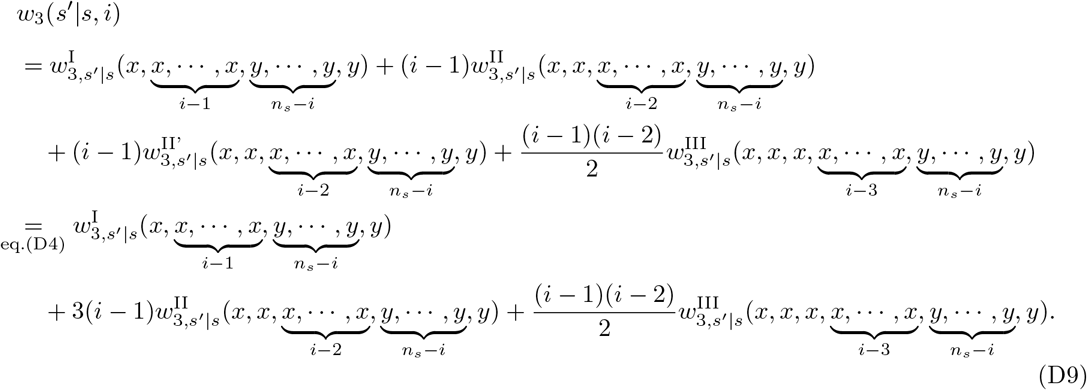

Its zeroth-order perturbation with respect to *δ* is

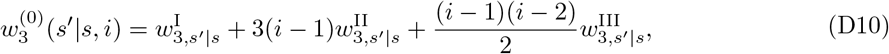

where all 3-fitness functions *w*_3_ are evaluated at (*y, · · ·, y*). Equation (D10) is the expected number of offspring in state *s′* under neutrality that descend from a focal mutant individual in state *s* and have two random neighbors (sampled without replacement) that are both mutants.

Note that the derivation above assumed 3 *≤ i ≤ n_s_*, but we can separately confirm that eq.(D10) is valid for any 1 *≤ i ≤ n_s_*, because some ill-defined terms for *n_s_* = 1 and 2 become nullified by the factors *i −* 1 and *i −* 2.

### E Derivation of the quantities with individual fitness functions

Here, we derive all the key results presented in section 3.3 in the main text.

#### Equation (33)

By setting *k* = 2 in eq.(C15) and substituting 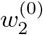 with eq.(29a), we obtain

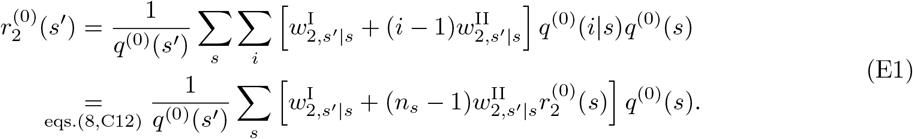

This proves eq.(33).

If 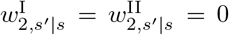 for *s′* ≠ *s* (this is the case, for example, when propagule dispersal is not allowed), we can use eq.(C16) instead of eq.(C15). Substituting eq.(29a) in eq.(C16) for *k* = 2 gives

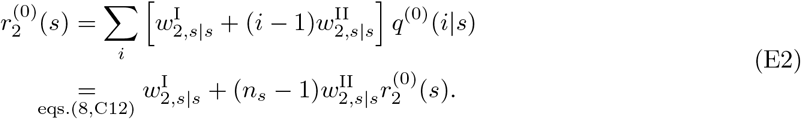

#### Equation (34)

First, substituting 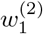 in eq.(22b) with eq.(25c) gives

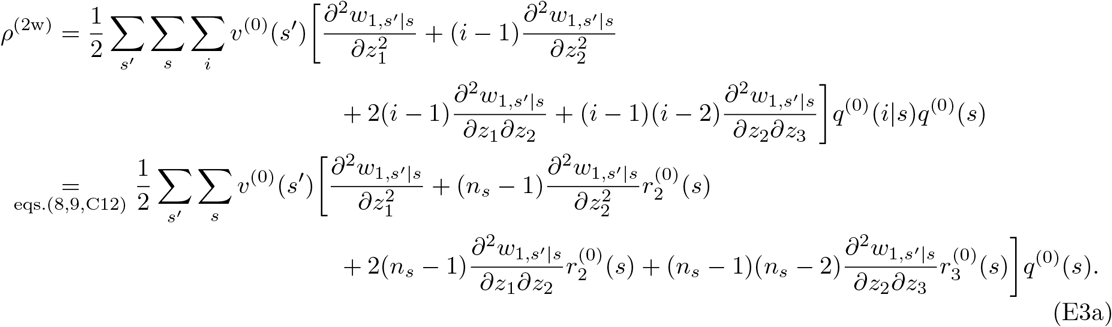

Second, substituting 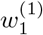 in eq.(22c) with eq.(25b) gives

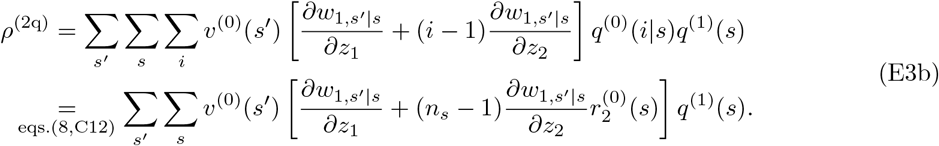

Third, substituting 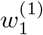 in eq.(22d) with eq.(25b) gives

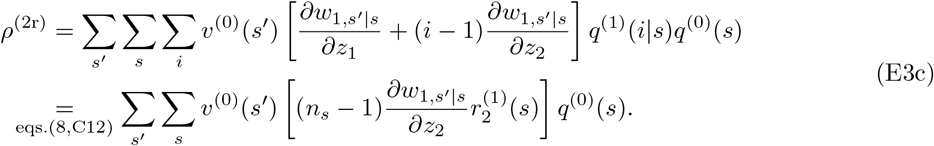

#### Equation (35)

Substituting 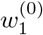 and 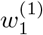 in eq.(C14) with eq.(25a) and eq.(25b), respectively, gives

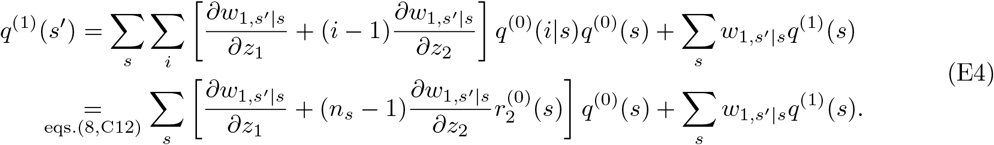

#### Equation (36)

By setting *k* = 3 in eq.(C15) and substituting 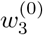 with eq.(30) we obtain

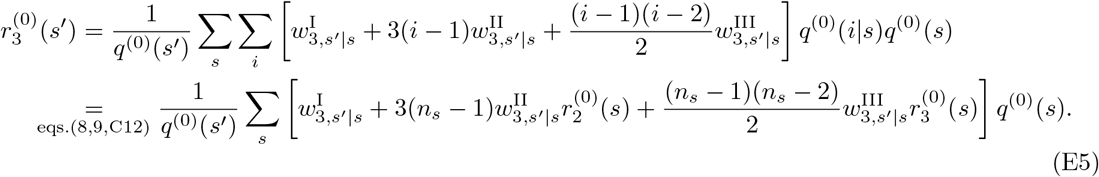

This proves eq.(36) in the main text.

If 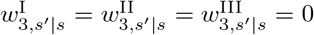 for *s′* ≠ *s* (this is the case, for example, when propagule dispersal is not allowed), we can use eq.(C16) instead of eq.(C15). Substituting eq.(30) in eq.(C16) for *k* = 3 then gives us

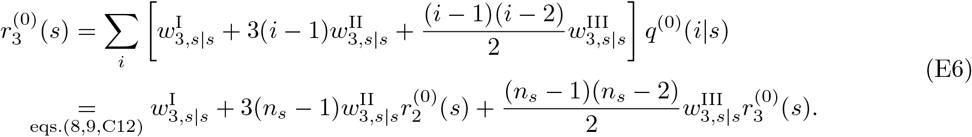

#### Equation (37)

By setting *k* = 2 in eq.(C19), substituting 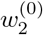 and 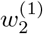 with eq.(29a) and (29b), respectively, we obtain

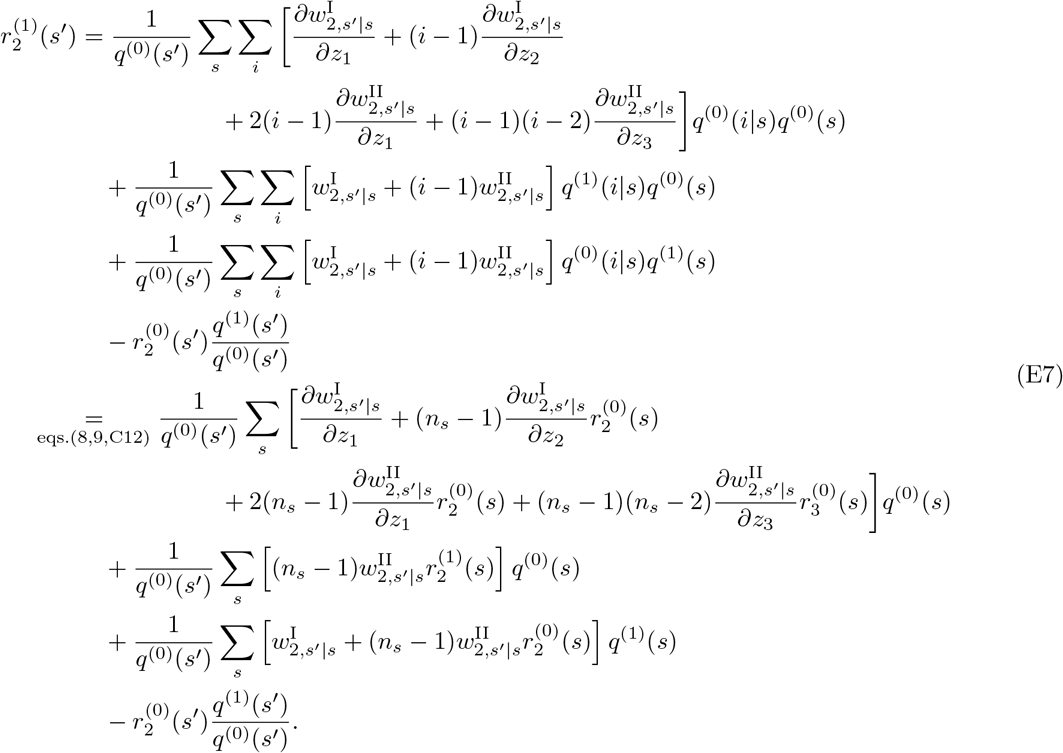

This proves eq.(37) in the main text.

If 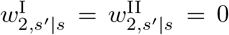 for *s′* ≠ *s* (this is the case, for example, when propagule dispersal is not allowed), we can use eq.(C20) instead of eq.(C19). Substituting eq.(29) in eq.(C20) for *k* = 2 gives us

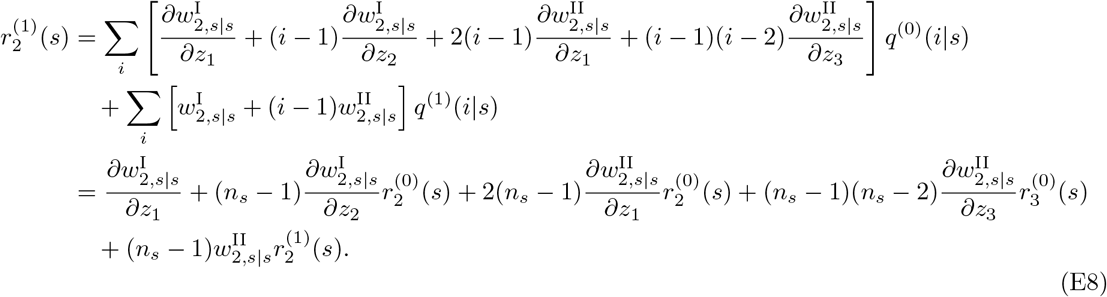

Therefore, 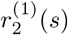 is solved as

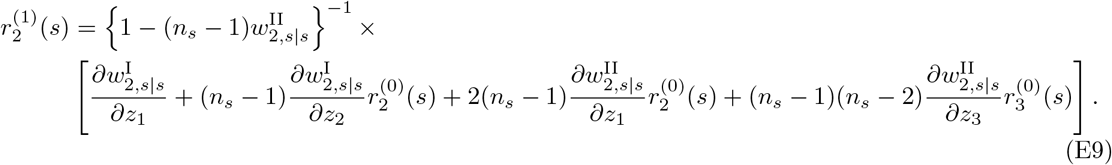

From eq.(E2) we see that 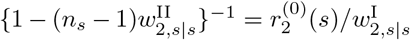 holds, so eq.(E9) can also be written as

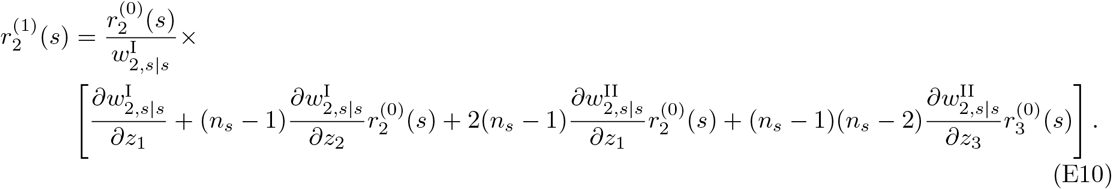

### F Derivation of the results for the lottery model

Here, we derive all the results presented in section 4.1 of the main text.

#### F.1 Calculations for *q*(*s*) and *v*^(0)^(*s*)

Under the assumptions (i)-(iii) described in section 4.1 of the main text, eq.(38c) therein can be further decomposed as

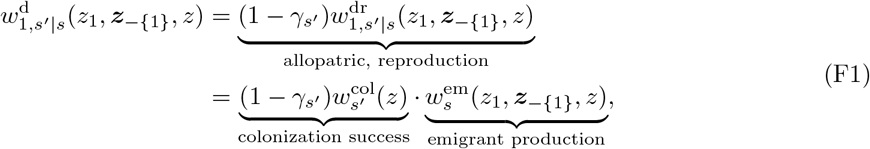

where 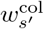 represents a component of colonization success of dispersing offspring arriving in groups in state 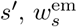 represents the emigrant production in groups in state *s*, and 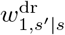 is given as a product of these two terms. This multiplicative decomposition is a key property that greatly simplifies the following analysis. It follows from the fact that when dispersal occurs individually and independently to a random destination, the production of emigrants in a group in habitat *s* does not depend on the habitat *s′* of the destination group and the colonization success depends only on the habitat of the destination group and the resident trait value.

Before proceeding to the derivation, we show that

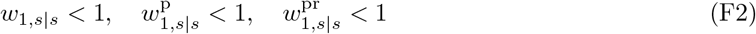

hold for all 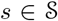, because we will frequently use these facts without a particular notice below. The proof starts from the observation that eq.(15a) is rewritten as

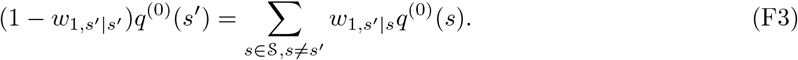

Remember that *q*^(0)^(*s*) *>* 0 holds for all 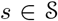, and therefore the right hand side of eq.(F3) is non-negative. Suppose the right hand side of eq.(F3) is zero. Then we have *w*_1,*s*′*|s*_ = 0 for all *s* ≠ *s′*, which contradicts that matrix ****W**** ^(0)^ is primitive (see section 2.3.4). Therefore, the right hand side of eq.(F3) must be strictly positive. Because *q*^(0)^(*s′*) *>* 0, it follows that *w*_1,*s*′|s_′ < 1 holds, and this argument is valid for all 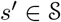. Second, because 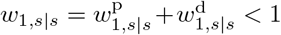, we have 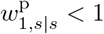. Third, from eq.(38b), one can see that the relation

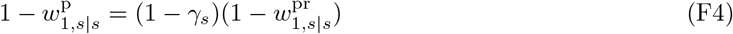

holds. Because we have just proven that 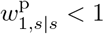 and because *γ_s_ <* 1, one concludes that 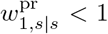 holds. End of the proof.

We now first calculate explicitly *q*^(0)^(*s*) and *v*^(0)^(*s*). For that purpose, the vector-matrix notation in eqs.(15a) and (15b) is helpful. In fact, from eqs.(38) and (F1), the fitness under neutrality is written as

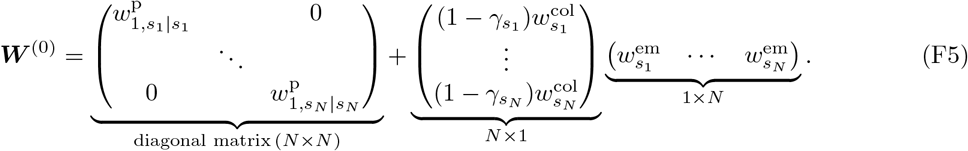

Note that 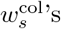 and 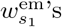 without variables are evaluated respectively at *y* and (*y, · · ·, y*) here and thereafter. To solve eq.(15a), we right-multiply eq.(F5) by the column vector ****q****^(0)^ and obtain

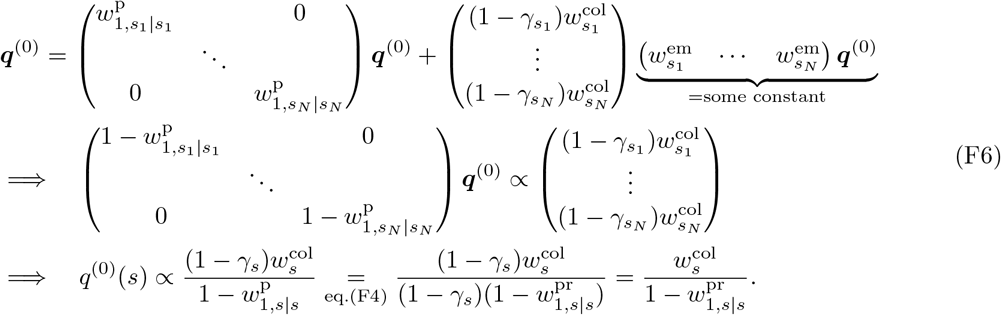

To solve eq.(15b) we left-multiply eq.(F5) by the row vector ****v****^(0)^ and obtain

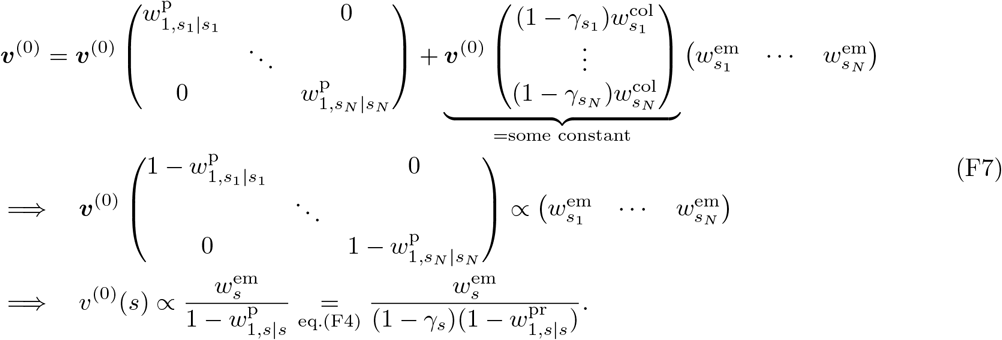

We normalize *q*^(0)^(*s*) and *v*^(0)^(*s*) to satisfy eq.(15c) and eq.(B1b) and obtain the following result:

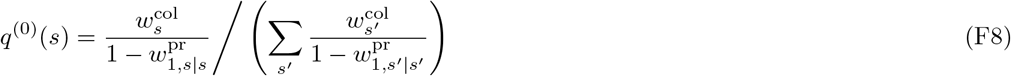

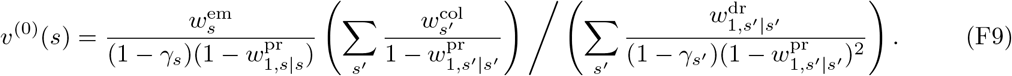

Combining eqs.(F8) and (F9) gives eq.(40) in the main text. In particular, their product

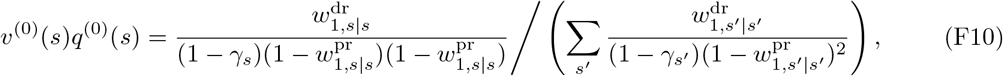

can be recognized as the the class reproductive value of a group in habitat *s* (Rousset, 2004).

We note that *q*^(0)^(*s*) can also be derived via a different pathway. To ease our understanding, let us temporarily consider a finite population that consists of *N*_G_ groups (later we will take the limit, *N*_G_ *→ ∞*, because we actually consider an infinitely large population in this paper) and consider the neutral case. The total number of individuals in groups in habitat *s′* in a next time step is the sum of the number of individuals in habitat *s′* produced by individuals in habitat *s* over all 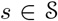, which is

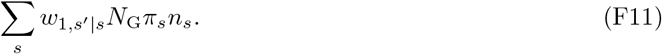

Here, *N*_G_*π_s_* is the total number of groups in habitat *s*, and *N*_G_*π_s_n_s_* is the total number of individuals in groups in habitat *s*. However, eq.(F11) can also be written as *N*_G_*π_s′_ n_s′_*, so equating these two quantities cancels out *N*_G_ and gives us

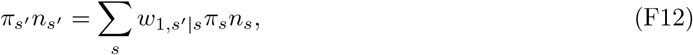

and therefore eq.(F12) is valid for *N*_G_ *→ ∞* as well (see also eq.(E.21) of Lehmann et al., 2016). Equation (F12) suggests that the column vector 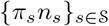 is an eigenvector of matrix ****W**** ^(0)^ corresponding to the leading eigenvalue of 1. From the Perron-Frobenius theorem the eigenspace of the matrix ****W**** ^(0)^ corresponding to the leading eigenvalue 1 is one-dimensional. Thus, from the uniqueness of the normalized eigenvector we obtain

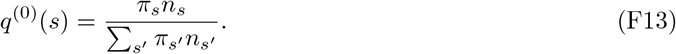

Note that eq.(F8) and eq.(F13) are both correct. Their equivalence suggests the existence of the following constraint in the choice of 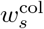 and 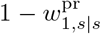 in our spatial lottery model, namely, that there exists a positive constant *C* which is independent of *s* and

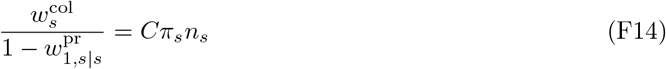

is satisfied for all 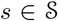.

Finally, we calculate *q*^(1)^(*s*) under the assumption that *ρ*^(1)^ = 0. From eq.(35), we then obtain

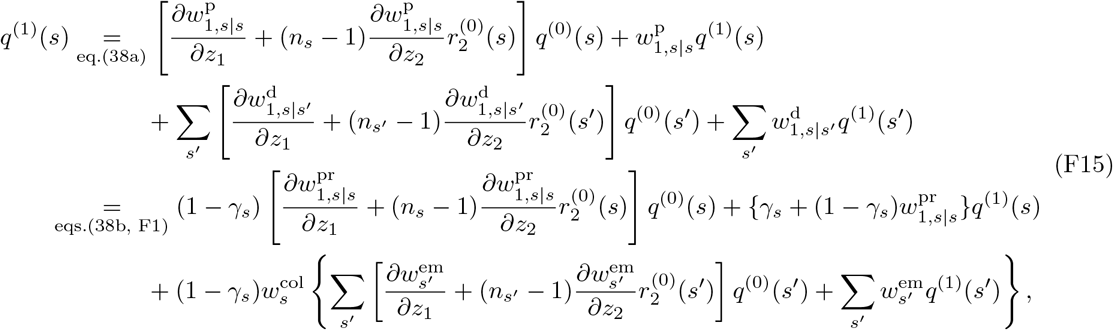

which is implicitly solved as

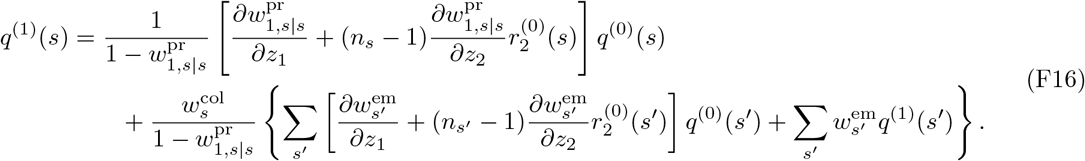

Using ∑ *q*^(1)^(*s*) = 0 (see eq.(C12)), we have

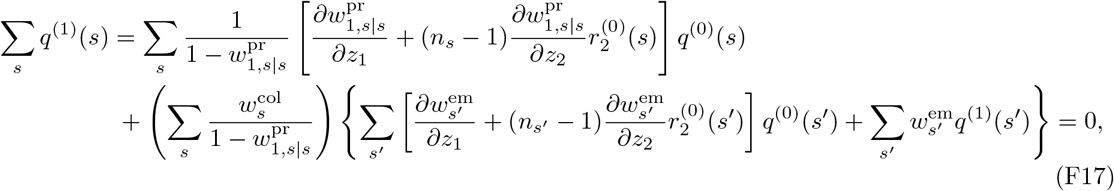

which shows that

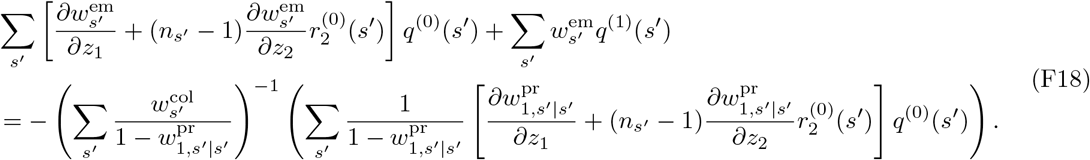

holds. Putting eq.(F18) back into eq.(F16) and using eq.(F8) gives us

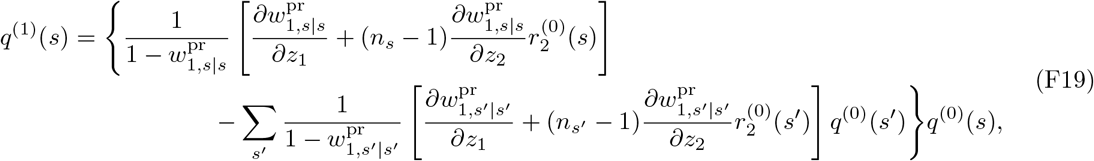

which proves eq.(43) in the main text.

#### F.2 Individual 2-fitness and 3-fitness for the lottery model and calculations for relatedness

The purpose of this subsection Is to derive pairwise relatedness and three-way relatedness under neutrality, as well as the first-order perturbation of pairwise relatedness under the assumption of the lottery model in section 4.1. For that purpose we will show that for this lottery model individual 2-fitness and individual 3-fitness are written in terms of the philopatric component of individual 1-fitness as in eq.(F23) and eq.(F31).

Let us label individuals in a focal group in habitat *s* from 1 to *n_s_*, and let their trait values be *z*_1_ to 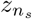, respectively. Let us also operationally define “positions” in this group, from “position 1” to “position *n_s_*”. If adult *k* survives (with occurs with probability *γ_s_*) we operationally assume that he/she occupies “position *k*” in the next generation. If adult *k* dies (which occurs with probability 1 *− γ_s_*) we assume that “position *k*” in the next generation becomes open to competition. In the latter case, adult *i* bears a descendant in this “position *k*” with probability 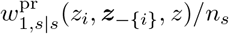, which we will write as *ω_i_* in what follows as a short-hand notation.

With these, the probability *ζ_i,k_* that an individual in position *k* in the next generation in the focal group “descends” from adult *i* in that group in the previous generation (thus including self through survival) is, for the lottery model, given by

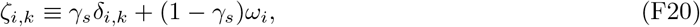

where *δ_i,k_* equals one if *i* = *k* and otherwise zero. In eq.(F20), the first term represents the probability that adult *i* survives and occupies position *k* (thus equal to *γ_s_* if *i* = *k*, zero otherwise), and the second term represents the probability that adult *k* dies, position *k* becomes open to competition, and a descendant of adult *i* occupies this position.

##### F.2.1 Pairwise relatedness

Then, same-parent individual 2-fitness of adult 1, which we take as a focal individual, denoted by 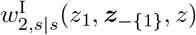 (recall eq.(26)), is written as

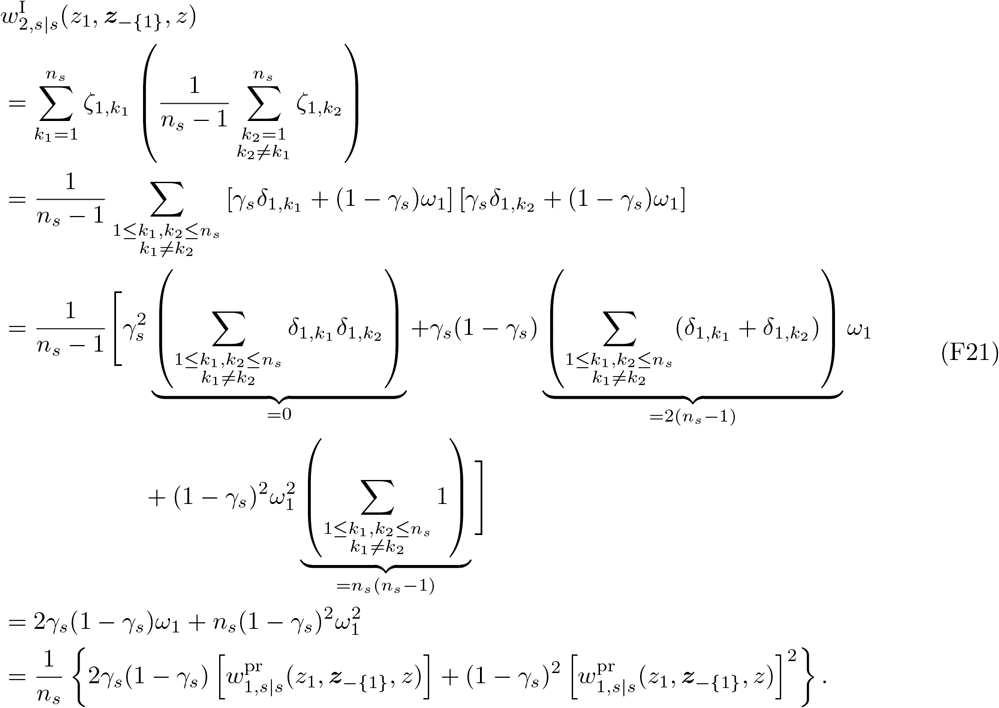

In a similar vein, different-parent individual 2-fitness is

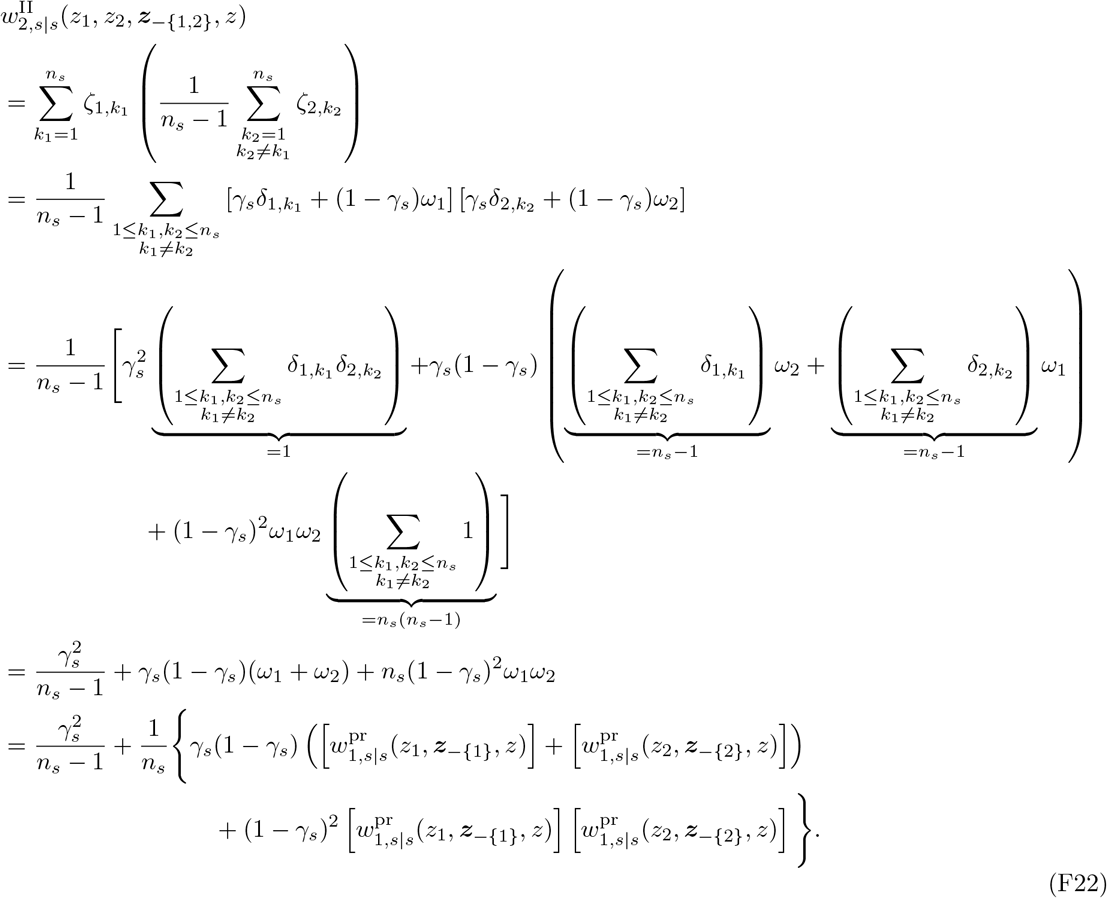

In contrast, when *s′* ≠ *s*, we have 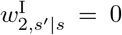 and 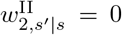 from the assumption of independent dispersal. With these expressions, we obtain

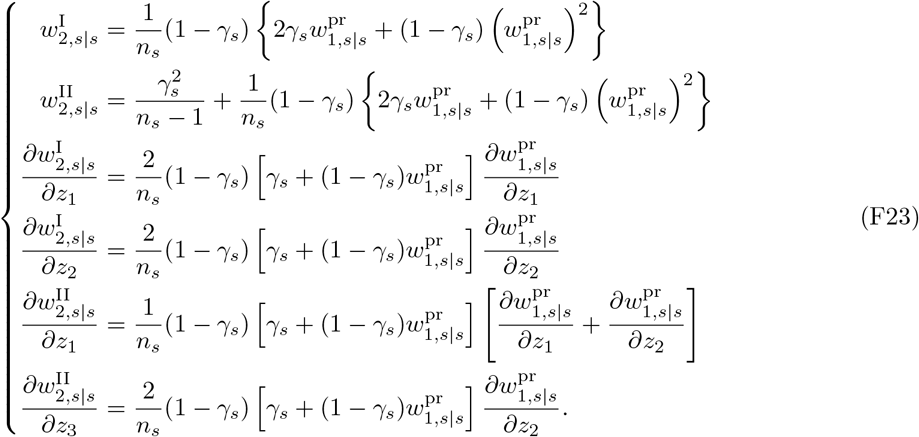

By substituting eq.(F23) in eq.(E2), we obtain a recursion on 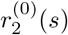 as

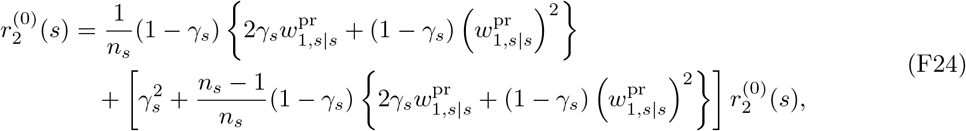

which is explicitly solved as eq.(41) in the main text.

##### F.2.2 Three-way relatedness

Next we calculate individual 3-fitness of three different types (from type-I to III) in order to calculate three-way relatedness. For type-I, we have

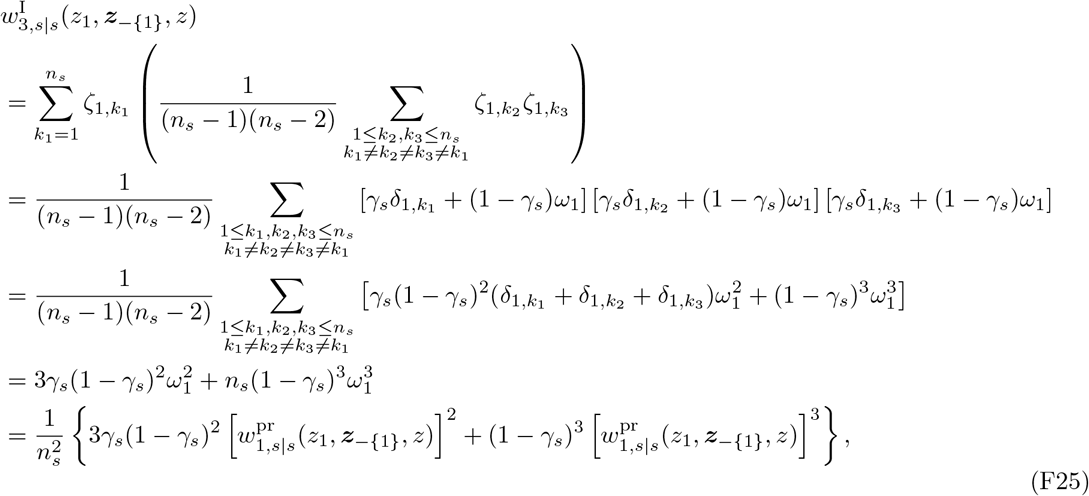

where we have used

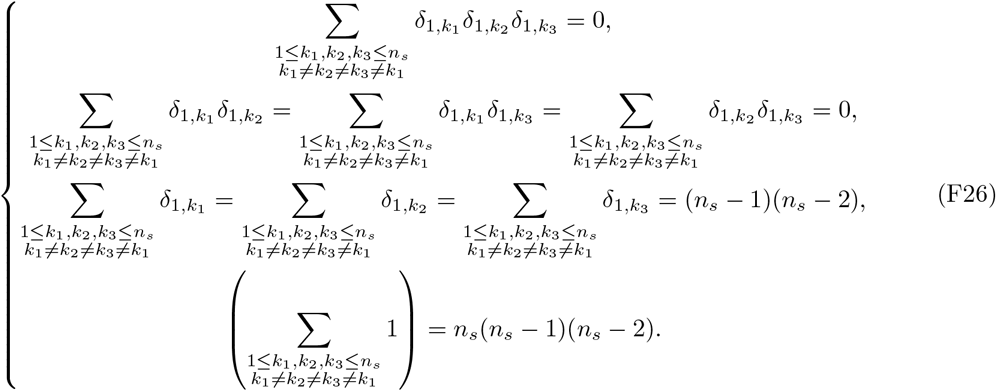

For type-II, we obtain

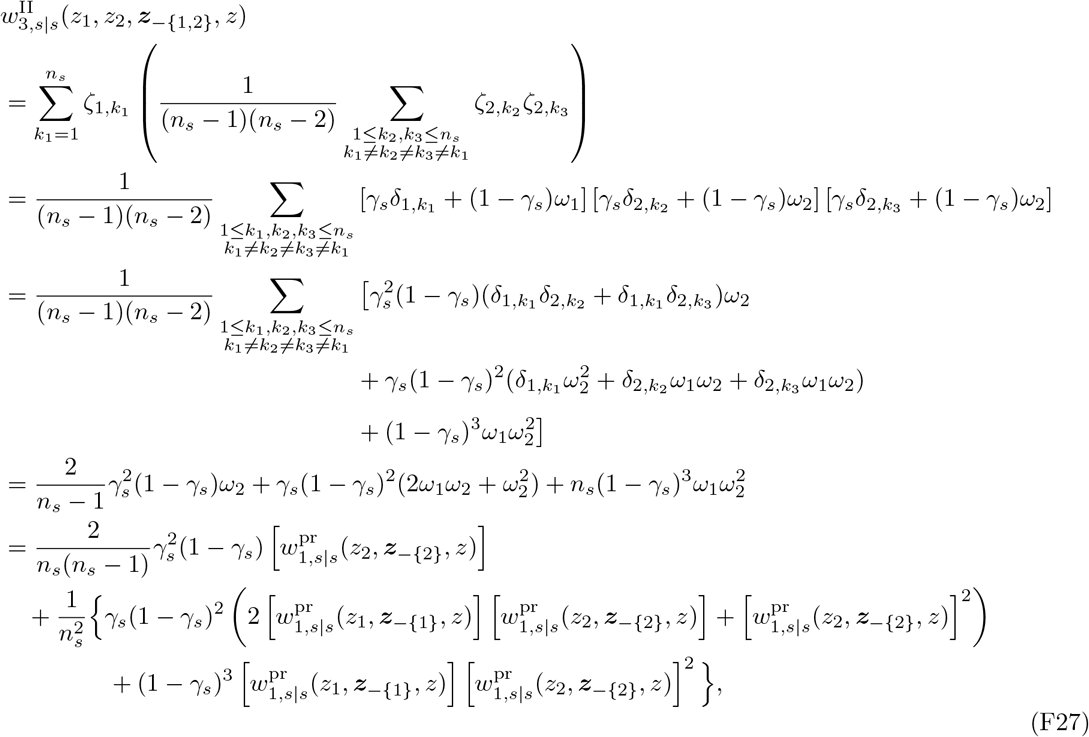

where we have used

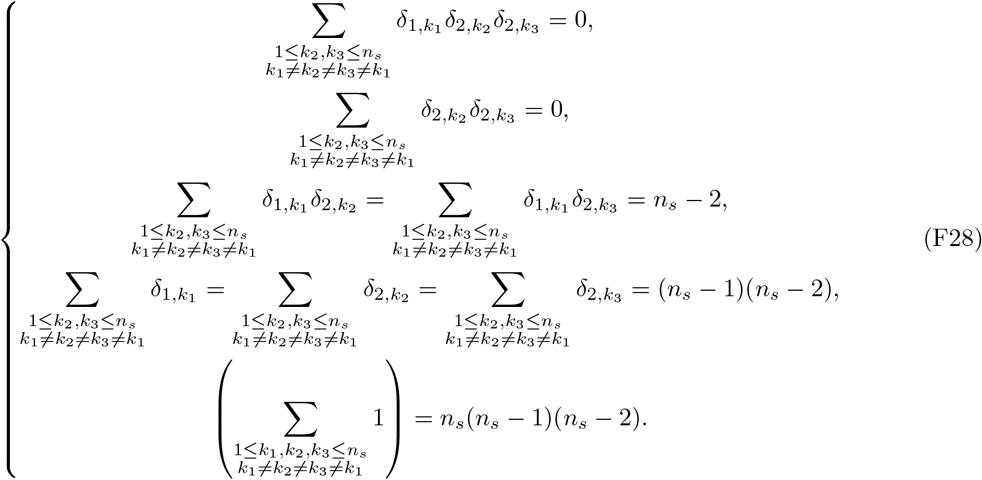

Finally, for type-III we obtain

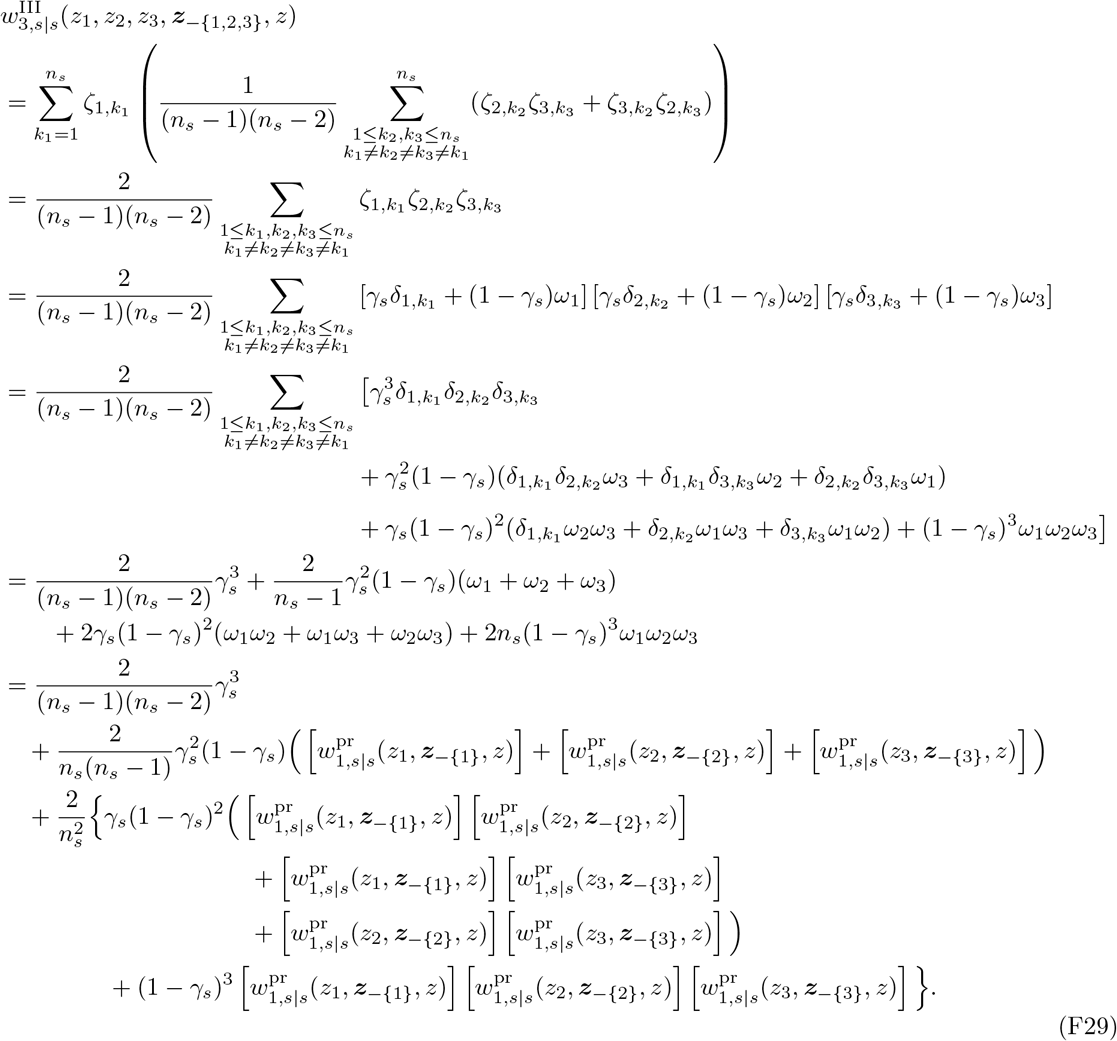

Here we have used

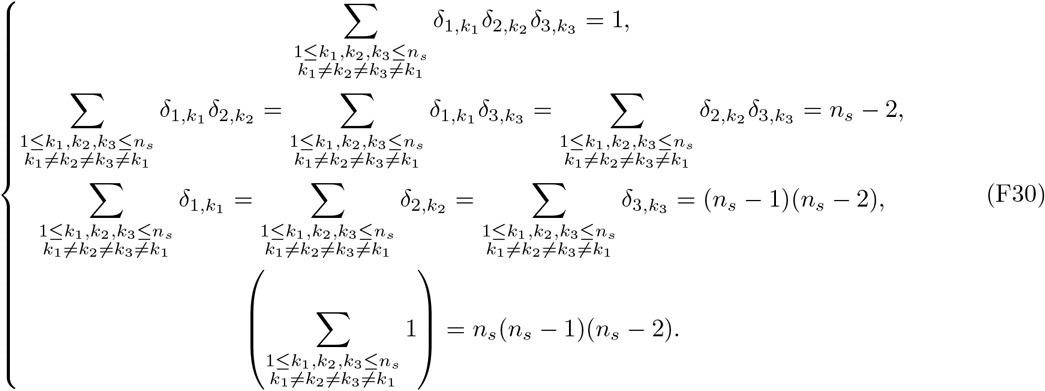

In contrast, when *s′* ≠ *s*, from the assumption of random dispersal we have 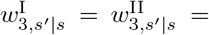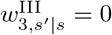. From these calculations we arrive at

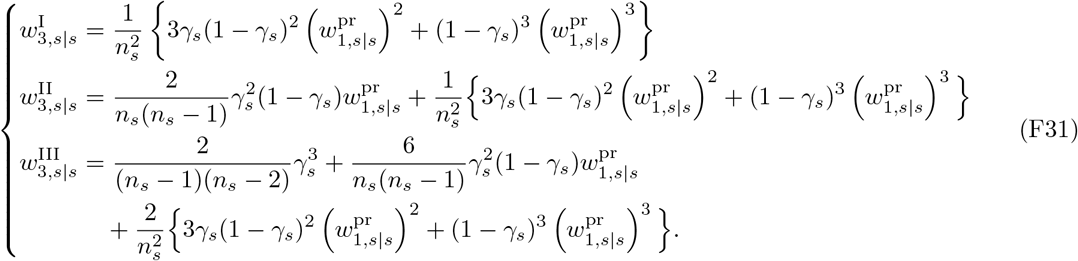

Substituting eq.(F31) in eq.(E6) gives us a recursion on 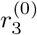 that includes 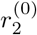. Solving it with the help of eq.(41) gives the following result:

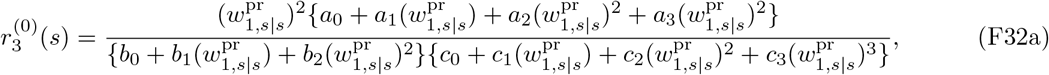

where

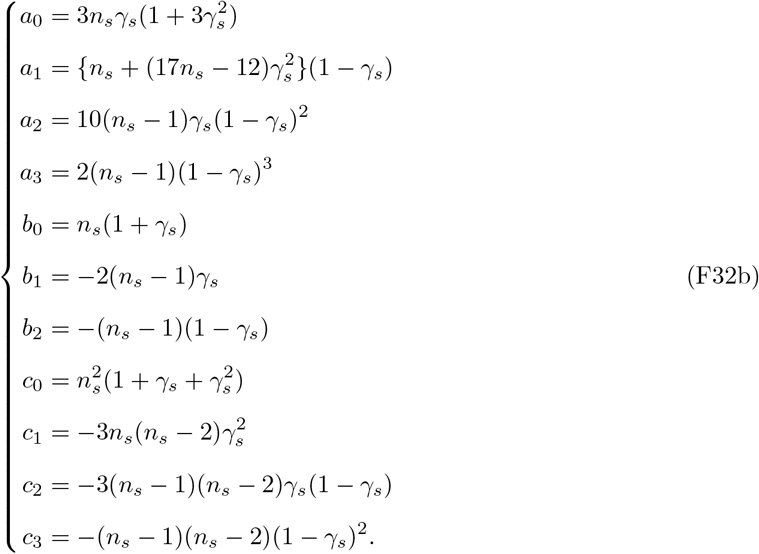

Setting *γ_s_* = 0 or *γ_s_ ∼* 1 (note that setting *γ_s_* = 1 means individuals never die and hence evolution does not occur, so we need *γ_s_* to be “close” to 1) in eq.(F32) gives eq.(42) in the main result.

##### F.2.3 First-order perturbation of pairwise relatedness

Here we assume *ρ*^(1)^ = 0 and calculate the first-order perturbation of pairwise relatedness, 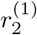.

Substituting eq.(F23) in eq.(E10) yields

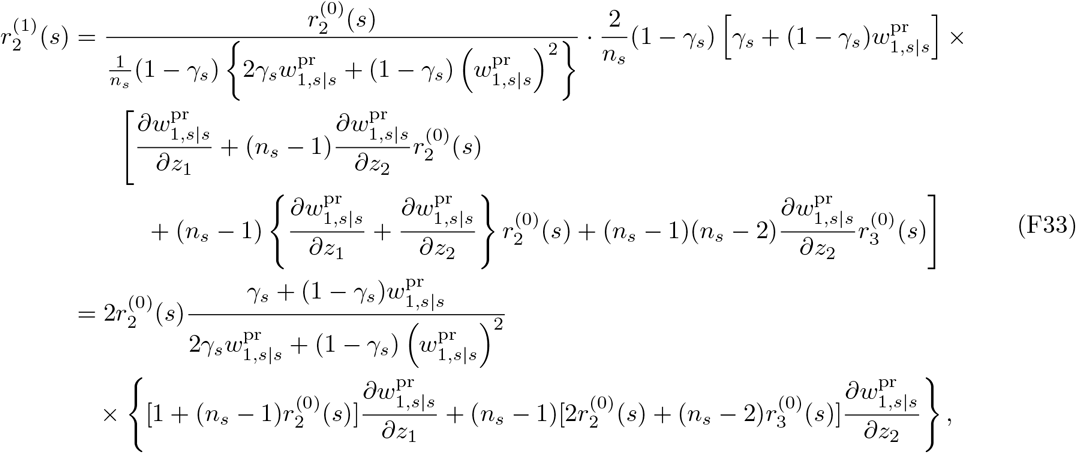

which is eq.(44) in the main text.

### G Consistency with previous results about relatedenss

We here show that we recover several previous results concerning relatedness from our model.

#### G.1 Neutral relatedness in the lottery model

Our result for the neutral pairwise relatedness given in eq.(41) agrees with *R_d_* in Pen (2000) (the solution to his eq.(A2)). To see this one has to set 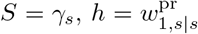 and *N* = *n_s_* in Pen (2000). Furthermore, for the special case that *γ_s_* = 0 in eq.(41) we obtain

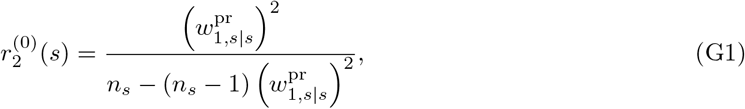

which agrees with 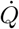 in eq.(2.9) in Rousset (2004), a standard results for the island model, by using 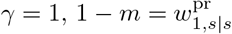 and *N* = *n_s_* there.

On the other hand, taking the limit *γ_s_ →* 1 in eq.(41) gives

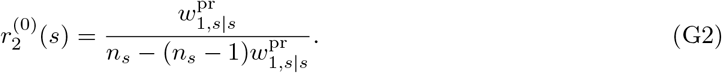

This result agrees with *r*_2_(***z, z***) = (1 *− m*(****z****))/(1 + *m*(****z****)(*N −* 1)) in Table 1 in Mullon et al. (2016) by setting 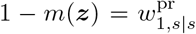 and *N* = *n_s_* in their formula. Note, that there was a typo in their original expression, which we corrected here.

Finally, as for 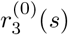, the first line of eq.(42) agrees with *R*_3_ in eq.(12b) in Ohtsuki (2010) by setting 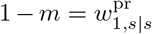 and *N* = *n_s_* there. The second line agrees with *r*_3_(***z, z***) in Table 1 in Mullon et al. (2016) by setting 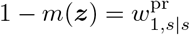 and *N* = *n_s_* there.

#### G.2 Neutral relatedness under fluctuating group size

We here prove the connection between our eq.(33) and eq.(29) of Rousset and Ronce (2004). In this latter model, individuals migrate independently from each other (no propagule migration) and states determine group size, which fluctuates stochasticaly between generations according to an ergodic Markov chain.

The probability *p*^(0)^(*s*) that a group is in state *s* in the neutral process at stationarity then satisfies

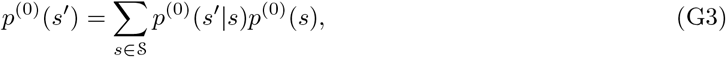

where *p*^(0)^(*s′|s*) = *p*^(0)^(*s′|s*; ****p****^(0)^) is the forward transition probability that a group in state *s* in the parental generation is in state *s′* in the offspring generation and this generally depends on whole population state ****p****^(0)^ = (*p*^(0)^(*s*_1_), *· · ·, p*^(0)^(*s_N_*)) (since groups are connected to each other by dispersal, see Metz and Gyllenberg (2001); Lehmann et al. (2006); Alizon and Taylor (2008) for concrete examples of such transition probabilities) and on the resident trait value.

In terms of these quantities, first note

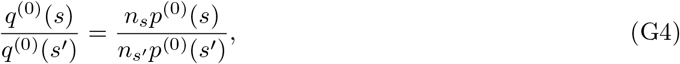

where the equality follows from eq.(E.21) of Lehmann et al. (2016). Intuitively speaking, eq.(G4) tells that the stationary fraction of individuals in group *s* under neutrality, *n_s_p*^(0)^(*s*), is proportional to the stationary distribution of mutants under neutrality, *q*^(0)^(*s*). Second, because the model of Rousset and Ronce (2004) did not allow any propagule dispersal, the only way that a focal individual in a group in state *s* earns individual 2-fitness is that the state of the group changes from state *s* in the parental generation to state *s′* in the offspring generation and the focal individual produces offspring in the focal group. Thus, we obtain

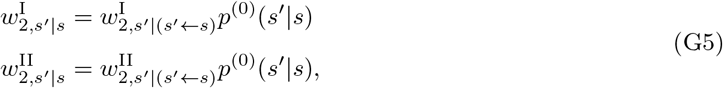

where 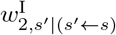 and 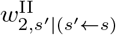 are conditional 2-fitness components of the focal individual (they follow the same definition as 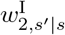 and 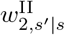 but are conditional on the parental generation being in state *s* and the offspring generation being in state *s′*) evaluated under neutrality.

Substituting eqs.(G4)–(G5) into eq.(33) and using the backward transition probability

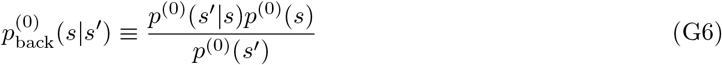

that a group in state *s′* in the offspring generation was in state *s* in the previous generation, we obtain that

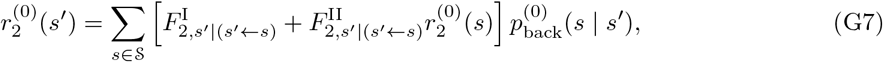

where

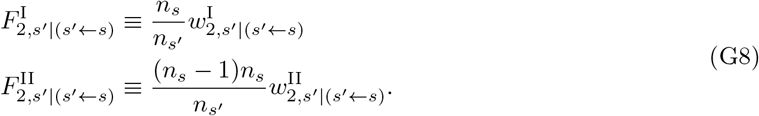

We now claim that 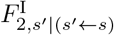 is the probability that two randomly sampled offspring in state *s′* both descend from the same parent in state *s* (*i.e.*, the coalescence probability) under neutrality and that 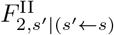 is the probability that two randomly sampled offspring in state *s′* both descend from two distinct parents in state *s* under neutrality. If this interpretation holds, then our eq.(G7) reduces to eq.(29) of Rousset and Ronce (2004).

We now proceed to prove this claim. Given that the state of the group is *s* in the parental generation and *s′* in the offspring generation, from the definition of (conditional) same-parent 2-fitness, we can write

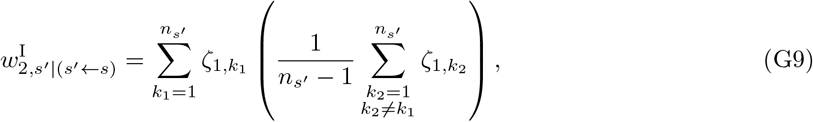

where, we use the *ζ*-notation once used in section F.2; this time *ζ_i,k_* represents the probability that an individual in position *k ∈* {1, *· · ·, n_s′_*} in the next generation in the focal group descends from adult *i ∈* {1, *· · ·, n_s_*} in that group in the previous generation under neutrality conditioned on that the group state has changed (forwardly in time) from *s* to *s′*.

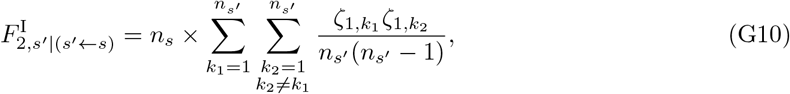

which, owing to neutrality (and thus exchangeability of individuals), can be rewritten as

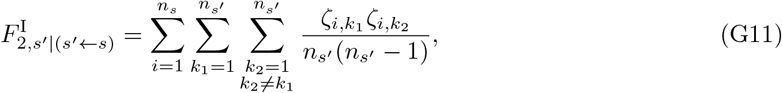

and therefore can be read as the ratio of the total number of ways of sampling two offspring from the same parent in a group of size *n_s_* to the total number of ways of sampling two offspring in a group of size *n_s′_* (*i.e.*, the coalescence probability). Likewise, we have

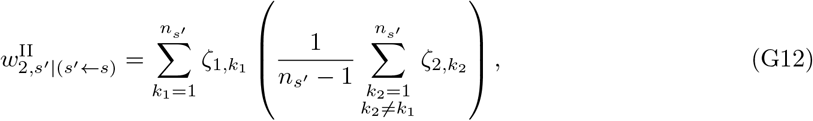

whereby

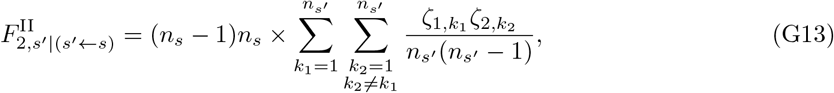

which, owing to neutrality and exchangeability of individuals, can be rewritten as

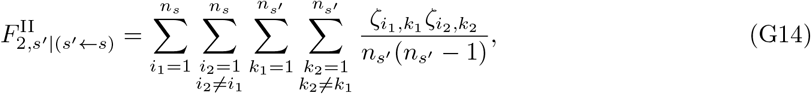

and thus can be read as the ratio of the total number of ways of sampling two offspring from the distinct parents in a group of size *n_s_* to the total number of ways of sampling two offspring in a group of size *n_s′_* (*i.e.*, the proability that offspring in state *s′* descend from two distinct parents in state *s*). This ends the proof of our aforementioned claim.

#### G.3 Perturbation of relatedness

As for 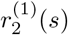, eq.(44) evaluated at *γ_s_* = 0 reads

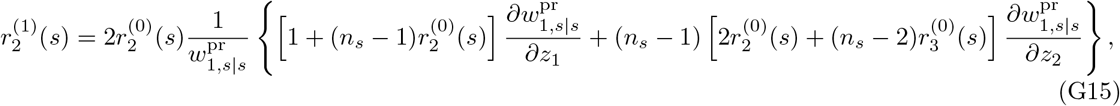

which corresponds to the expression that appear in Δ*r* in Wakano and Lehmann (2014) (see the bottom of their Appendix B, after their eq.(B.46)), which essentially shows that the perturbation of pair-relatedness is

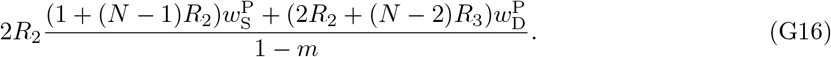

The correspondence between eq.(G15) and eq.(G16) is as follows. Set 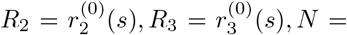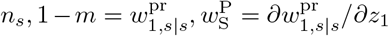, and 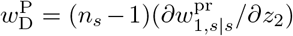 in eq.(G16). Note that the original expression at the bottom of Appendix B in Wakano and Lehmann (2014) contains the factor 4, not 2. This two-fold difference comes from the fact that Wakano and Lehmann (2014) considered variance dynamics of a trait distribution, whereas we here consider directly the perturbation of *r*_2_(*s*).

### H Perturbations for the hard selection lottery model

Here, we derive the expressions for *ρ*^(1)^ and *ρ*^(2)^ for the lottery model under hard selection. The resulting expression are complicated and do not appear in the main text, but they can be useful for numerical calculations as they apply to any fecundity function given the model’s other assumptions.

We recall that for hard selection, from eqs.(38b) and (38c) with eqs.(45a) and (46a), we have

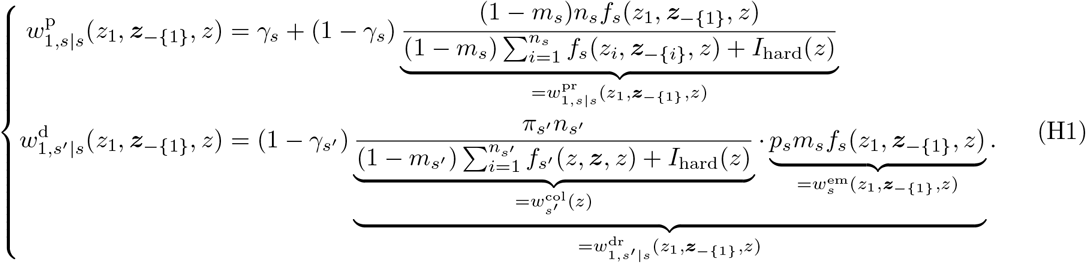

#### H.1 First-order perturbation of invasion fitness

Our goal here is to calculate the first-order perturbation of invasion fitness, given by eq.(32). For that purpose, we will calculate the components that appear in eq.(32), as below.

Firstly, we will calculate *v*^(0)^(*s′*)*q*^(0)^(*s*). With the definition of backward migration probability, eq.(49), we see that

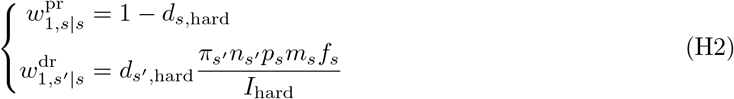

hold. Substituting them into eq.(40) gives

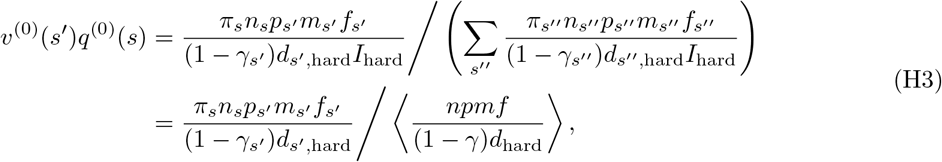

where we set

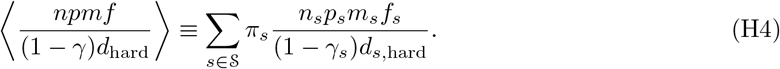

Secondly, we will calculate two different derivatives of individual 1-fitness that appear in eq.(32). A direct calculation of eq.(H1) shows that

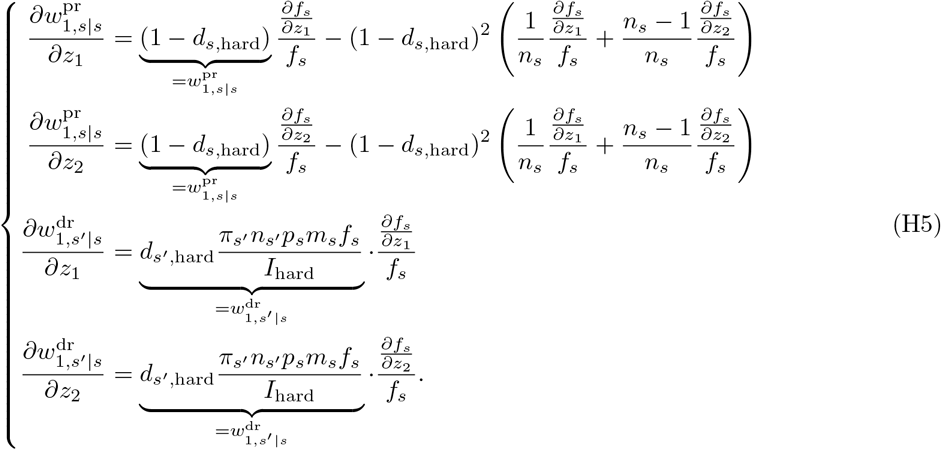

Remember that from eq.(38)

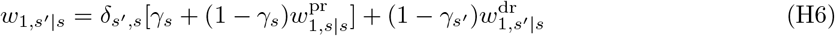

(where *δ_s′,s_* is Kronecker’s delta; it is 1 if *s′* = *s* and is 0 if *s′* ≠ *s*) holds. Taking the first-order derivative of eq.(H6), substituting eq.(H5) therein, and using eq.(H6) again for rewriting give us

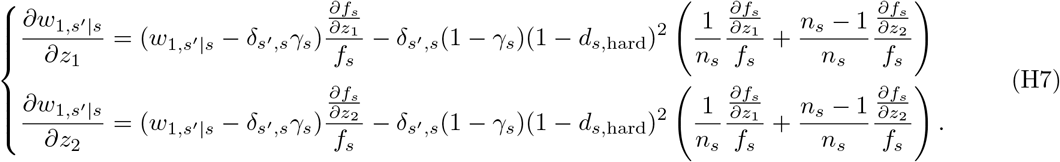

Now we are ready to calculate the first-order perturbation of invasion fitness, eq.(32). We substitute eq.(H7) into eq.(32). We show this calculation piece by piece. Firstly,

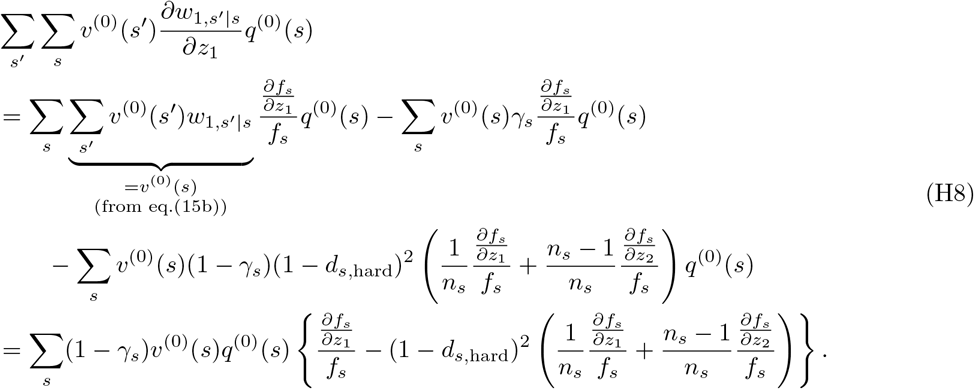

Secondly, a quite similar calculation to above leads to

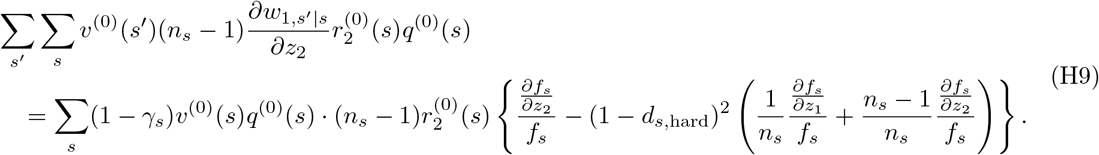

By combining eqs.(H8) and (H9) in eq.(32), we obtain the first-order perturbation as

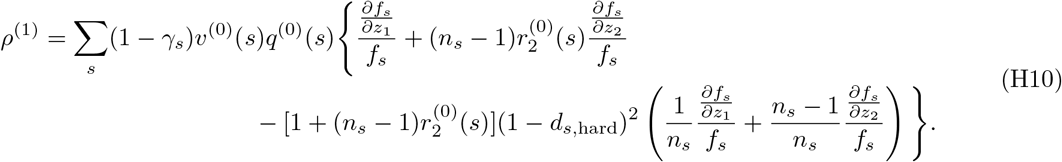

Rewriting it by using eqs.(50) and (H3) gives eq.(48) in the main text. Here, 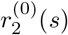 is calculated by substituting eq.(H2) into eq.(41). The result is shown in eq.(51) in the main text.

#### H.2 Second-order perturbation of invasion fitness

We now assume that *ρ*^(1)^ = 0. Our goal here is to calculate the second-order perturbation of invasion fitness, given by eq.(34). We have already calculated first-order derivatives that appear there, as given in eq.(H7). We will then calculate various second-order derivatives that appear in eq.(34). A direct calculation of eq.(H1) shows that relevant terms are

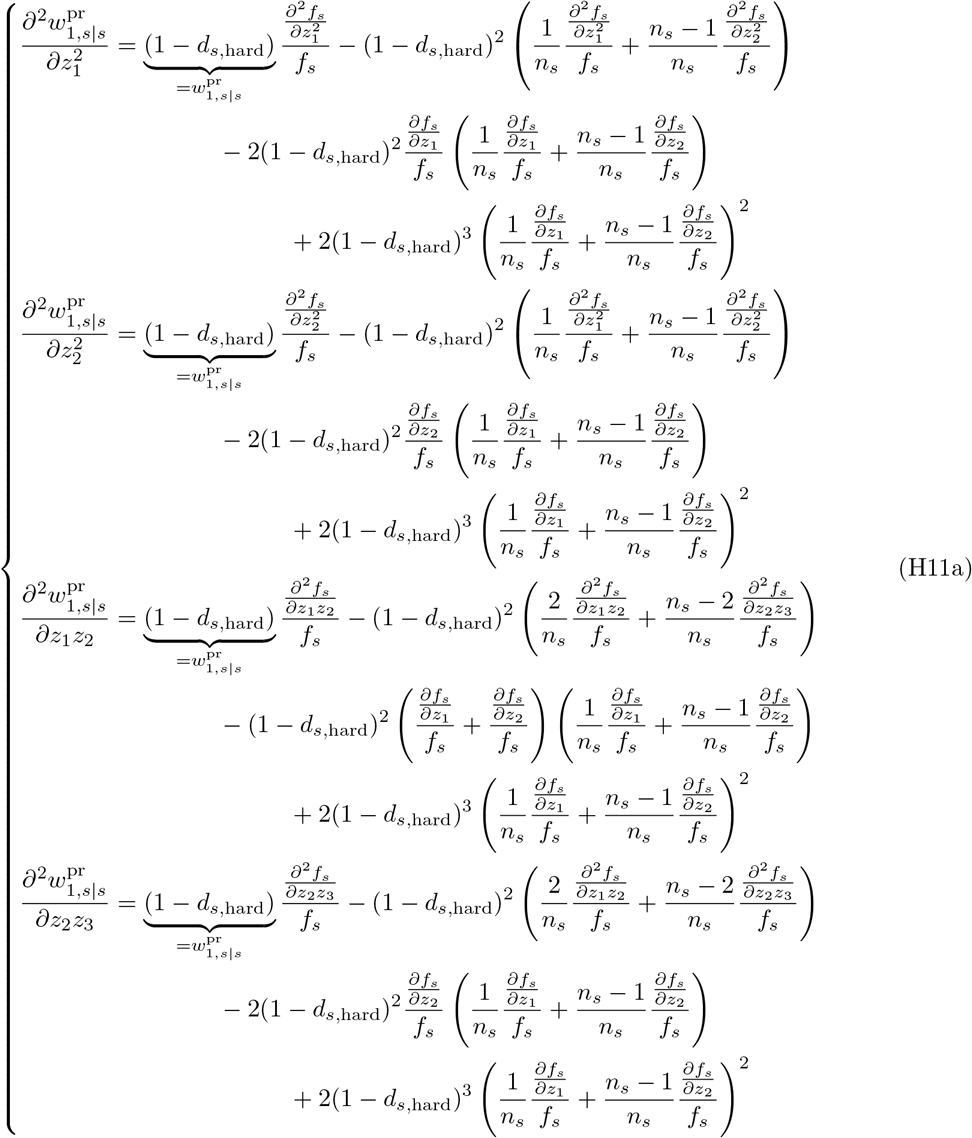

and

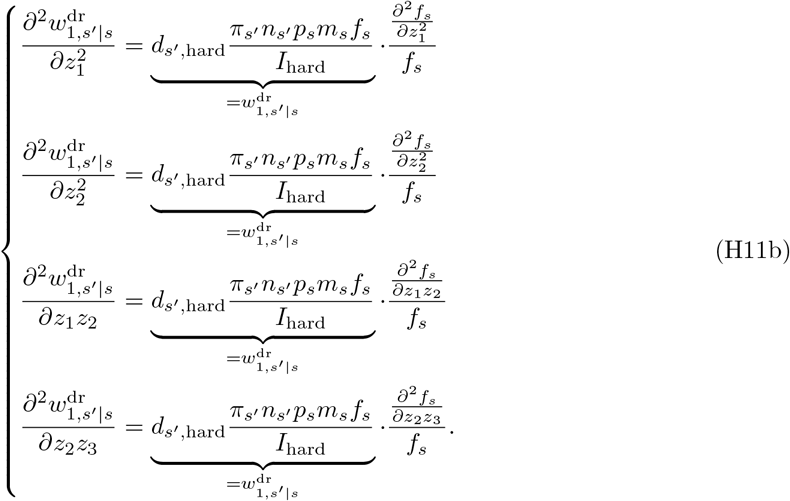

Taking the second-order derivative of eq.(H6), substituting eq.(H11) therein, and using eq.(H6) again for rewriting yields

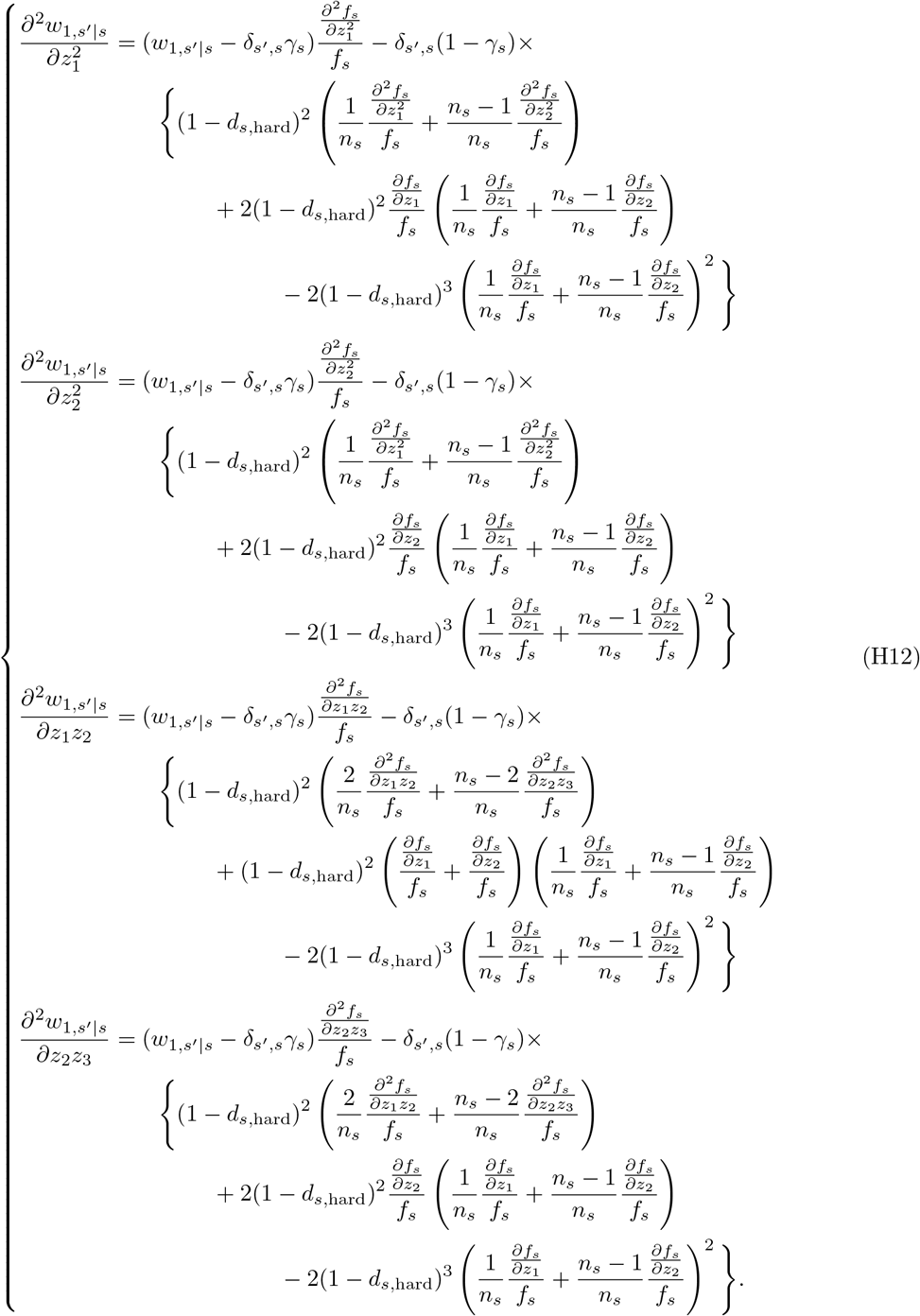

Now we are ready to calculate eq.(34). We start from *ρ*^(2w)^, which is given by eq.(34a). We substitute eq.(H12) in eq.(34a). The four terms in eq.(34a) are then calculated as follows. For example,

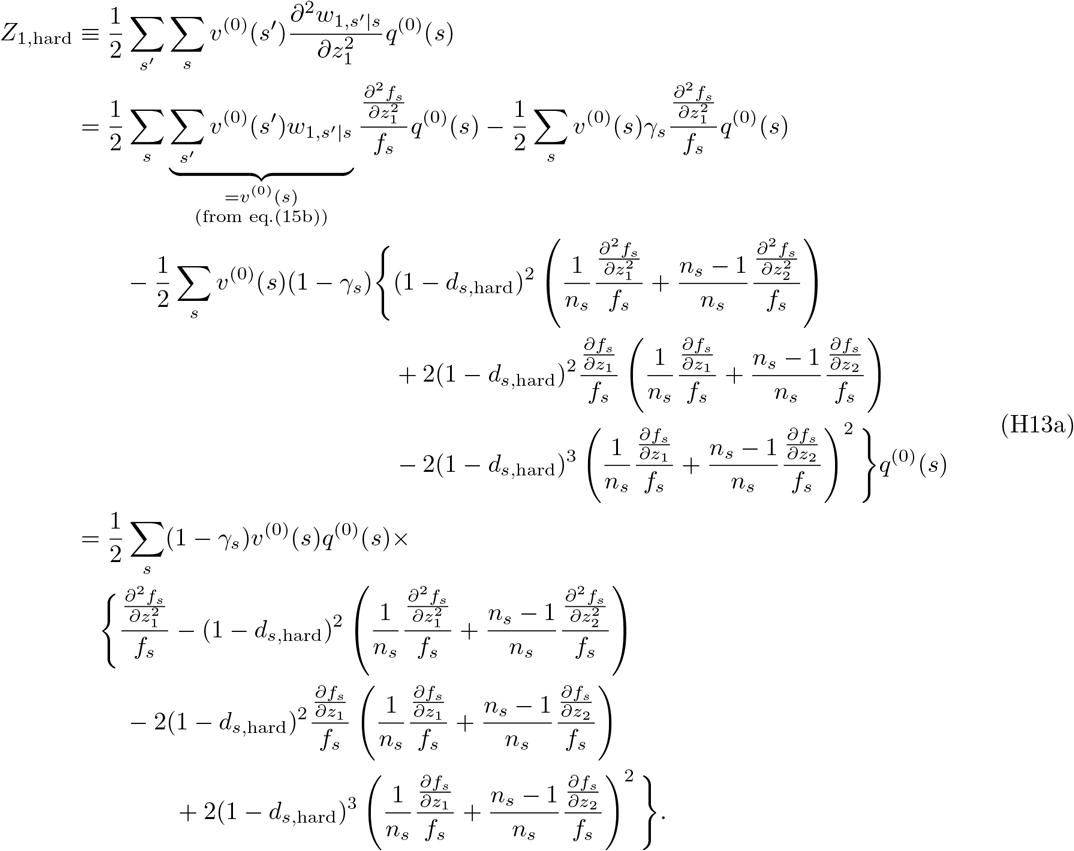

The other terms that appear in eq.(34a) are calculated in essentially the same way as

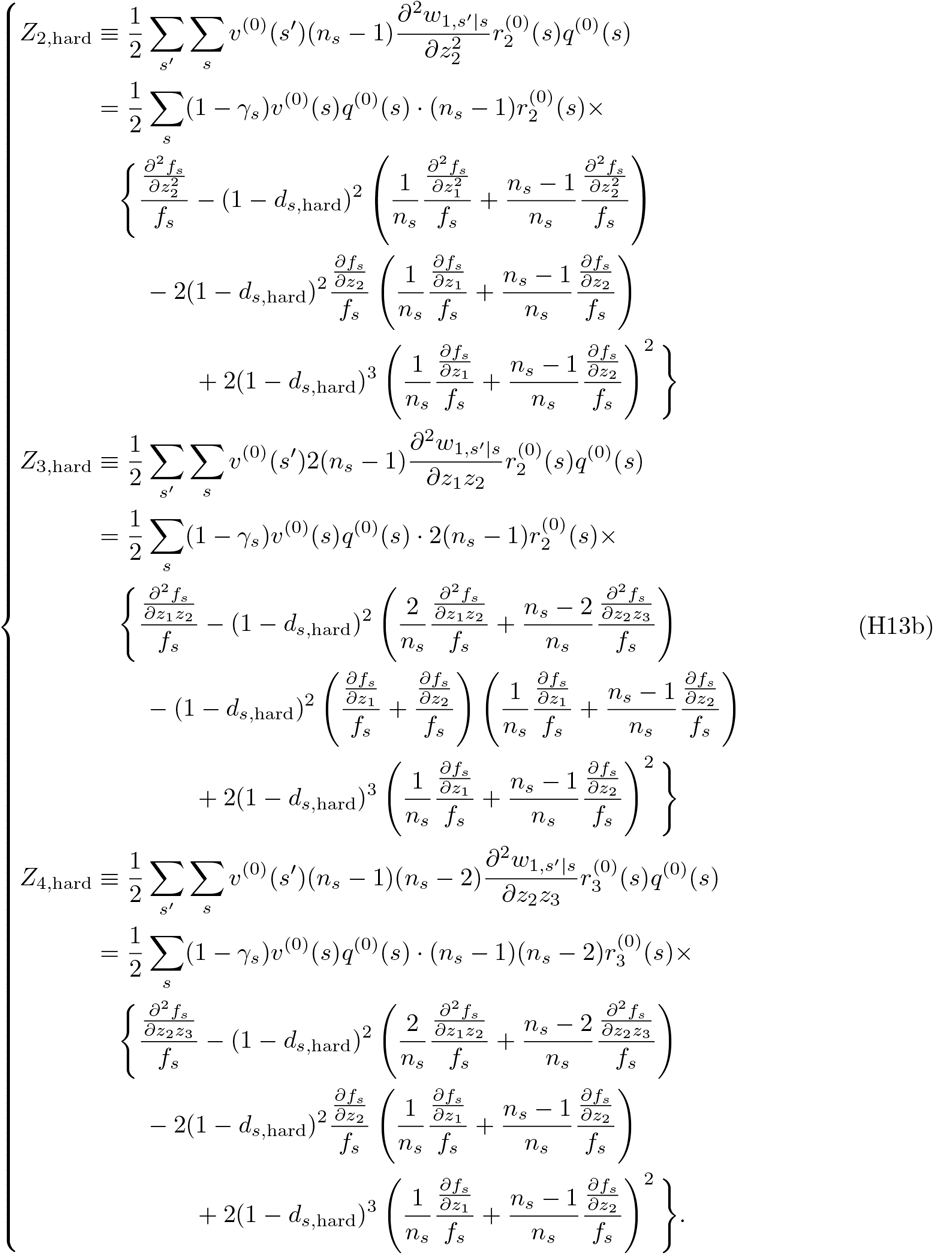

Collecting the *Z*-terms in eqs.(H13a) and (H13b) gives us

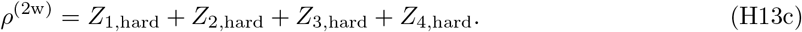

Here, 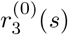 is calculated by substituting eq.(H2) in eq.(F32).

Next, we calculate *ρ*^(2q)^, which is given by eq.(34b). We repeat the same calculations as eqs.(H8) and (H9), but this time *q*^(0)^ is replaced with *q*^(1)^ there. From eq.(H10), the result is

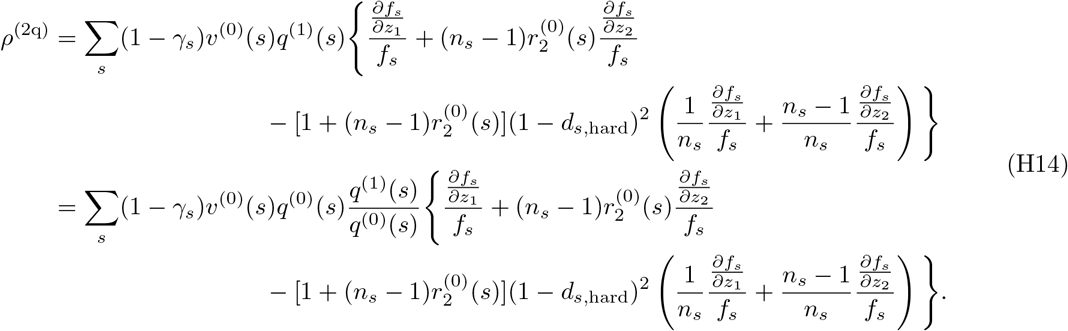

By substituting eqs.(H2) and (H5) into eq.(43) we obtain

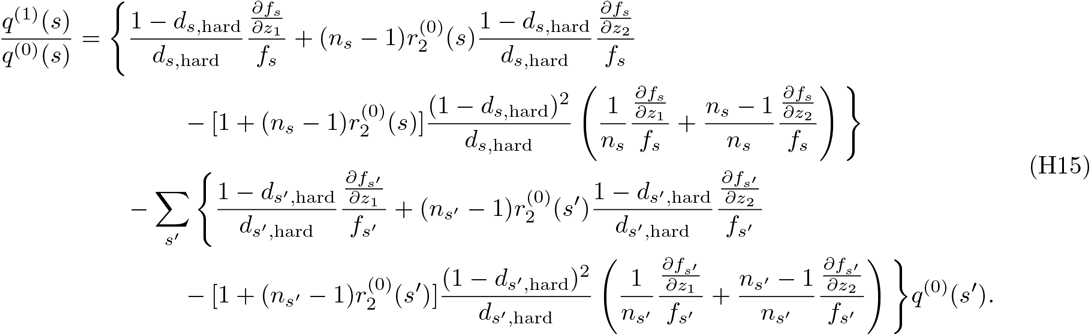

Now we substitute eq.(H15) into eq.(H14). Before doing so, we observe that the second term in eq.(H15), which is made of the sum over *s′*, is just a constant, and that putting a constant in the place of *q*^(1)^(*s*)*/q*^(0)^(*s*) in eq.(H14) gives *ρ*^(1)^ times this constant (see eq.(H10)), which is zero by the assumption in this subsection. Thus we can substitute only the first term of eq.(H15) into eq.(H14) to obtain *ρ*^(2q)^, which leads to

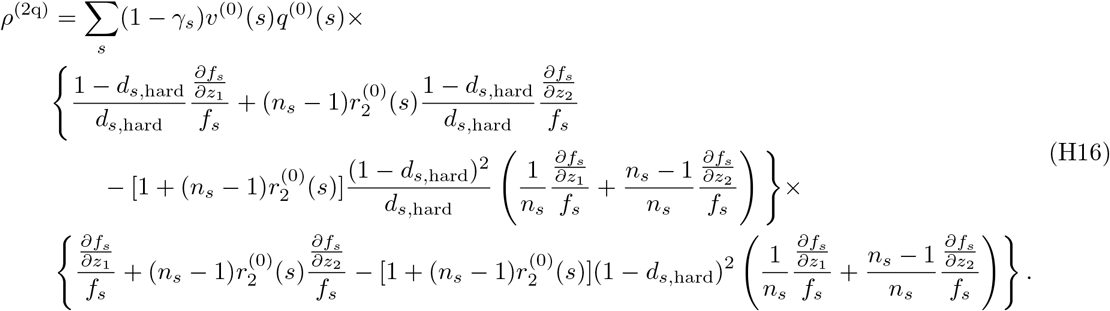

Thirdly, we calculate *ρ*^(2r)^, which is given by eq.(34c). For this, we repeat the calculation in eq.(H9), but this time 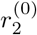 is replaced with 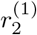 there. The result is

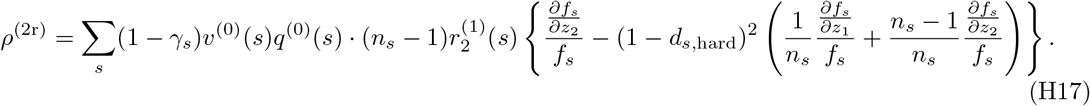

Here, we obtain 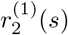 by substituting eqs.(H2) and (H5) in eq.(44), as

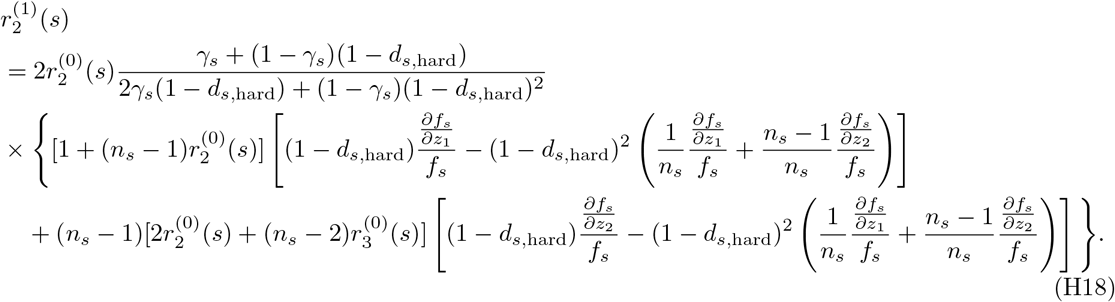

#### H.3 Consistency with previous results about perturbations

We here show that we recover several previous results from our model.

##### H.3.1 A model with overlapping generations by Lehmann and Rousset (2010)

Suppose there is a single habitat (*N* = 1) and no mortality in migration (*p_s_* = 1). Then eq.(48) becomes

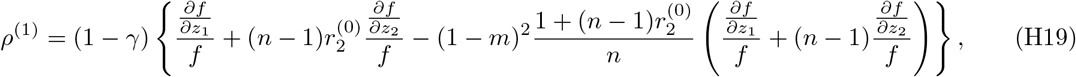

which precisely recovers the inclusive fitness effect *S*_IF_ shown in eq.(A.21) in Lehmann and Rousset (2010) with the following correspondence; 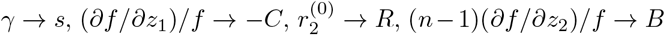, and 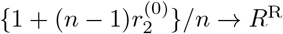.

##### H.3.2 A spatially heterogeneous model by Rodrigues and Gardner (2012)

Rodrigues and Gardner (2012) studied effects of spatial and temporal heterogeneity of patch quality on the evolution of helping/harming. Their results on spatial heterogeneity (Results 1 and 2 therein) readily follow from our eq.(48). To recover them, set *N* = 2 (high/low quality patches) and 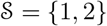. Also, set *γ*_1_ = *γ*_2_ = 0, *n*_1_ = *n*_2_ = *n*(*≥* 2), *p*_1_ = *p*_2_ = 1, and *m*_1_ = *m*_2_ = *m*(*>* 0), which leads to

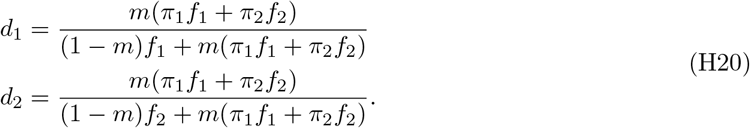

Therefore, eq.(48) becomes

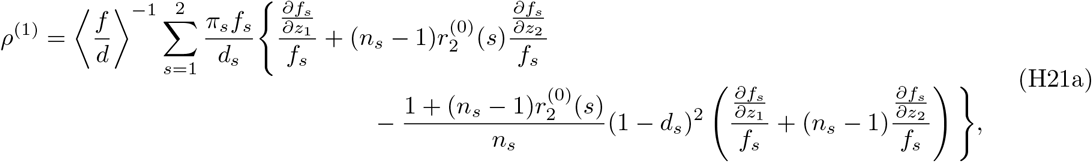

where

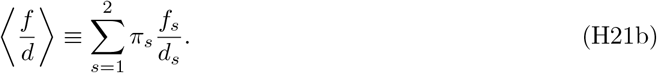

For the Wright-Fisher update (*γ*_1_ = *γ*_2_ = 0), via a direct calculation of eq.(51) we can confirm

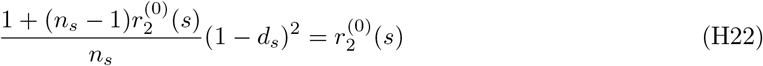

holds for *n ≥* 2. Simplifying eq.(H21) by using eq.(H22) leads to

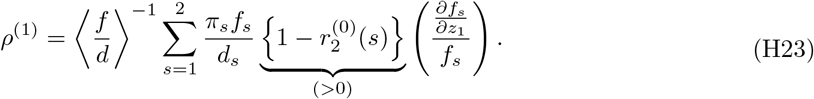

Notice that effect on others’ fecundity, *∂f_s_/∂z*_2_, is completely absent above. Thus we conclude that, as long as there is cost in helping/harming in both group states (*i.e. ∂f_s_/∂z*_1_ *<* 0 for *s* = 1, 2), neither obligate nor facultative helping/harming evolves, which essentially echoes Results 1 and 2 in Rodrigues and Gardner (2012).

##### H.3.3 A cancellation result by Mullon et al. (2016)

Consider *N* = 1 (only one state) and the limit of *γ_s_ →* 1 (Moran process). Also, suppose *ρ*^(1)^ = 0 (first-order perturbation of invasion fitness is null). This means that the expression inside the curly brackets of eq.(H10) is null. By applying eq.(51), this condition can be rewritten as

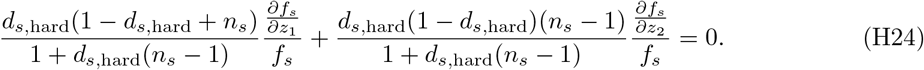

Meanwhile, consider 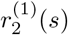. It is proportional to the expression inside the curly brackets of eq.(H18), that is

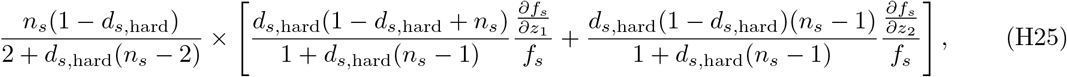

which is zero. Hence 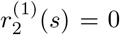. This suggests that the first-order perturbation of relatedness in the Moran process is null, and therefore that the component of second-order perturbation of invasion fitness due to perturbation of relatedness is

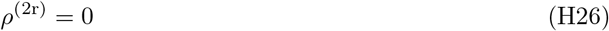

in the Moran process.

Mullon et al. (2016) found essentially the same result (see their eq.(16)) for their Moran model, although their model was slightly different from ours here; Mullon et al. (2016) assumed that in each (non-extinct) patch exactly one adult individual always dies at each update step, whereas our model assumes that death occurs randomly to each individual and that it occurs rarely (*γ_s_ →* 1).

##### H.3.4 Second-order results by Parvinen et al. (2018)

Parvinen et al. (2018) calculated the metapopulation fitness of mutants, *R*_m_, in a subdivided population by assuming non-overlapping generations, *γ_s_* = 0 (Wright-Fisher process), uniform migration rate (*m_s_* = *m*), and uniform death rate during dispersal (*p_s_* = *p*). It is known that *R*_m_ *−* 1 has the same sign as *ρ −* 1 (Lehmann et al. (2016)), and therefore that metapopulation fitness can be used as a proxy to determine evolutionary success of mutants.

Parvinen et al. (2018) calculated a Taylor expansion of *R*_m_ with respect to mutational deviation, *δ*:

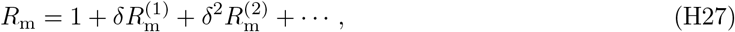

where 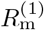 corresponds to their *D*_1_ (*s*_res_) (see their eq.(3.5)), and 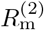 corresponds to their *D*_2_ (*s*_res_)/2 (see their eq.(3.10)).

By using our eq.(H10) we can confirm (calculations are not shown here because they are too long) that

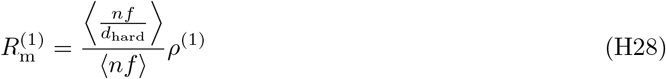

holds, where

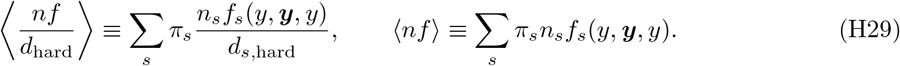

This result was firstly shown by Parvinen et al. (2018) (see their eq.(B.26)). Similarly, when *ρ*^(1)^ = 0, by using eqs.(H13), (H16) and (H17) we can confirm (calculations are not shown here because they are too lengthy) that

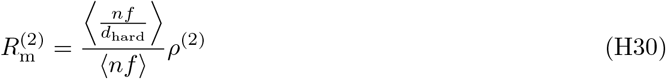

holds, as expected. These work as indirect confirmations that our results, eq.(H10) (for first-order) and eqs.(H13), (H16) and (H17) (for second-order), are correct.

### I Perturbations for the soft selection lottery model

In this section, we derive the expressions for *ρ*^(1)^ and *ρ*^(2)^ for the lottery model under soft selection. Recall that for this model; namely, eqs.(38b) and (38c) with eqs.(45b) and (46b), we have

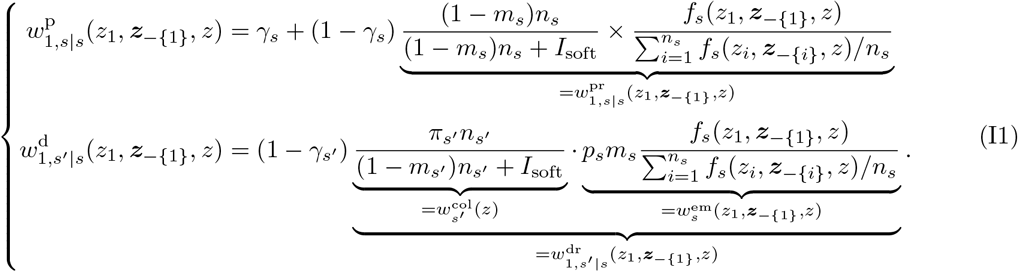

#### I.1 First-order perturbation of invasion fitness

The goal here is to calculate the first-order perturbation of invasion fitness, given by eq.(32). For this purpose we will calculate its components one by one. First, we derive *v*^(0)^(*s′*)*q*^(0)^(*s*). With the definition of the backward migration probability (eq.(63)) we obtain

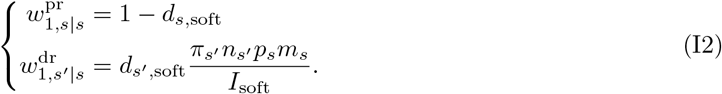

Substituting them into eq.(40) yields

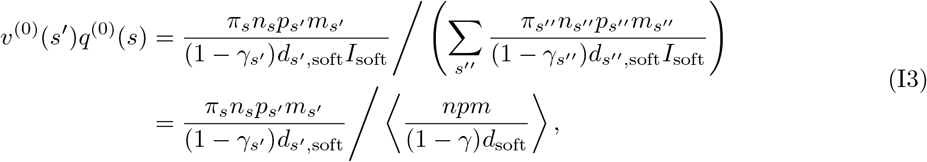

where

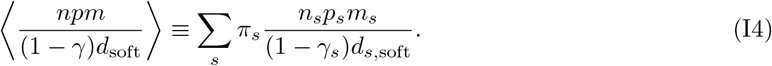

Second, we calculate first-order derivatives of 1-fitness that appear in eq.(32). A direct calculation of eq.(I1) shows that relevant first-order derivatives are

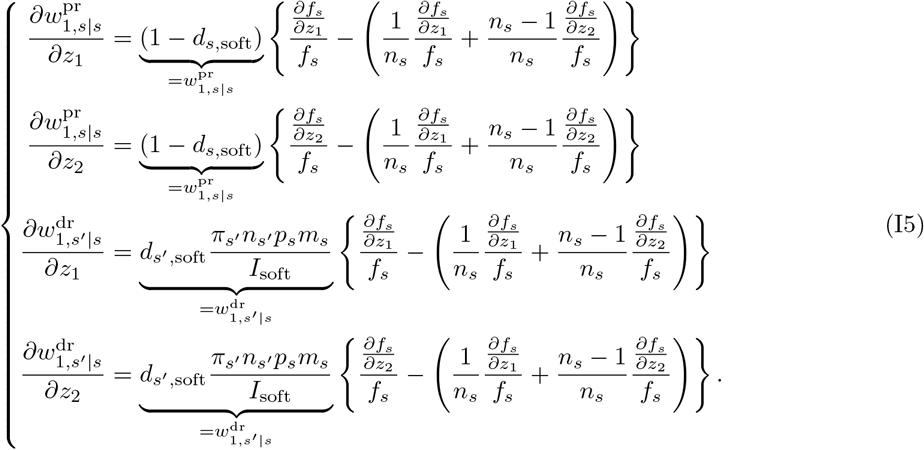

Taking the first-order derivative of eq.(H6), substituting eq.(I5) therein, and using eq.(H6) again for rewriting give us

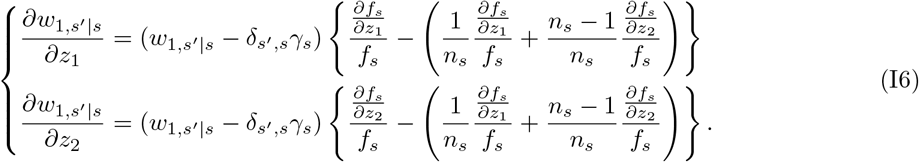

Finally, with eqs.(I3) and (I6) we can calculate *ρ*^(1)^. We observe that each equation in eq.(I6) has the factor (*w*_1,*s*′|s_ − δ_s′,s_γ_s_) in common. Thus the following relation is useful in the following calculation; for any function *F* (*s*) we have

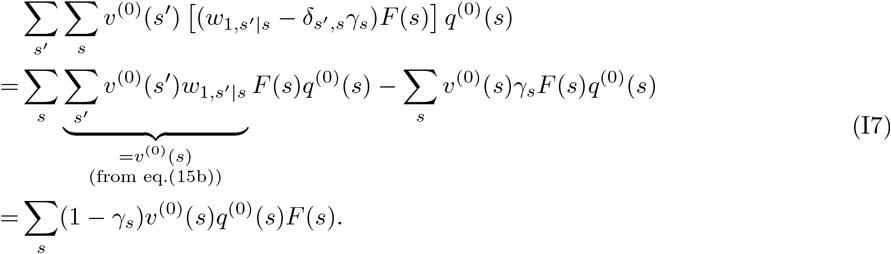

The first term in eq.(32) is then

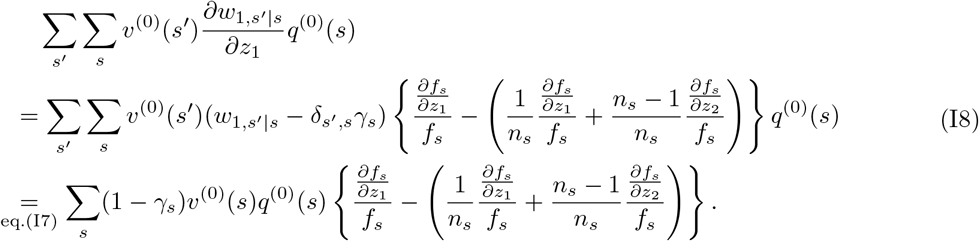

The second term in eq.(32) is similarly calculated as

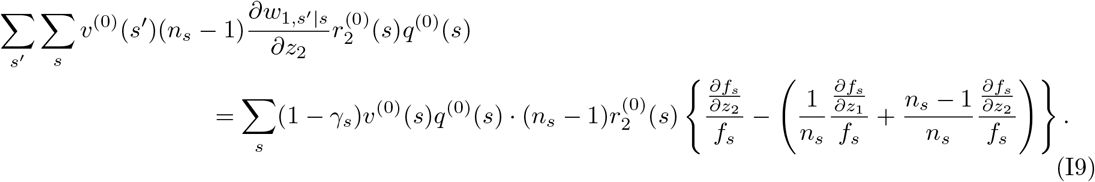

By combining eqs.(I8) and (I9) we obtain

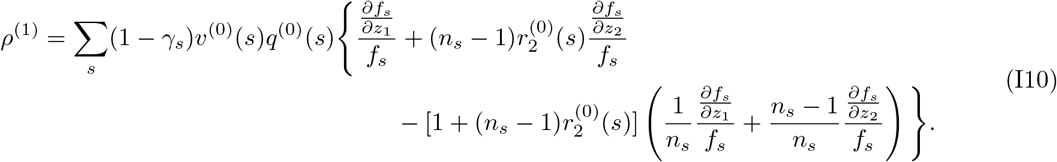

Rewriting eq.(I10) with eqs.(50) and (I3) gives eq.(62) in the main text. Here, 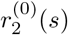 is obtained by substituting eq.(I2) into eq.(41). We find that 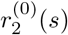 takes exactly the same form as eq.(51) except that all *d_s,_*_hard_ there should be replaced by *d_s,_*_soft_.

#### I.2 Second-order perturbation of invasion fitness

Below we assume *ρ*^(1)^ = 0. The goal here is to calculate the second-order perturbation of invasion fitness, given by eq.(34). As in the hard selection case, we have already calculated first-order derivatives that appear there, as given in eq.(I6). We will then calculate various second-order derivatives that appear in eq.(34). A direct calculation of eq.(I1) shows that relevant terms are

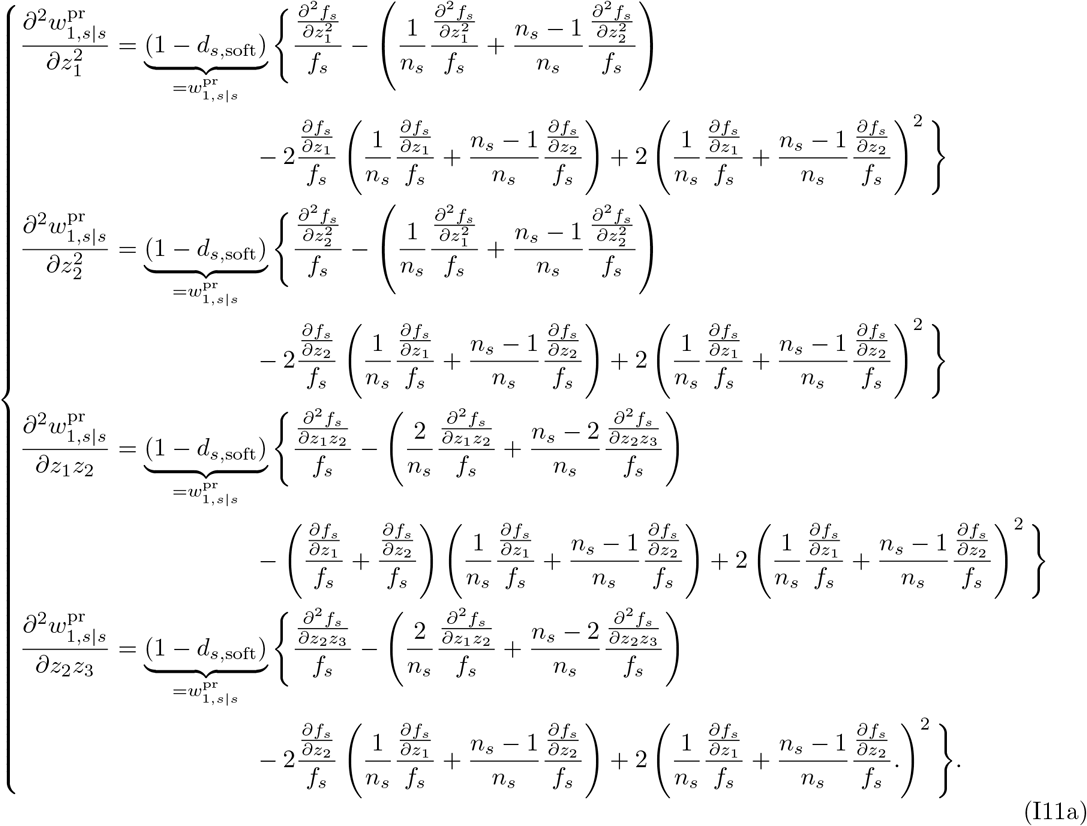

and

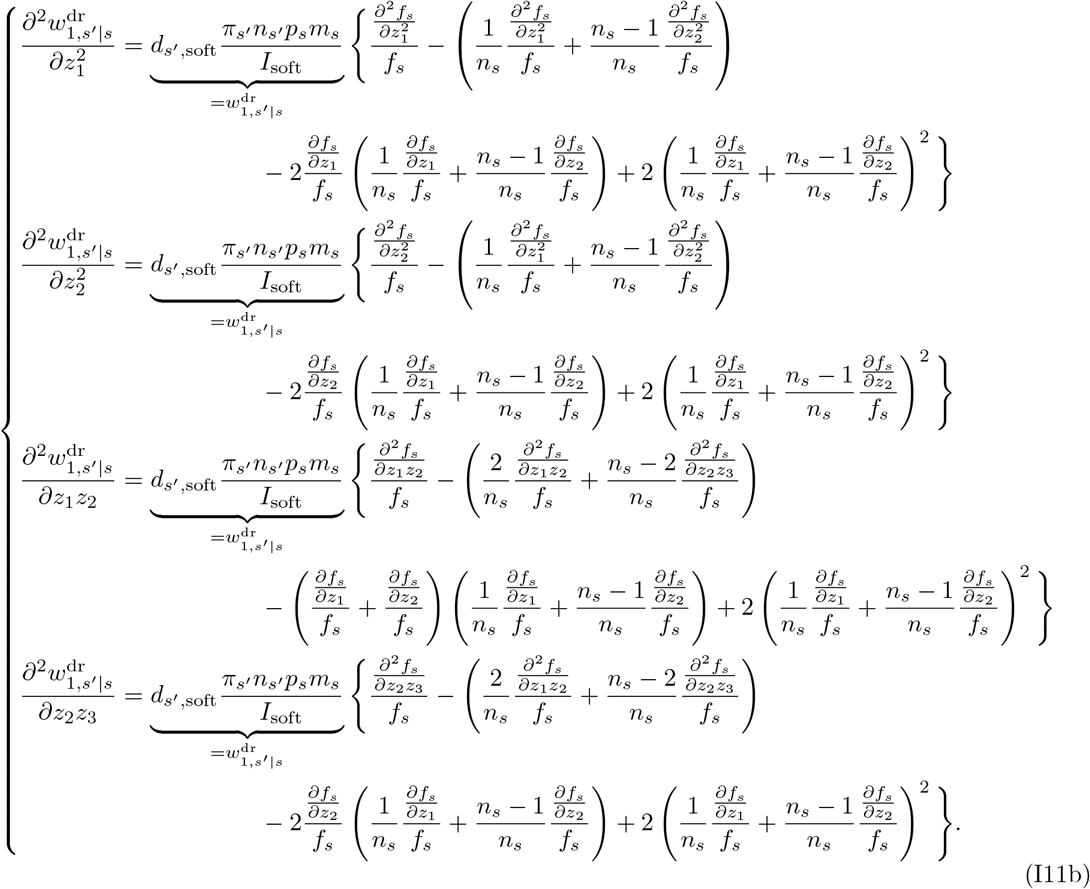

Taking the second-order derivative of eq.(H6), substituting eq.(I11) therein, and using eq.(H6) again for rewriting yields

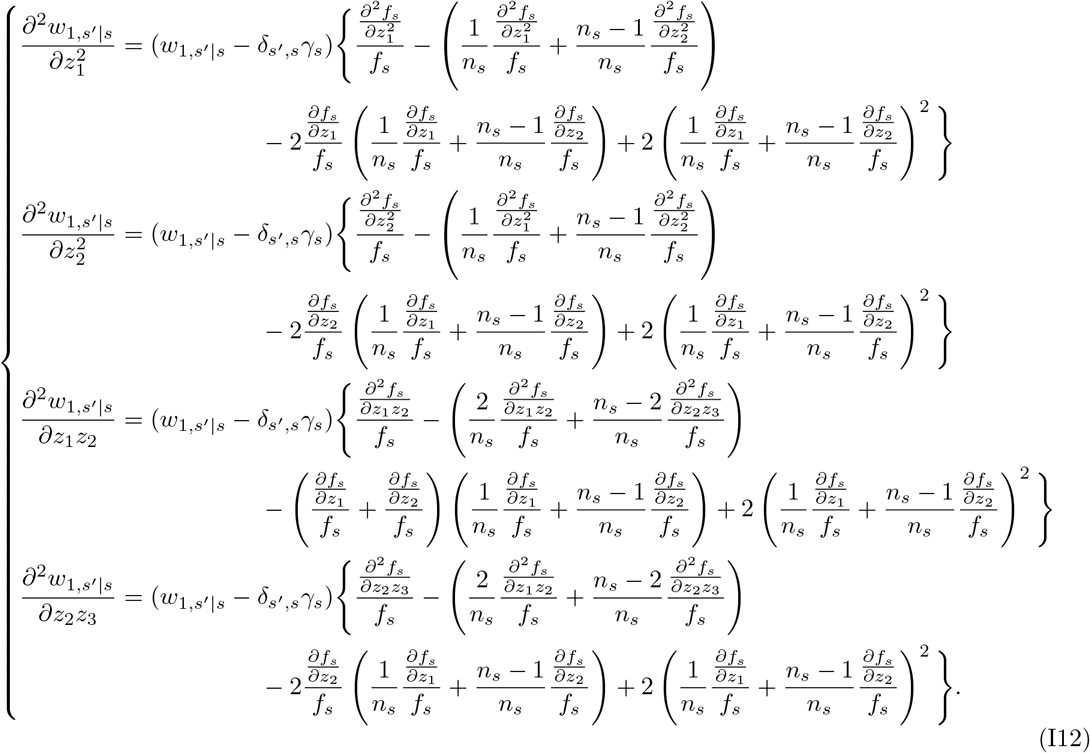

Now we are ready for calculating the second-order derivative of invasion fitness, given by eq.(34). We start from from *ρ*^(2w)^, given by eq.(34a). Substituting eq.(I12) in eq.(34) and using eq.(I7) produces the following four terms:

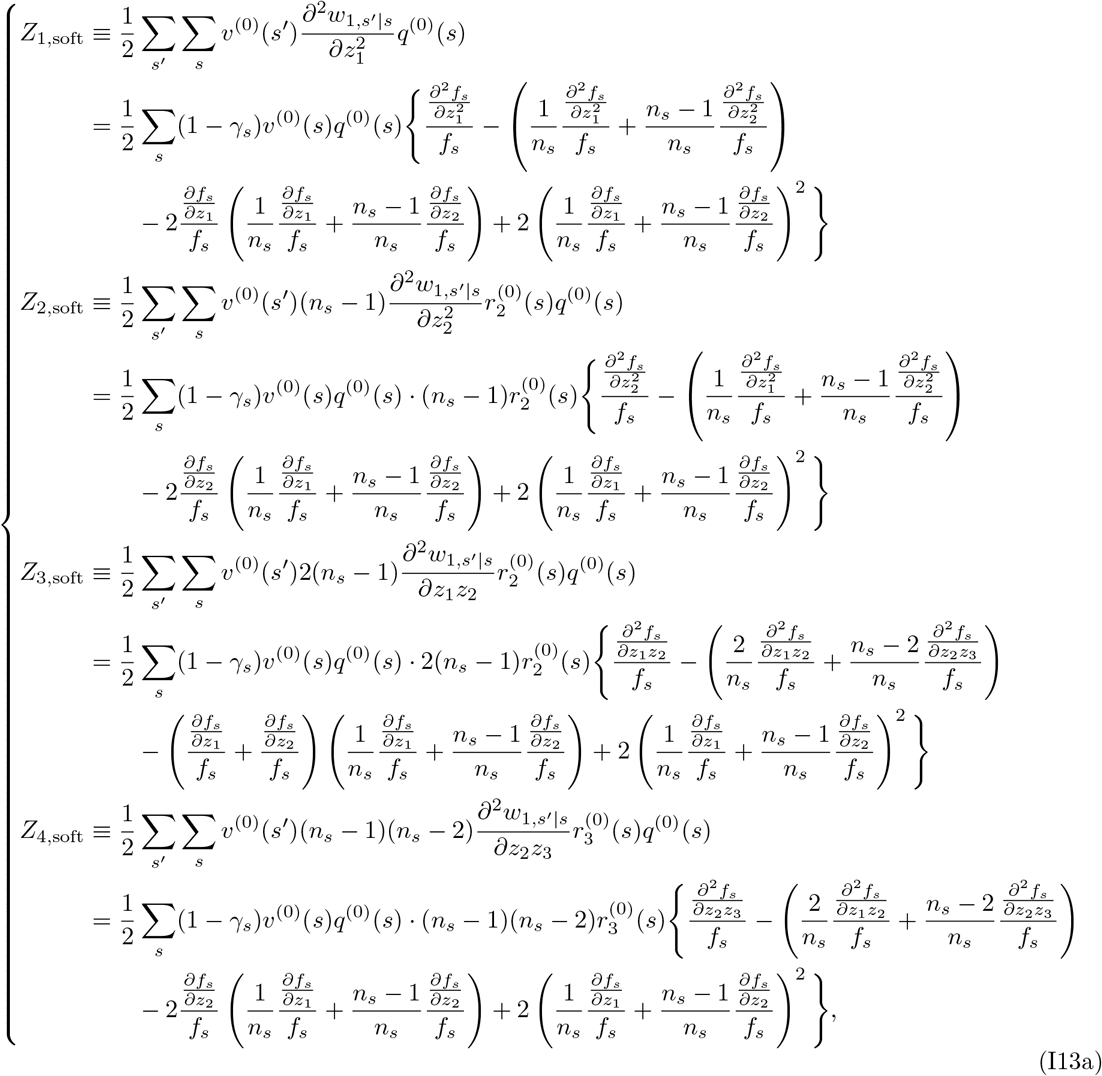

and we have

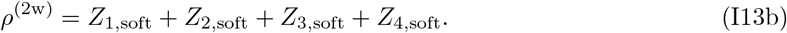

Here, 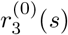 is calculated by substituting eq.(I2) in eq.(F32).

Next, we calculate *ρ*^(2q)^, given by eq.(34b). We repeat the same calculations as eqs.(I8) and (I9), but *q*^(0)^ there should be replaced with *q*^(1)^. From eq.(I10) we have

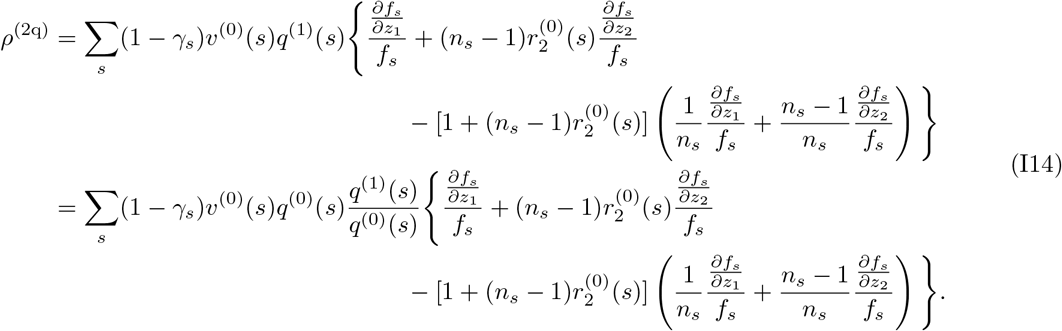

By substituting eqs.(I2) and (I5) in eq.(43) we obtain *q*^(1)^(*s*)*/q*^(0)^(*s*) as

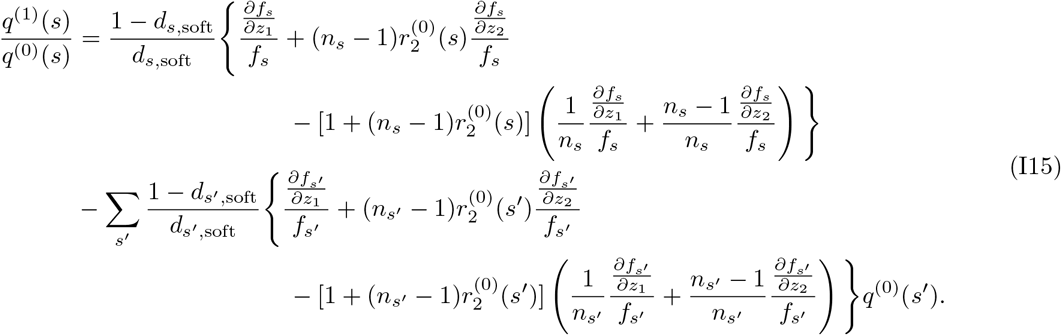

Now we substitute eq.(I15) into eq.(I14). Note that the second term in eq.(I15), which consists of the sum over *s′*, is just a constant, and that putting a constant in the place of *q*^(1)^(*s*)*/q*^(0)^(*s*) in eq.(I14) gives *ρ*^(1)^ times this constant (see eq.(I10)), which is zero by the assumption in this subsection. Thus we can substitute only the first term of eq.(I15) into eq.(I14) to obtain *ρ*^(2q)^, by which we obtain

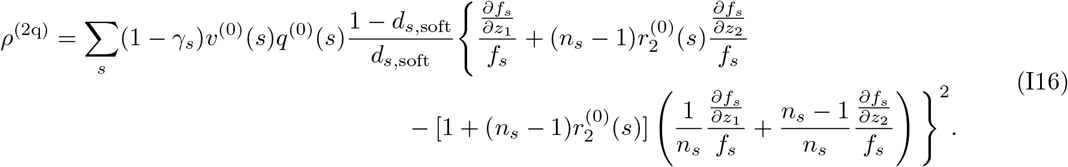

Quite notably, *ρ*^(2q)^ in eq.(I16) is always non-negative, which is apparently seen from its expression.

Thirdly, we calculate *ρ*^(2r)^, given by eq.(34c). For this, we repeat the calculation in eq.(I9), but 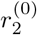 there should be replaced by 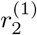. The result is

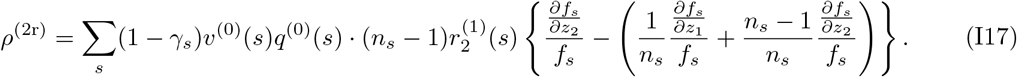

Here, we obtain 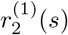 by substituting eqs.(I2) and (I5) in eq.(44), as

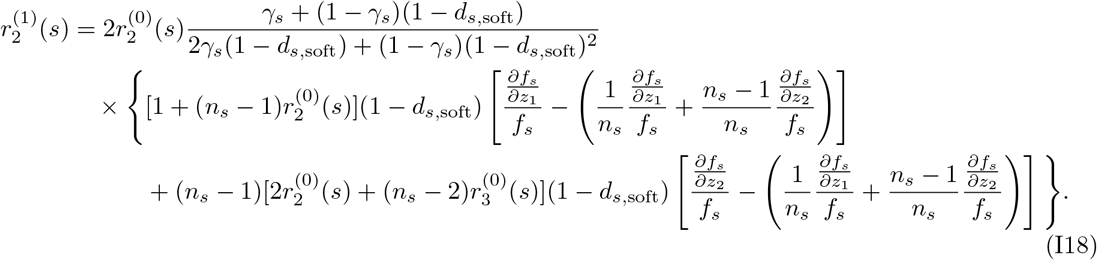

#### I.3 Consistency with previous results

Here, we again show that we recover previous results.

##### I.3.1 A model with “Regulation before dispersal” by Lehmann and Rousset (2010)

When there is a single habitat type (*N* = 1), no mortality in migration (*p_s_* = 1), and no overlap of generations (*γ_s_* = 0), eq.(62) reduces to

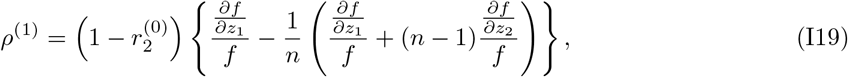

which reproduces eq.(A.7) of Lehmann and Rousset (2010) for their “regulation before dispersal” model with the following correspondence; 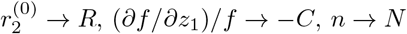, and 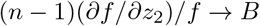.

##### I.3.2 A model with local adaptation by Svardal et al. (2015)

Svardal et al. (2015) studied a soft selection model with spatial and temporal environmental heterogeneity and effectively infinitely large group size. In the absence of temporal heterogeneity, their model fits our soft selection framework by setting *π_s_* = *c_s_*, *n_s_* = *n*_0_(*→ ∞*), *p_s_* = 1, *m_s_* = *m*, *γ_s_* = *γ* and *f_s_*(*z*_1_, ***z**_−{_*_1}_, *z*) = *f_s_*(*z*_1_). Then our eq.(62) predicts

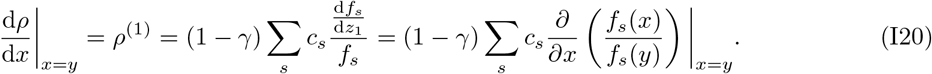

In the notation of Svardal et al. (2015) this can be written as (1 *− γ*)E_s_[*∂ρ*], which equals their first-order derivative of invasion fitness (see their Appendix B.1). Similarly, when the first-order derivative is null, eqs.(I13), (I16), and (I17) predict

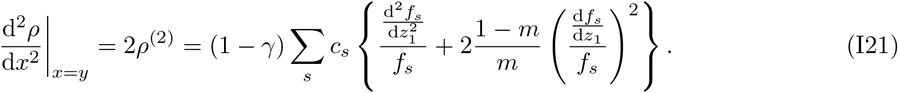

In the notation of Svardal et al. (2015) this can be written as (1 *− γ*){E_S_[*∂*^2^*ρ*] + {2(1 *− m*)*/m*}E_S_[(*∂ρ*)^2^]}, which agrees with their second-order derivative of invasion fitness, because in the absence of temporal heterogeneity their eq.(B.27) becomes

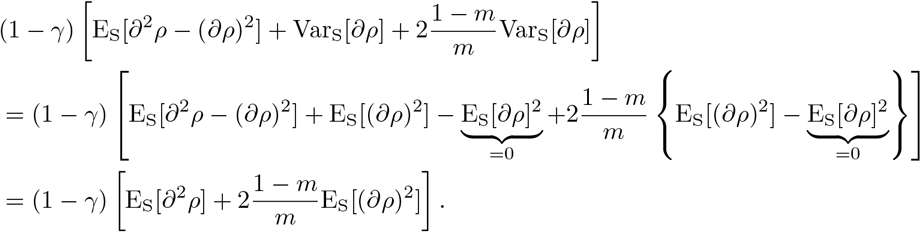

### J Evolutionary analysis of a lottery model with local adaptation

In this section, we derive the results for the special case presented in section 4.3 of the main text based on the assumptions that (i) the demography follows the assumptions in section 4.2 (either hard or soft selection), (ii) the Moran process limit is considered (*γ_s_* = *γ ∼* 1 for all *s*), and (iii) fecundity of an adult only depends on its trait value, as *f_s_*(*z*_1_, ***z**_−{_*_1}_, z) = *f_s_*(*z*_1_).

#### J.1 Hard selection

##### Analysis under assumptions (i) to (iii)

Under these assumptions, eq.(48) simplifies to

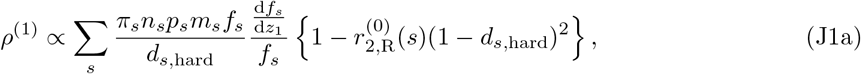

where the proportionality constant is

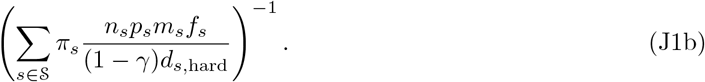

Remember that in eq.(J1), *ds,*hard, *fs* and its derivative, and 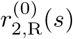 all depend on *y*. A singular strategy *y\** is the one at which *ρ*^(1)^ vanishes.

Next we study the second order effect of selection at the singular point *y\** by using eqs.(H13), (H16) and (H17). Note that our *f_s_* depends only on its bearer’s trait value, so most of the derivatives of *f_s_* that appear there are null. We start from eq.(H13):

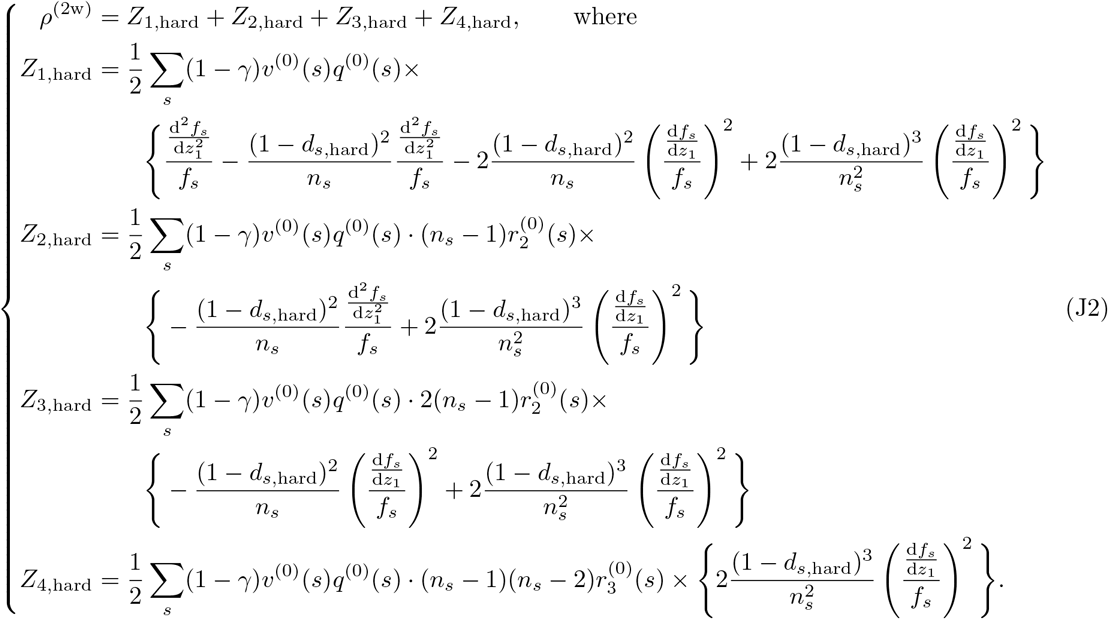

Next we calculate eq.(H16):

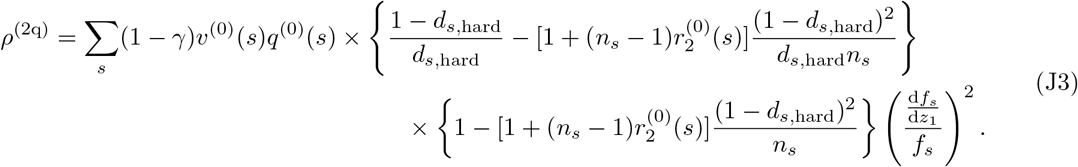

Then, we calculate eq.(H17):

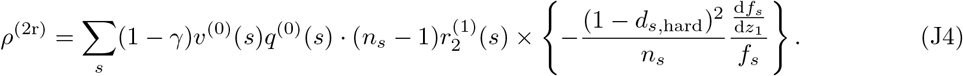

Relatedness values are calculated from eqs.(51), (42) (with 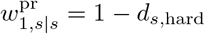 see eq.(H2)), and (H18) respectively, as

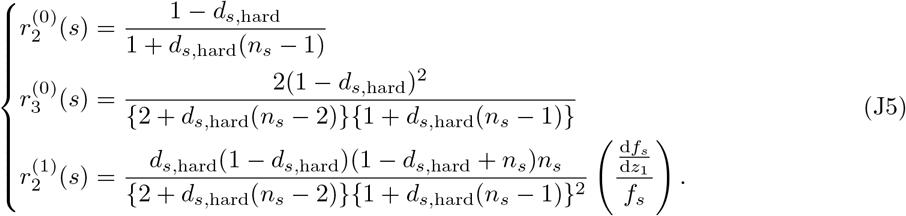

By using eqs.(J2)–(J5) and eq.(H3), and after doing some algebra, we arrive at the expression eq.(52) in the main text.

###### With two additional assumptions

Here, we add two more assumptions to make our model fully tractable: (iv) group size *n_s_*, the probability to migrate *m_s_*, and survival of migrating individuals *p_s_* are identical across habitats (*n_s_* = *n*, *m_s_* = *m*, and *p_s_* = *p* for all *s*), and (v) fecundity in groups in habitat *s* is given by eq.(53) in the main text.

For the selection gradient, by using eqs.(50), (53) and (J5), we calculate eq.(J1) as

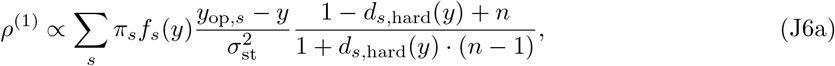

where the proportionality constant is

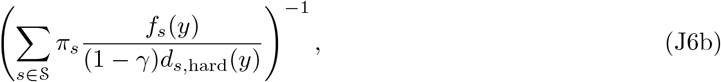

and the dependence of *f_s_* and *d_s,_*_hard_ on *y* is made explicit. The singular strategy *y\** is only implicitly solved as

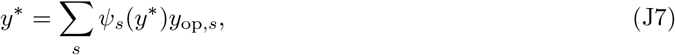

which is a weighted average of *y*_op,*s*_, the optimal trait values in groups in state *s*, with the weights *ψ_s_* given by

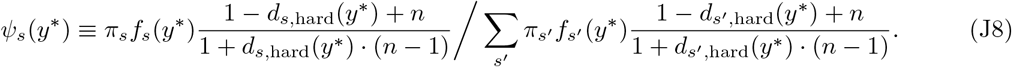

For the disruptive selection coefficient at the singular strategy, *y\**, a direct calculation of derivatives of eq.(53) in eq.(52a) leads to

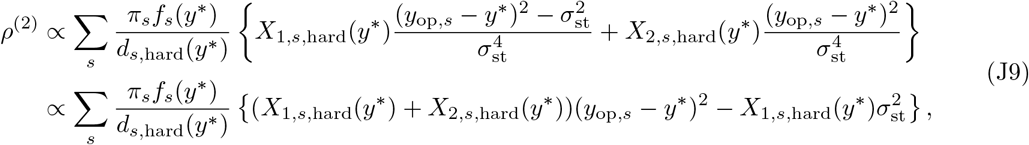

so the condition for *ρ*^(2)^ *>* 0 can be expressed as

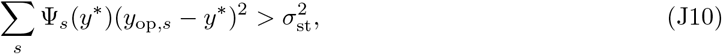

where

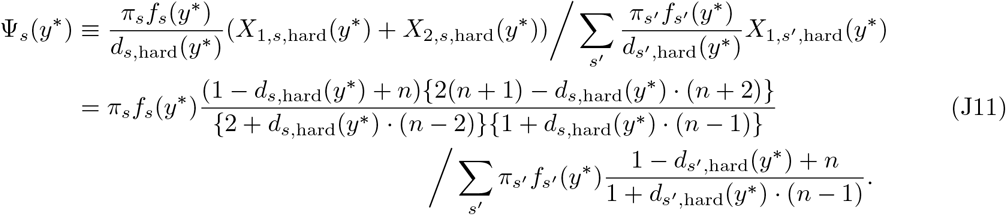

###### Further assumptions

Here we make three more assumptions: (vi) no mortality during dispersal, *p* = 1, (vii) two habitats with equal proportions, 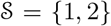 and *π*_1_ = *π*_2_ = 1/2, and (viii) optima are symmetric in the sense of *y*_op,2_ = *−y*_op,1_.

In this case *y\** = 0 is a singular strategy due to symmetry. To see this, from the symmetry of the fecundity functions *f*_1_(*y*) and *f*_2_(*y*), we have *f*_1_(0) = *f*_2_(0) (see eq.(53)). Then, from the definition of *d_s,_*_hard_ (eq.(49); see also eq.(47)), we have

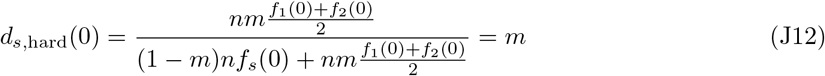

for *s* = 1, 2. Substituting these into eq.(J8) gives us *ψ*_1_(0) = *ψ*_2_(0) = 1/2, and hence eq.(J7) is satisfied.

We are interested under what conditions *ρ*^(2)^ *>* 0 holds. By using *π*_1_ = *π*_2_ = 1/2, *f*_1_(0) = *f*_2_(0) and eq.(J12), the weight given by eq.(J11) can be written as

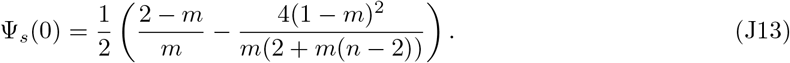

Therefore, when we define the variance of the habitat optima 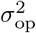 by eq.(57), the condition (J10) can be re-written as eq.(58) in the main text (note that we use *π*_1_ = *π*_2_ = 1/2 there).

Finally, we determine convergence stability of *y\** = 0. Calculating eq.(19) for the selection gradient eq.(J6) gives us, after some algebra,

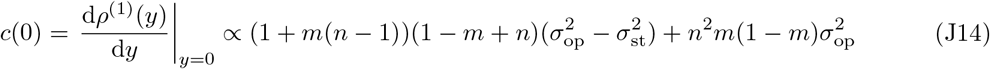

with a proportionality constant that is positive for *m >* 0. Rearranging the convergence stability condition, *c*(0) *<* 0, gives condition (60) in the main text. To obtain a sufficient condition for convergence stability, we substitute eq.(61) into the coefficient of 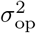 in (60) and obtain

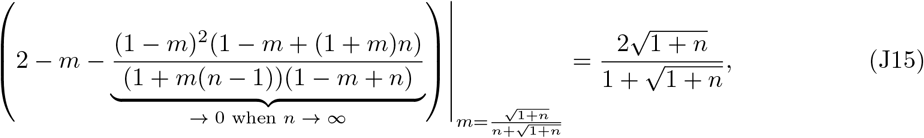

and therefore

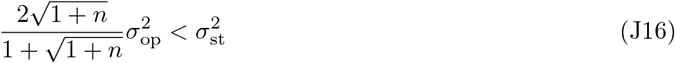

is a sufficient condition for convergence stability for fixed *n*. In addition, since eq.(J15) is upper-bounded by two, 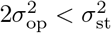 is a sufficient condition for convergence stability for any *n*.

##### J.2 Soft selection

###### Analysis under assumptions (i) to (iii)

Under these assumptions, eq.(62) simplifies to

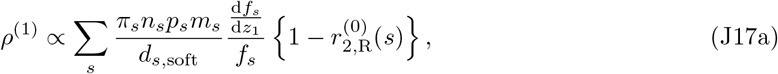

where the proportionality constant is

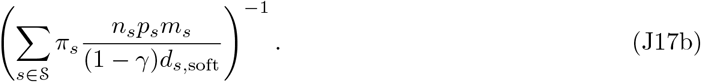

Remember that in eq.(J17), *f_s_*, its derivative, and 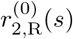 depend on the resident strategy *y*, whereas *d_s,_*_soft_ is independent of *y* (see eqs.(47) and (63)). A singular strategy *y\** is the one at which *ρ*^(1)^ vanishes. The disruptive selection coefficient at the singular point *y\** is calculated through eqs.(I13), (I16) and (I17) under the assumption that *f_s_* depends only on its bearer’s trait value, so most of the derivatives of *f_s_* that appear in *ρ*^(2)^ are null. In particular, from eq.(I13), we have

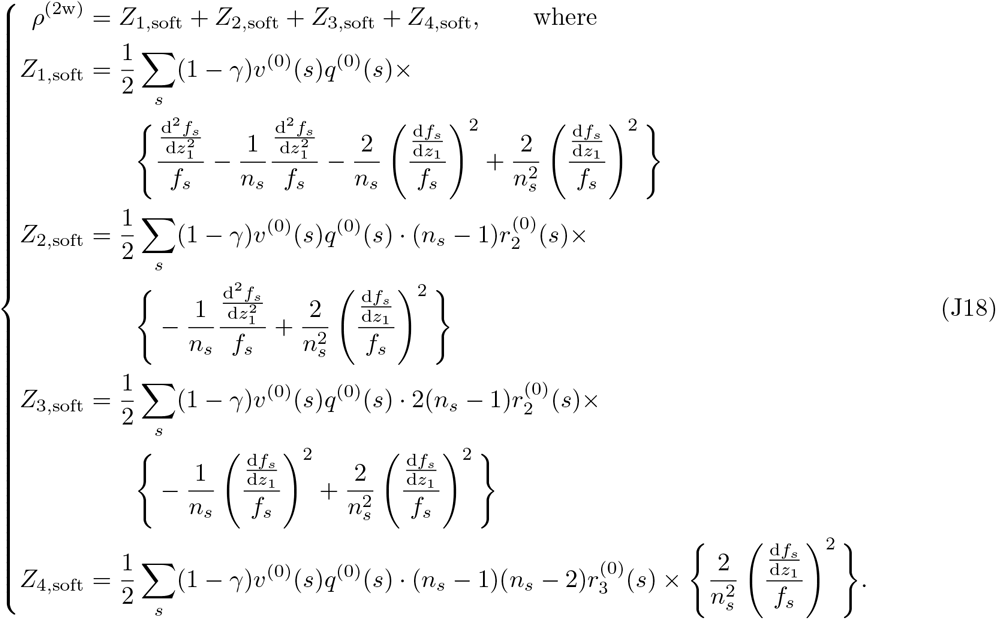

Second, eq.(I16) becomes

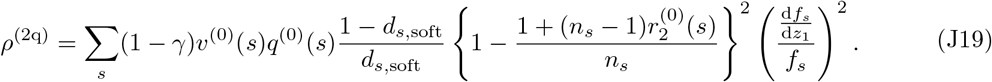

Third, eq.(I17) becomes

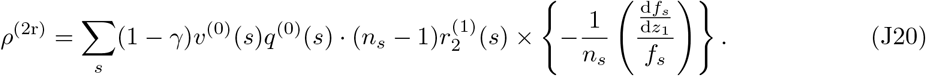

Relatedness values are calculated in the following way: 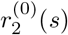 is from eq.(51) (where all *d_s,_*_hard_ are replaced by *d_s,_*_soft_; see section 4.3.2 in the main text), 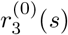 is from eq.(42) (with 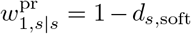 see eq.(I2)), and 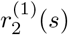 is from eq.(I18) combined with 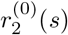 and 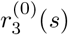 as just derived. We then obtain

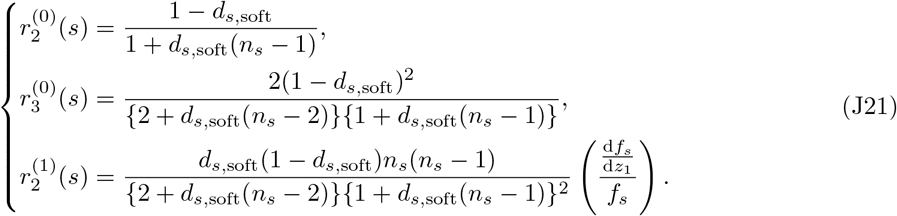

By using eqs.(J18)–(J21) and eq.(I3), and after some algebra, we arrive at the expression eq.(64) in the main text.

###### With two additional assumptions

Similarly to the hard selection case, we here add two more assumptions: (iv) group size *n_s_*, the probability to migrate *m_s_*, and survival of migrating individuals *p_s_* are identical across habitats (*n_s_* = *n*, *m_s_* = *m*, and *p_s_* = *p* for all *s*), and (v) fecundity in groups in habitat *s* is explicitly given by eq.(53) in the main text.

With these assumptions, from eqs.(47) and (63) we see that

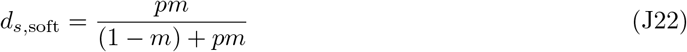

does not depend on *s*, and hence we write it as *d*_soft_ from now on. An immediate consequence is that *X*_1,*s,*soft_ and *X*_2,*s,*soft_ in eqs.(64b) and (64c) are independent of *s*, and we write them as

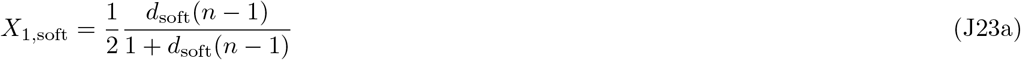

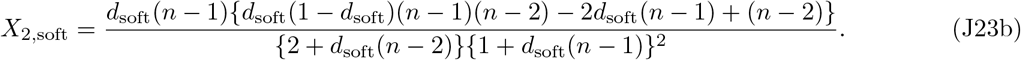

For the selection gradient, by using eqs.(50), (53) and (J21), we calculate eq.(J17) as

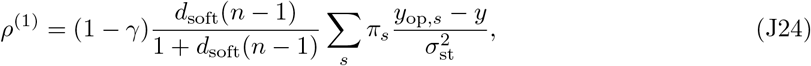

and therefore, the singular strategy is simply given by

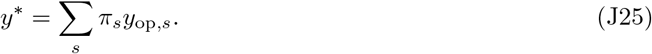

For convergence stability, we take the derivative of eq.(J24) with respect to *y* and obtain

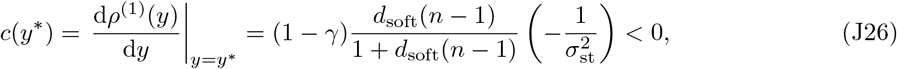

which shows that a singular strategy is always convergence stable. This parallels the results of Svardal et al. (2015).

For disruptive selection coefficient at the singular strategy *y\** a direct calculation of eq.(53) in eq.(64a) leads to

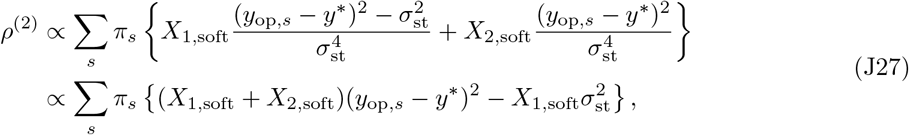

and the condition for *ρ*^(2)^ *>* 0 becomes

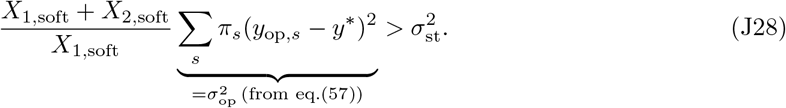

###### No mortality in dispersal

If we further assume that (vi) no mortality occurs during dispersal, *p* = 1, then substituting *p* = 1 into eq.(J22) gives *d*_soft_ = *m*. Inserting this into eq.(J23) and calculating condition (J28) with the expressions obtained for *X*_1,soft_ and *X*_2,soft_ results, after rearranging terms, in condition (66) in the main text.

## Footnotes

1 It is important to note that the conditioning in *w*(*s′|s, i*) is only on the state of the parental generation (as emphasized by the notation) and that *w*(*s′|s, i*) depends on group transition probabilities in models in which the state *s* of a group can change in each generation. See eqs.(E.1–E.2) in Lehmann et al. (2016) as well as section G.2 in the Supplementary Material for more details.

2 This can be seen by noting that when the focal and target individual from the focal set leave a realized number of *A*_1_ and *A*_2_ offspring, respectively, in the same group of size *n_s_*, then this group contributes to the focal’s 2-fitness 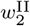 with *A*_1_ (the number of focal’s offspring) times *A*_2_/(*n_s_ −* 1) (the probability that a random neighbor of focal’s offspring is the target’s offspring), which equals to *A*_1_*A*_2_/(*n_s_* 1). Since *A*_1_*A*_2_/(*n_s_* 1) is symmetric with respect to *A*_1_ and *A*_2_, changing the roles of the focal and target individual does not alter the realized fitness count. The same logic applies when the focal and target individuals leave offspring to different groups, because in this case the counts per group are simply summed over all groups. A single individual’s 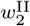 is the expectation of such counts over all realizations of offspring number in the same and different groups (where the expectation is taken over all single generation stochastic events affecting reproduction and survival), and the invariance holds because it holds for each realization.

3 For a homogeneous population with a single habitat *s*, a singular point is characterized by d*f_s_/*d*z*_1_ = 0, and therefore eq.(52) predicts that the sign of the disruptive selection coefficient is solely determined by the sign of 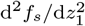 no matter whether dispersal is complete or locally limited. A similar result has been shown in Parvinen et al. (2017) by assuming a Wright-Fisher process.

1 We assume that this expectation exists for all *τ* and *τ′*. Using generating the function 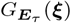, eq.(A4) is calculated as 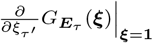, where **1** is a vector of ones.

